# Reversible biological motion with unidirectional catalysis through inversion of ATPase orientation

**DOI:** 10.64898/2026.02.24.707734

**Authors:** Gregory B. Whitfield, Ian Y. Yen, Lori L. Burrows, P. Lynne Howell, Yves V. Brun

## Abstract

Biological and engineered machines generally achieve reversibility through regulated switches in the directionality of fixed-orientation motors. Bacterial Tad pilus nanomachines extend and retract pili using a single motor with a unidirectional catalytic mechanism, a capability with no precedent. Using AlphaFold3 modeling, comparative structural analyses, and pilus activity assays, we find that the Tad motor ATPase CpaF achieves bidirectionality through physical inversion; alternating which face of the ATPase toroid engages the platform complex. The two orientations contact the pilus machinery in mutually exclusive extension- or retraction-specific configurations that drive reversible pilus dynamics using the same unidirectional catalytic cycle. Retraction requires conserved C-terminal residues in CpaF whose nucleotide-driven motions oppose those of the extension interface. Thus, nature has adopted a solution for reversible movement not yet conceived by human engineering.

## Main Text

Biological nanomachines are ubiquitous in nature and essential for life. These machines are powered by motors that convert chemical energy into directed mechanical motion through conformational changes (*1*). Many motor systems exhibit reversible or bidirectional activity, allowing for the dynamic regulation of processes such as intracellular transport, energy conversion, and motility. Some systems achieve reversibility by swapping between distinct sets of unidirectional motors with opposing activities (*2*), while others utilize positionally-fixed, bifunctional rotary motors that reverse activity through defined conformational transitions of key regulatory subunits (*3–5*).

The type IV filaments (TFFs) are a widespread superfamily of bacterial and archaeal surface appendages produced by evolutionarily related cell envelope-spanning nanomachines that facilitate diverse functions including motility, surface sensing and adhesion, protein secretion, and DNA uptake (*6–8*). To perform these functions, many TFFs must undergo dynamic cycles of extension and retraction, driven by motors composed of hexameric ATPases that engage a transmembrane platform protein complex to achieve filament polymerization and depolymerization (*9–13*). The largest and best-studied TFF group are the type IV pilus (T4P) systems, of which the type IVa pilus (T4aP) subgroup is the prototypical member. In T4aP systems, the opposing activities of pilus extension and retraction are mediated by interchangeable motor ATPases: PilB, which powers extension, and PilT, which powers retraction (Fig. 1A) (*14, 15*). Structural predictions suggest that these motor ATPases engage the platform complex through opposing hexameric faces (*16, 17*), implying that control of filament dynamics in these systems is achieved using opposing unidirectional motors combined with changes in ATPase orientation.

**Fig. 1:**
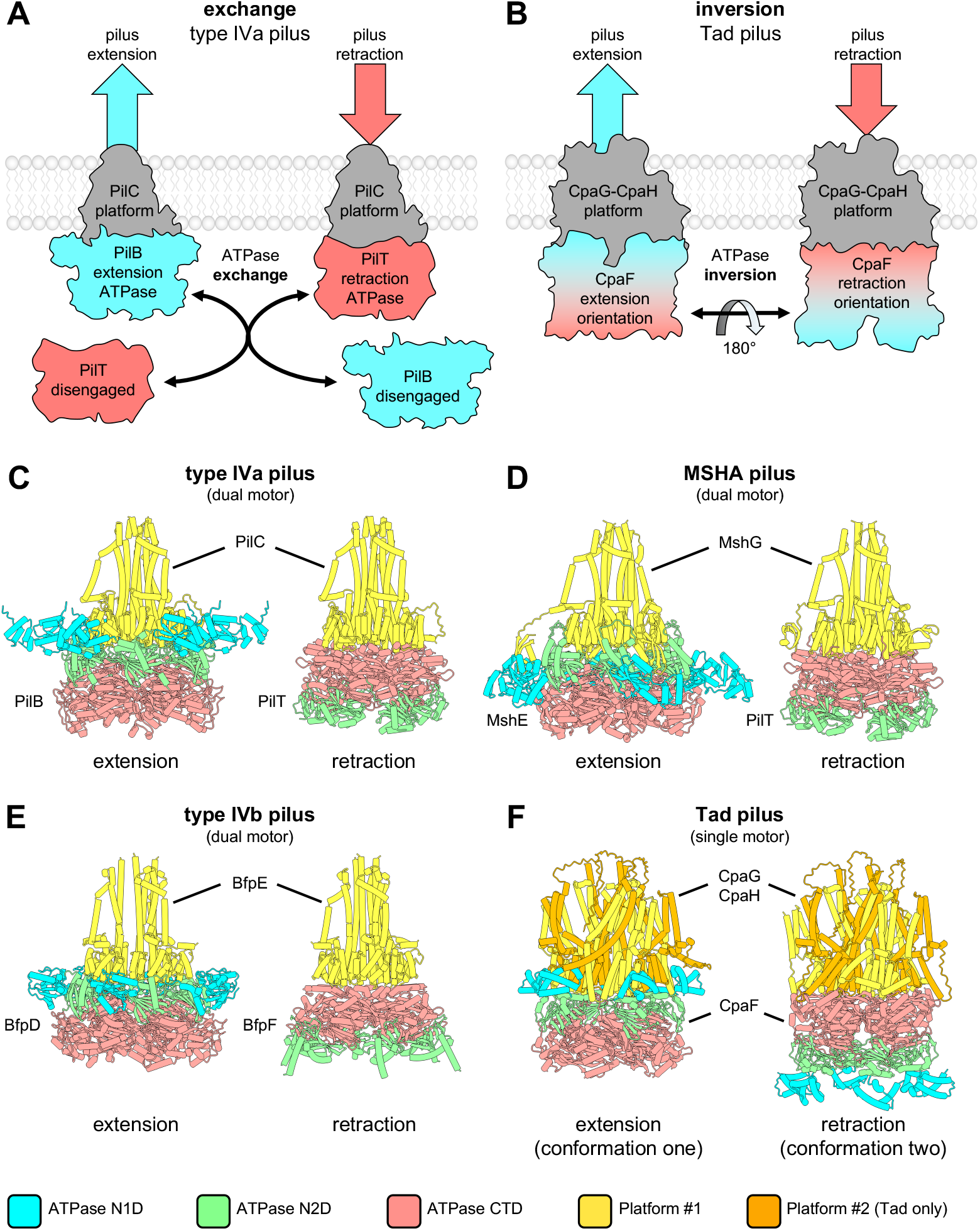
The Tad pilus motor is predicted to adopt two distinct conformations that mirror the extension- and retraction-specific motors of other pilus systems. (**A**) The type IVa pilus motor achieves reversibility by exchanging ATPases with opposing unidirectional activities. ATPase colour indicates their activity (blue, pilus extension; red, pilus retraction). (**B**) The Tad pilus motor achieves reversibility by inverting a single bifunctional ATPase with unidirectional catalytic activity relative to the platform proteins. (**C-E**) Structures of the extension- and retraction-specific motors of the T4aP from *P. aeruginosa* (panel C), the MSHA pilus from *V. cholerae* (panel D), and the T4bP bundle forming pilus from *E. coli* (panel E), predicted by AF3. (**F**) Structures of the two conformations of the Tad pilus motor from *C. crescentus* predicted by AF3. These conformations mirror those observed in pilus systems with dedicated extension and retraction ATPases (panels C-E) and so are named the extension- and retraction-specific conformations. N1D, variable N-terminal domain; N2D, conserved N-terminal domain; CTD, C-terminal domain.

By contrast, members of the widespread tight adherence (Tad) T4P subgroup employ a single motor ATPase, CpaF, to drive both extension and retraction of their pili (*18*). Initial structural characterization of CpaF provided strong mechanistic support for a unidirectional catalytic cycle driving pilus extension, and proposed a plausible allosteric means of achieving reversibility (*19*). However, while a subsequent investigation that identified a larger variety of CpaF conformational states further reinforced the extension-specific catalytic cycle, it was unable to corroborate the proposed reversibility mechanism (*17*). Therefore, there is no consensus on the mechanism of CpaF bifunctionality, with the preponderance of data from both studies supporting a non-reversible, unidirectional rotary mechanism of catalysis similar to extension ATPases like PilB. The simplest mechanistic solution to the ambiguity of a catalytically non-reversible ATPase driving reversible filament dynamics would require that CpaF engage the platform protein complex using one dedicated hexameric face for pilus extension, and the opposing hexameric face for retraction (Fig. 1B). Such a mechanism would mirror the opposing orientations through which PilB and PilT are predicted to interact with their cognate platform complex (Fig. 1C and Fig. S1A) (*16, 17*). If this hypothesis is correct, the Tad motor subcomplex (1) could be modelled in both of these conformations, (2) would have conserved residues on both hexameric faces of CpaF to facilitate these interactions, and (3) mutating conserved residues on one of these faces *in vivo* would disrupt the retraction-specific motor conformation, producing a retraction-deficient mutant.

To test this hypothesis, we began by generating models of multi-motor T4P subcomplexes using AlphaFold3 (AF3). Modeling of the T4aP extension (PilB) and retraction (PilT) specific motors indicated that the N-terminal face of the PilB hexamer (i.e., comprising the variable N1 and conserved N2 domains) engages with the cytoplasmically-exposed face of the PilC trimer, while the same interaction is mediated by the C-terminal domain (CTD) of the PilT hexamer (Fig. 1C and Fig. S1A). Similar configurations were observed for the extension- and retraction-specific motors of the mannose-sensitive hemagglutinin (MSHA) pilus (Fig. 1D and Fig. S1B) and the type IVb (T4bP) bundle forming pilus (Fig. 1E and Fig. S1C). These results are consistent with prior observations that platform engagement in dual motor T4P systems may occur using opposing ATPase orientations (i.e., N-versus C-terminal side of the hexamer) for extension versus retraction (*16, 17*).

When models of the single ATPase *Caulobacter crescentus* Tad motor subcomplex were generated, AF3 returned one of two distinct structures. Conformation one oriented the N-terminal face of the ATPase CpaF towards the CpaG-CpaH platform complex, while conformation two utilized the CTD of CpaF for this interaction (Fig. 1F). The AF3 prediction confidence metrics for these two models are nearly identical (Fig. S1D), implying that both are equally probable. Comparison with the multi-motor T4P systems suggested that conformation one most closely resembled extension motors while conformation two most closely resembled retraction motors (Fig. 1C-F). Thus, we refer to conformation one of the Tad motor subcomplex as the extension conformation and conformation two as the retraction conformation.

Next, we asked whether Tad motors from other bacterial species are also predicted to adopt these two orientations. AF3 structural modelling revealed that a selection of Tad motors from across the bacterial kingdom were predicted to adopt both extension and retraction conformations (Fig. S2) with comparable confidence metrics (Fig. S3, Table S1). However, we identified a subset of Tad motor orthologs that were predicted only in the extension conformation, despite attempts to favour the retraction conformation (Fig. S4, Table S1). Thus, we find supporting evidence for single motor T4P systems that associate via two differing conformations, with a likely subgroup of Tad motors that cannot adopt the retraction-specific conformation.

Next, we leveraged these modelling data to identify sequence features of CpaF orthologs unique to dual-orientation motors. We selected ten representative examples each of single- and dual-orientation ATPases (Table S1) for multiple sequence alignments. Core residues important for ATPase activity and platform protein interactions in the extension orientation (*17*) were generally in agreement between the two groups (Fig. S5). However, we noted significant differences in the C-terminus, corresponding to the platform protein interaction interface in the retraction conformation models. Three regions of interest were identified where single-orientation CpaF orthologs showed significant sequence loss and degeneration: (1) the N-terminus of α-helix 11 (α11 N-term), encompassing P386 and R387 (Fig. 2A, left), (2) β-strands 13 and 14 (β13-β14 turn) including several highly conserved residues (Fig. 2A, right), and (3) the C-terminal motif comprising β-strand 15, the flanking loops 26 and 27, and α-helices 13 and 14 (Fig. S6A). Mapping these regions onto the experimentally determined structure of CpaF (*17*) revealed that they account for much of the surface of the CpaF C-terminal face and that there are several conserved, solvent-exposed residues on this surface that could participate in interactions with the platform proteins (Fig. S6B). Particularly, in the retraction-orientation model generated by AF3, R387 is optimally positioned to contact both CpaG and CpaH (Fig. 2B). These results demonstrate that conserved residues, unique to dual-orientation ATPases, are clustered on the C-terminal face of the CpaF hexamer, where they are predicted to interact with the platform proteins in retraction-orientation models.

**Fig. 2:**
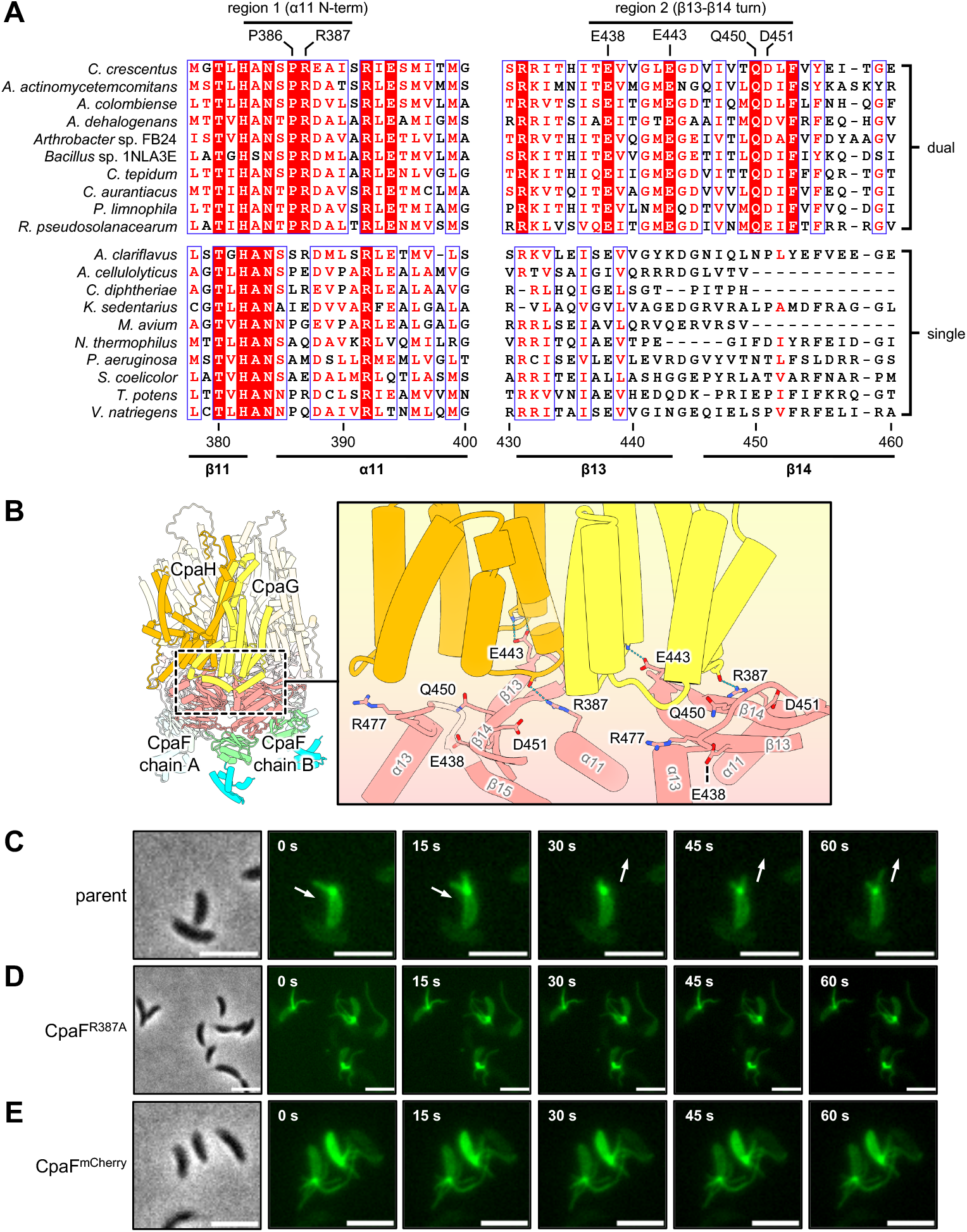
Conserved residues located on the C-terminal face of the CpaF hexamer are required for pilus retraction in *C. crescentus*. (**A**) Sequence alignment of ten representative CpaF orthologs predicted by AF3 to adopt extension and retraction orientations (dual, top) compared to ten representative CpaF orthologs predicted to adopt only the extension orientation (single, bottom). Positions with invariant residues are coloured white and highlighted with a red background, positions with ≥ 70% equivalent residues are coloured red. Regions of interest that are conserved in the dual-orientation alignment but not the single-orientation alignment are indicated at the top. Residue numbering and secondary structural elements are based on the *C. crescentus* sequence and structure. (**B**) The structure of the *C. crescentus* Tad motor predicted by AF3 in the retraction conformation, highlighting conserved residues on the C-terminal platform interaction interface that were targeted for mutagenesis. Predicted hydrogen bonding interactions are highlighted with blue pseudobonds. Proteins and protein domains are coloured as indicated in Figure 1. (**C-E**) Representative time-lapse images of the indicated strains labelled with AF488-maleimide (green). Maleimide spontaneously reacts with an engineered cysteine residue in the major pilin subunit, PilA. White arrows indicate the direction of movement (extension or retraction) of select pili in the parental example. A representative phase contrast image of each time-series is shown on the left. Scale bars, 3 μm. Parent = NA1000 *pilA*^T36C^.

If the ATPase inversion hypothesis for Tad motors is correct, mutation of these conserved residues in *C. crescentus* should destabilize the retraction orientation and produce a retraction-deficient mutant. To test this, we performed alanine mutagenesis on select residues from each of these regions in *C. crescentus* (Fig. 2B). As several of the mutations reduced CpaF protein levels (Fig. S7A), we overexpressed these variants to increase their abundance (Fig. S7B). Next, we directly examined the effect of these mutations on pilus biosynthesis and activity via fluorescence microscopy (*20, 21*). When compared to the parental strain (Fig. 2C and Movie S1), the R387A mutant produced pili that did not retract (Fig. 2D, Movie S2), displaying a phenotype comparable to cells that are unable to retract their pili due to chemical blocking (Fig. S7C, Movie S3) (*20, 22, 23*). The remaining mutants exhibited intermediate phenotypes characterized by pili that did not retract (Fig. S7D and Movies S4-S8) and pilus retraction events with frequent stalling (Fig. S8 and Movies S9-S13). However, since pili that do not retract were also occasionally observed in the parental background (Fig. S7D and Movie S14), we quantified the proportion of cells from each population that produced, but did not retract, a pilus (Fig. S9A). We also determined the number of non-retracting pili produced by each piliated cell (Fig. S9B), and their average length (Fig. S9C). These phenotypes were all significantly increased for the R387A mutant compared to the parent and were statistically comparable to populations in which pilus retraction was chemically blocked. Consistent with qualitative observations, many of the other CpaF mutants exhibited intermediate pilus phenotypes (Fig. S9A-C). However, unlike cells with chemically blocked pili, and despite the absence of observed pilus retraction in time courses, cells in the R387A mutant population exhibited fluorescent bodies comparable to the parental strain (Fig. S9D). This is indicative of externally labelled pilins that have been reincorporated into the inner membrane (*20, 21*), suggesting that pilus retraction may still occur in this mutant, albeit imperceptibly via fluorescence microscopy. Nevertheless, this data collectively supports the hypothesis that the C-terminal face of the CpaF hexamer is important specifically for pilus retraction activity.

Given these findings, we theorized that sterically blocking access to the C-terminal face of the CpaF hexamer should also lead to a retraction deficiency. To test this, we fused mCherry to the C-terminus of CpaF and examined the effect of this perturbation on pilus activity. In support of our hypothesis, no pilus retraction events were observed for this fusion in time courses (Fig. 2E and Movie S15), and quantified pilus phenotypes were comparable to, or elevated above, the R387A mutant (Fig. S9A-C). Furthermore, fluorescent cell bodies were present in the mCherry fusion population, but at a significantly reduced frequency compared to the R387A mutant (Fig. S9D), suggesting that physical obstruction of the C-terminal hexameric face of CpaF further reduces retraction activity. Collectively, these pilus assay results demonstrate that perturbations targeting the C-terminal face of the CpaF hexamer drastically impair pilus retraction phenotypes, providing functional support for the role of this surface in retraction-specific platform engagement (Fig. 1B).

Since residues located on the C-terminal face of CpaF are critical for pilus retraction, we next asked how these regions contribute to this activity. PilT-like retraction ATPases have a conserved motif (AIRNLIRE) that is required for pilus retraction but not ATPase activity (*24*). This motif is located on the C-terminal face of the PilT hexamer and is predicted to interface with the platform protein PilC (Fig. 1C and Fig. S10, top middle) (*17*). Alignment of PilT and CpaF places the β13-β14 turn motif of CpaF directly on top of the PilT AIRNLIRE motif (Fig. S10, top right). Similarly, CpaF has two conserved motifs on the extension face of the hexamer that are predicted to contact the platform proteins: α-helix 5 and the adjacent loop 8 (α5-loop 8 motif), and loop 17 that extends towards the centre of the hexamer (extended pore loop, Fig. S10, bottom middle) (*17*). Aligning the extension and retraction models positions the extended pore loop of CpaF directly coincident with the α11 N-term and β13-β14 turn motifs, and the α5-loop 8 motif adjacent to the β13-β14 turn (Fig. S10, bottom right). This suggests that there may be mechanistic parallels between the CpaF retraction motifs and the extension-specific elements previously mapped on the N-terminal face of CpaF, as well as the AIRNLIRE motif of PilT orthologs.

Structural determination of *C. crescentus* CpaF by cryo-EM resolved a catalytic mechanism that involves cycling between two conformations, termed ‘compact’ and ‘expanded’, as the axis of symmetry rotates in a clockwise direction about the center of the hexamer, as viewed from the extension face of the complex (Fig. S11) (*17*). We wondered whether there were conformational differences between the extension and retraction faces of the CpaF hexamer during nucleotide turnover and, if so, whether these differences were consistent with the opposing activities proposed in the ATPase inversion model. On the extension face, nucleotide turnover was linked to significant inter-chain movement in the hexamer with, for example, an approximately 19 Å expansion of opposing loop 8 motifs between the compact and expanded conformations (Fig. S11, top) (*17*). In comparison, the effects of nucleotide turnover on the retraction face of the CpaF hexamer led to similar but opposing changes: the axis of symmetry of the hexameric complex rotates in a counterclockwise direction between the compact and expanded conformations, with directly opposing residues of the β13-β14 turn motif contracting by up to 26 Å in the same dimension (Fig. S11, bottom).

Nucleotide turnover also led to significant intra-chain conformational changes. The structure of *C. crescentus* CpaF is C2 symmetric, therefore we focused our analysis on chains A, B, and C of the structure that form a trimeric complex comprising half of the full hexameric structure (*17*). In chain C, the extended pore loop moved towards the N-terminal face of the hexamer by approximately 12 Å between the compact and expanded conformations (Fig. 3A, red boxes). In the adjacent monomers, chain B acts as a hinge point, allowing chain A to translate laterally outwards from the centre of the hexamer, with α5 moving by approximately 12 Å (Fig. 3A, black boxes). Again, analysis of the effect of nucleotide turnover on intra-chain movements on the retraction face of the hexamer identified equivalent but opposing changes. In chain C, the retraction-specific motifs move away from the C-terminal face of the hexamer, with α11 N-term shifting by approximately 8 Å (Fig. 3B, red boxes). In chains A and B, chain A acts as the hinge point for the lateral translation of chain B inwards towards the centre of the hexamer, with the β13-β14 turn motif moving by approximately 8 Å (Fig. 3B, black boxes). These analyses collectively show that nucleotide turnover leads to both inter- and intra-chain conformational changes in the extension and retraction specific CpaF structural motifs that are diametrically opposed (Fig. 3C, Fig. S11). As these motifs are predicted to interface with the platform proteins, it follows that their opposing movements may disparately impact the conformation of the platform protein complex during nucleotide turnover, which may influence whether major pilins are incorporated into or removed from the pilus filament.

**Fig. 3:**
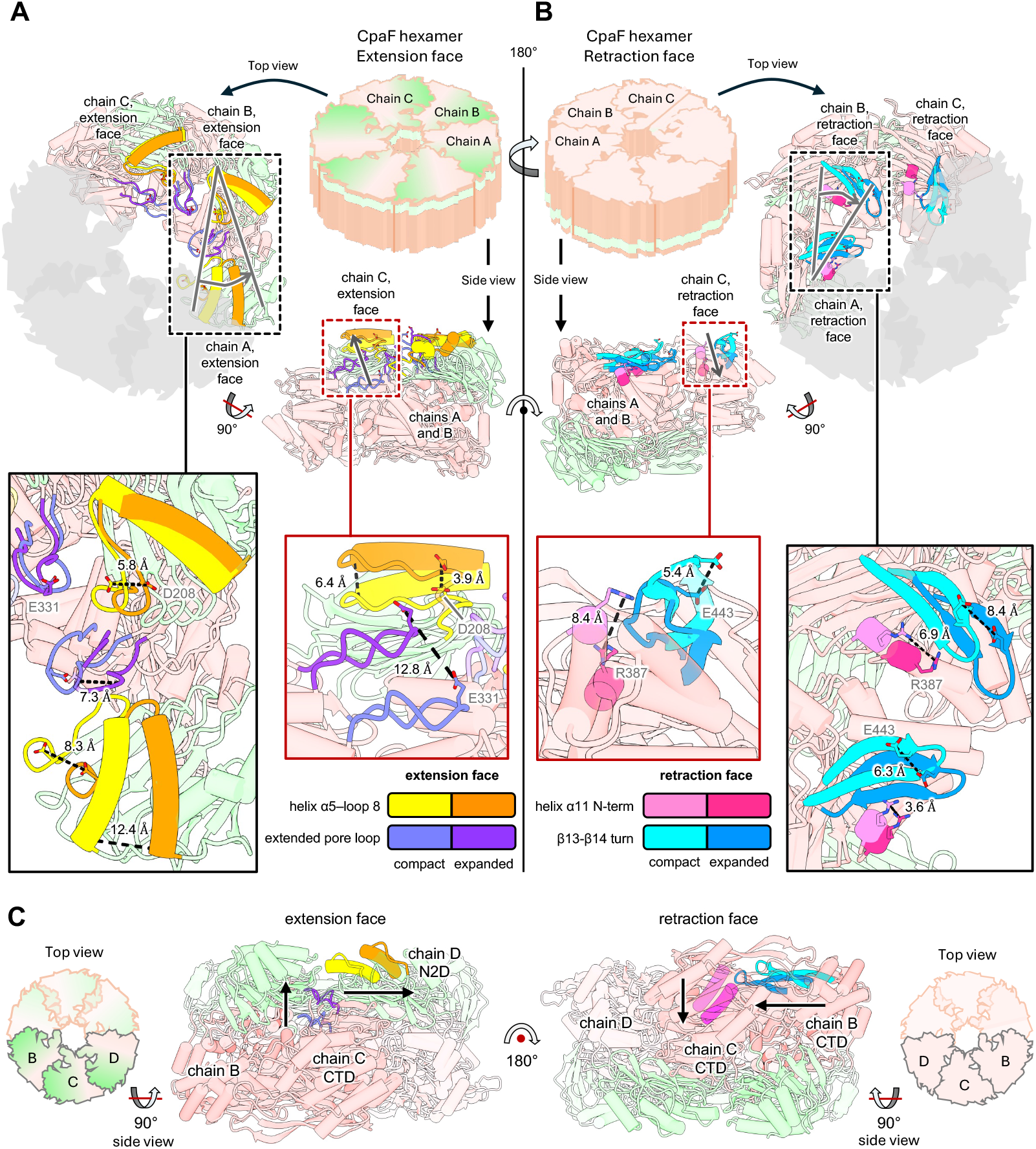
The retraction-specific structural motifs on the CpaF C-terminal face undergo conformational changes that oppose the extension-specific structural motifs on the N-terminal face. (**A**) Comparison of the extension-specific platform interaction motifs (legend at bottom) on the N-terminal face of the CpaF hexamer from the experimentally-determined compact (PDB 9E26) and expanded (PDB 9E29) conformations, which represent the changes that the CpaF hexamer undergoes as it turns over ATP. Due to the C2 symmetry of the CpaF hexamer, only a trimeric representation (chains A, B, and C) is depicted. In chains A and B, these motifs swing outwards from the centre of the hexamer, with chain B serving as a hinge point allowing chain A to undergo the largest lateral translation (black boxes). In chain C, these motifs move towards the extension face of the hexamer (red boxes). Distance measurements are indicated with black dashed lines. (**B**) Comparison of the retraction-specific platform interaction motifs (legend at bottom) on the C-terminal face of the CpaF hexamer. In chains A and B, these motifs swing inwards towards the centre of the hexamer, with chain A serving as a hinge point allowing chain B to undergo the largest lateral translation (black boxes). In chain C, these motifs move away from the retraction face of the hexamer (red boxes). CpaF domains are coloured as described in Figure 1. The N1 domains of CpaF have been omitted for clarity. (**C**) Summary of intra-chain conformational changes between the compact and expanded conformations of the CpaF hexamer. The extension-specific structural motifs (left) collectively move towards the extension face of the hexamer and away from the toroid centre, while the retraction-specific structural motifs (right) collectively move away from the retraction face of the hexamer and towards the toroid centre.

Having established ATPase inversion as a model to explain Tad motor bifunctionality, we next asked why one motor orientation may be adopted versus the other. As a starting point, we compared the extension and retraction orientation models for motor subcomplex orthologs from different species and examined which regions underwent the largest conformational changes. This revealed that helices at the periphery of the platform protein subcomplex underwent large predicted positional shifts, exceeding 40 Å in some examples (Fig. 4A and Fig. S12). Closer inspection of these models revealed that the helices undergoing the largest conformational changes were located at the N-termini of both CpaG and CpaH (Fig. 4B and Fig. S13). When the extension and retraction models were aligned via the platform subcomplexes, the N-terminal helices of CpaG and CpaH from the retraction models spatially overlapped with the N1 domain of CpaF from the extension models (Fig. 4B and Fig. S13). These observations collectively suggest that conformational changes in the N-terminal α-helices of the platform proteins could serve as a mechanism to promote an extension-to-retraction orientation switch through steric clashes with the N1 domain of CpaF.

**Figure 4:**
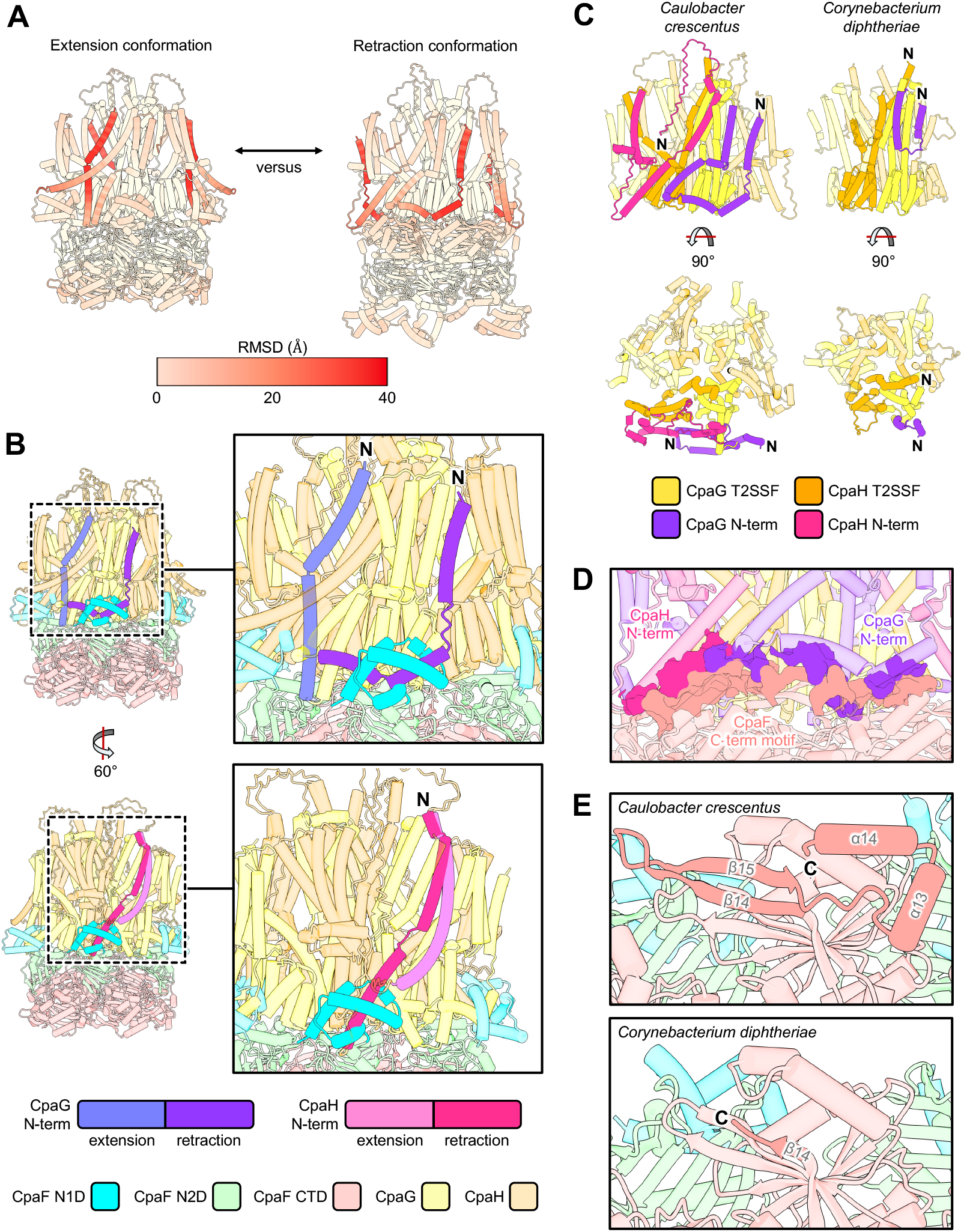
Platform protein N-termini are predicted to undergo conformational changes that could force ATPase reorientation through steric clashes. (**A**) Models of the extension (left) and retraction (right) conformations of the *C. crescentus* Tad motor were aligned and coloured according to the root mean square deviation (RMSD, legend at bottom) at each amino acid position. The platform protein complex and the CpaF hexameric complex were compared independently to exclude conformational deviations due to ATPase inversion. (**B**) Structural alignment of the *C. crescentus* Tad motor predicted by AF3 in the extension orientation with the platform protein complex from the retraction orientation model. Focus is on the conformational change of the CpaG (top) or CpaH (bottom) N-terminus between the extension and retraction conformations. In the retraction conformation, the platform N-termini bisect the CpaF N1 domain (cyan) from the extension orientation model. Protein and protein domain colouring is as indicated in the legend at the bottom. (**C**) Comparison of the N-terminal region adjacent to the conserved T2SSF domain of the platform proteins from a dual-orientation motor (*C. crescentus*, left) and a single-orientation motor (*C. diphtheriae*, right) as predicted by AF3. The N-terminal regions are coloured only for one of the three CpaG-CpaH heterodimers in the trimeric platform complex. (**D**) The interaction interface between the platform protein N-termini and the C-terminal motif of the ATPase, from the retraction-orientation prediction of the *C. crescentus* motor generated by AF3. Regions of interfacing residues are highlighted by the surface representation. Colouring is as indicated in panel B. (**E**) Comparison of the C-terminal motif of CpaF from a dual-orientation motor (*C. crescentus*, top) and a single-orientation motor (*C. diphtheriae*, bottom) as predicted by AF3.

If the above hypothesis is correct, then sequence losses in the platform protein N-termini would impact the ability of Tad motors to adopt the retraction-specific conformation. By analyzing CpaG and CpaH sequence alignments, we found that the N-terminal region adjacent to the conserved type II secretion system type F (T2SSF) domain of both proteins exhibited substantial differences (Fig. S14 and S15). In particular, we identified a 32% and 48% decrease in the average overall length of single-orientation CpaG and CpaH N-termini, respectively, compared to their dual-orientation counterparts (Fig. 4C, Fig. S16A-B, and Table S2). However, not all single-orientation motors exhibited significant sequence losses in their platform protein N-termini (Fig. S16 and Table S2). We observed that CpaG and CpaH N-termini were often predicted to interact with the CpaF C-terminal motif in retraction models (Fig. 4D and Fig. S17), and found that the C-terminal motif of CpaF orthologs undergoes significant sequence degeneration and loss in single-versus dual-orientation ATPases (Fig. S6A). Even in motors whose platform protein N-termini were intact, the C-terminal regions of their CpaF orthologs experienced sequence deterioration that impacted the predicted tertiary structure of this motif, which in dual-orientation motors was largely conserved (Fig. 4E, Fig. S18, and Table S2). These observations suggest that degeneration of the interaction between platform protein N-termini and the ATPase C-terminal region in single orientation motors may be related to the inability of these motors to adopt the retraction conformation. This further supports our hypothesis that the N-termini of the platform proteins may play a role in the mechanism of the extension-to-retraction switch, as cumulative sequence losses in this region could ultimately prevent the adoption of platform conformations that sterically clash with the N1 domain of the ATPase (Fig. 4B and Fig. S13).

Tad pili are part of the TFF superfamily of evolutionarily related, filament producing nanomachines in bacteria and archaea (*6, 7*). Like Tad pili, several members of this superfamily have only a single ATPase, despite evidence that their filaments can retract (*11–13, 21, 25*). We thus wondered whether ATPase inversion could explain how these systems are able to retract their filaments. To test this, we selected several representative systems from each of the single motor TFF family members (*6*) and predicted the structure of their motor subcomplexes with AF3. This analysis revealed that only motors from the archaeal EppA-dependent (Epd) T4P systems were predicted to adopt both extension and retraction conformations (Table S3). As bacterial Tad pili are thought to originate from horizontal transfer of the archaeal Epd system (*6*), these predictions support our prior assumption that single-orientation Tad motors underwent sequence degeneration and loss relative to ancestral dual-orientation motors. All other TFF motors were modelled only in the extension conformation. As a further test, we analyzed the conservation of surface residues on both faces of ATPase hexamers from representative TFF family members (Fig. S19-S22), as well as electrostatic compatibility between these faces and their cognate platform protein complex (Fig. S23-S25), which supported the outcome of the AF3 modelling (Table S4; see the corresponding Supplementary Results section). The asymmetries in potential interaction interfaces suggest that most single motor TFF systems, apart from the Tad and Epd pili, utilize ATPases that do not undergo inversion relative to the platform proteins to achieve activity reversals. Therefore, we propose that there are three types of motor arrangements in the TFF superfamily: the canonical class I motors that use interchangeable monofunctional ATPases, as found in the T4aP, class II motors that use a single non-inverting ATPase, as found in the T2SS and most T4bP and archaeal T4P systems, and class III motors that use a single inverting ATPase, as found in the Tad and Epd pilus systems (Fig. 5).

**Figure 5:**
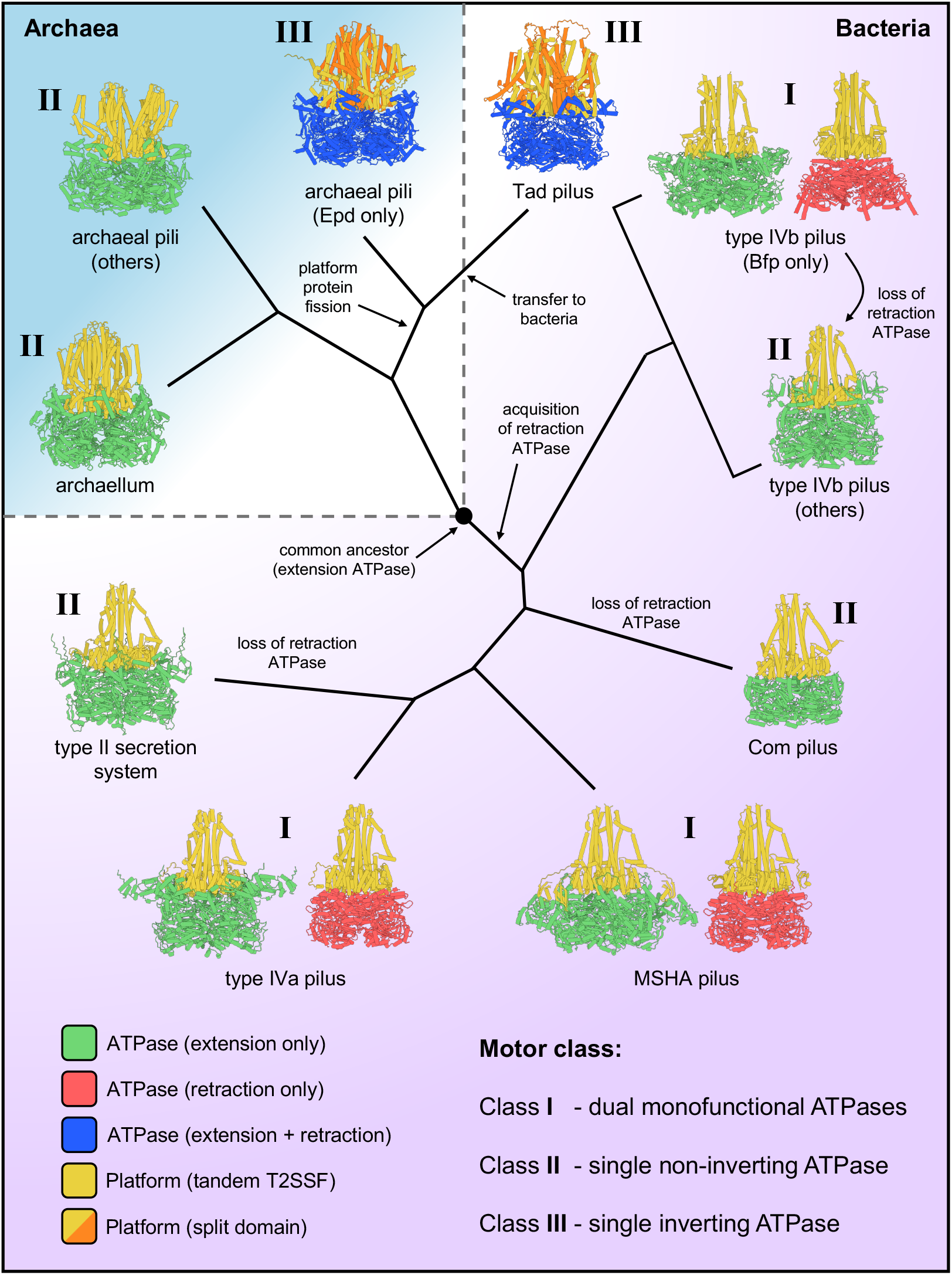
Prediction of motor functional mechanism is consistent with established type IV filament (TFF) system phylogeny. Representative tree, for illustrative purposes only, that captures the salient branching points and evolutionary events that define the TFF superfamily, adapted from Refs (*6, 7*). All ATPases in the TFF superfamily evolved from a common ancestor, which was likely an extension-specific ATPase (black circle, centre). The bacterial lineage acquired retraction ATPases early in their evolution, which led to the systems with dedicated extension and retraction ATPases (class I motor systems). In a subset of these systems, the retraction ATPase was lost, leading to systems with a single ATPase incapable of inversion (class II motor systems). The archaeal systems evolved from the same common ancestor, but no retraction ATPase was acquired in this lineage, thus class II motor systems are also found in archaea. Early in their evolution, however, a subset of these archaeal systems underwent fission of a tandem T2SSF domain-containing platform protein into two distinct polypeptide chains, which produced the Epd pilus that is capable of ATPase inversion (class III motor system). The Epd pilus was then transferred to bacteria, which evolved to become the Tad pili that are also capable of ATPase inversion. Branch lengths are not to scale.

The activity of biological nanomachines is dependent on the coordinated, directional activities of motor subunits (*1*). Reversibility is a key feature of many of these systems, which can be achieved through the regulated exchange of motors with opposing unidirectional activities. Such an arrangement explains the ability of T4P machineries, such as the T4aP, to drive dynamic cycles of pilus extension and retraction (*9*). In single-motor nanomachines, bifunctionality is achieved using rotary motors with reversible activities (*3–5*). However, these established paradigms fail to explain the bifunctionality of the single motor that powers Tad pilus extension and retraction, as multiple studies have largely supported the conclusion that CpaF utilizes a unidirectional rotary mechanism of catalysis (*17, 19*). This work identifies ATPase inversion as a previously unrecognized solution to achieve reversible motor activity in biological nanomachines using an enzyme that is catalytically non-reversible. By integrating previously documented, experimentally resolved CpaF structures (*17*) with the orientation-specific AF3 models presented herein (Fig. 1F), we show that extension- and retraction-associated motifs undergo diametrically opposed intra- and inter-subunit movements over the course of a catalytic cycle (Fig. 3 and Fig. S11). These counteracting motions likely influence the direction of force transmission imparted by the ATPase on the platform complex during filament polymerization or depolymerization. Thus, ATPase inversion not only switches the motor-platform interface but also reverses the consequences of nucleotide-driven conformational changes, without altering the catalytic cycle of the enzyme. This provides a plausible explanation for how a single unidirectional rotary ATPase (*17, 19*) can alternately promote pilin insertion or extraction from the filament base.

An unexpected yet consistent prediction from the models generated by AF3 is that the N-terminal helices of the platform proteins CpaG and CpaH undergo large positional shifts between extension and retraction models (Fig. S12). In retraction models, these helices intersect the space normally occupied by the CpaF N1 domain in extension models (Fig. 4B and Fig. S13), creating the potential for steric clashes that could favor inversion of the ATPase hexamer. Consistent with this hypothesis, Tad motors predicted to be incapable of inversion exhibit degeneration of the platform N-termini and the ATPase C-terminus (Fig. 4C-E, Fig. S16-S18, and Table S2). These correlations hint that platform remodeling may serve as the trigger for ATPase inversion. These observations may also explain the restriction of class III motors to the Tad and Epd pilus systems. The fission of a tandem T2SSF domain-containing platform protein into two distinct polypeptide chains is a unique aspect of the class III motor lineage (Fig. 5), and our analysis highlights that conformational flexibility of platform N-termini is a consistent feature distinguishing extension and retraction orientations (Fig. 4A and Fig. S12). Therefore, platform fission may serve to introduce the conformational flexibility necessary to induce ATPase inversion. In summary, our work indicates that the Tad ATPase CpaF alternately engages its cognate platform using distinct hexameric faces, and we define the structural and evolutionary determinants of this capacity, thus positioning the Tad system as the paradigm for a new class of bifunctional motors.

## Acknowledgements

We thank all members of the Brun lab for their valuable guidance and feedback as the project matured and the manuscript was prepared, Emma Mulholland, Sahar Alousi, Marie Dealby, Cecile Berne, Courtney K. Ellison, Shuaiqi Guo, Julien Herrou, and Sidney S. Shaw for critical reading of the manuscript, and Velocity Hughes (Synthesis by Velocity, Malmö, Sweden) and Andrew Jermy (Germinate Science Consulting) for editorial assistance. GBW was supported by a Natural Sciences and Engineering Research Council of Canada postdoctoral fellowship and a Fonds de Recherche du Québec – Nature et Technologies postdoctoral fellowship. IYY was supported by a Natural Sciences and Engineering Research Council of Canada PGS-D scholarship and a Hospital for Sick Children Research Institute Restracomp scholarship. This work was funded by project grant PJT-169053 from the Canadian Institutes of Health Research to LLB, PLH, and YVB. YVB is supported by the Canada 150 Research Chair in Bacterial Cell Biology.

## Author contributions

GBW performed structural predictions and analyses, generated chromosomal *C. crescentus* mutants, and performed microscopic imaging and quantification of pilus activity. IYY performed structural predictions and analyses and provided critical feedback that shaped the central hypothesis of the study. GBW wrote the manuscript and prepared figures. LLB, PLH, and YVB conceived, supervised, and coordinated the project. All authors contributed to the editing of the manuscript.

