## Supplementary materials for "Reversible biological motion with unidirectional catalysis through inversion of ATPase orientation"

**The PDF file includes:**

Materials and Methods
Supplementary Text
Figs. S1 to S25
Tables S1 to S6
References (*26*-*36*)

**Other Supplementary Materials for this manuscript include the following:**

Movies S1 to S15

### Materials and Methods

Bacterial strains, plasmids, and growth conditions

Bacterial strains, plasmids, and primers used in this study are listed in Table S5. *C. crescentus* strains were grown at 30 °C in peptone-yeast extract (PYE) medium (*26*). PYE was supplemented with 5 μg/mL kanamycin (Kan), where appropriate, for plasmid maintenance. Commercially available, chemically competent *Escherichia coli* DH5a (NEB5α, New England Biolabs) was used for plasmid construction and was grown at 37 °C in lysogeny broth (LB) supplemented with 25 μg/mL Kan, where appropriate, for plasmid maintenance.

Plasmids were transferred to *C. crescentus* by electroporation, as described previously (*27*). Chromosomal mutations were made by double homologous recombination using pNPTS138-derived plasmids, as previously described (*28*). Briefly, plasmids were introduced into *C. crescentus* by electroporation, then two-step recombination was performed using Kan resistance to select for single crossovers, followed by sucrose resistance to identify plasmid excision events. All mutants were validated by Sanger sequencing using primers targeting outside the region of recombination to confirm the presence of the mutation.

For construction of the pNPTS138-derived plasmids, ~300-600 bp regions of DNA flanking either side of the desired mutation were amplified from *C. crescentus* NA1000 genomic DNA. Upstream regions were amplified using upF and upR primers, downstream regions were amplified using downF and downR primers, and CpaF point mutations were incorporated into the 5' flanking regions of upR and downF primers (Table S5). upF and downR primers contained flanking sequences to facilitate insertion into pNPTS138 digested with EcoRV (New England Biolabs) by Gibson assembly (HiFi DNA Assembly Master Mix; New England Biolabs). Assembled plasmids were transformed into *E. coli* NEB5a (New England Biolabs), and clones with a positive insert were verified by Sanger sequencing.

For the plasmid pNPTS138::*cpaF*$^{\mathrm{mCherry}}$, ~ 500 bp regions upstream and downstream of the *cpaF* stop codon were amplified from *C. crescentus* NA1000 genomic DNA using the indicated upF + upR and downF + downR primers, respectively (Table S5), and the gene encoding mCherry was amplified from the vector pRVCHYN-2 (*29*) using midF and midR primers. Sequence encoding a linker (GSAGSAAGSGEF) between *cpaF* and mCherry was incorporated into the upR and midF primers. The mCherry variant encoded in pRVCHYN-2 contains a functional isoform due to an alternative start site, which can be eliminated by mutating methionine 10 to glutamine (M10Q) in the mCherry coding sequence (*30*). To do this, all three DNA products described above were used as template to amplify an upstream fragment using the *cpaF*$^{\mathrm{mCherry}}$ upF and mCherry$^{\mathrm{M10Q}}$ upR primers, and a downstream fragment using the mCherry$^{\mathrm{M10Q}}$ downF and *cpaF*$^{\mathrm{mCherry}}$ downR primers, with the M10Q point mutation incorporated into the 5' flanking regions of the mCherry$^{\mathrm{M10Q}}$ upR and downF primers (Table S5). These fragments were assembled into pNPTS138 as described above, the assembled plasmid was transformed into *E. coli* NEB5a (New England Biolabs), and clones with a positive insert were verified by Sanger sequencing.

CpaF overexpression constructs were made using pJC585, a reporter plasmid that contains RFP under the control of a taurine inducible promoter (*31*). Expression from this promoter is leaky, so it is not necessary to add taurine to growth medium for gene expression. *cpaF* genes with the desired mutation were generated by amplifying the upstream and downstream fragment of the *cpaF* open reading frame from *C. crescentus* NA1000 genomic DNA, centred on the desired mutation, using $P_{tau}$ upF + $P_{tau}$ upR and $P_{tau}$ downF + $P_{tau}$ downR primers (Table S5). CpaF point mutations were incorporated into the 5' flanking regions of $P_{tau}$ upR and $P_{tau}$ downF primers. $P_{tau}$

upF primers encoded a synthetic ribosome binding site (TTTAAGAAGGAGATATACAT). The upstream and downstream fragments were assembled by overlap extension PCR with $P_{tau}$ upF and $P_{tau}$ downR primers. The PCR products were digested with EcoRI and BamHI (New England Biolabs) and ligated into pJC585 digested with the same enzymes, which removes the vector-encoded *rfp* sequence. Assembled plasmids were transformed into *E. coli* NEB5a (New England Biolabs), and clones with a positive insert were verified by Sanger sequencing.

Pilus labeling, blocking, imaging, and quantification

The pili of *C. crescentus* were labelled as described previously (*21*). Briefly, 25 µg/mL of Alexa Fluor 488 $C_5$ Maleimide (AF488-mal, ThermoFisher Scientific) was added to 100 µL of early exponential phase *C. crescentus* cell culture ($OD_{600}$ = 0.1–0.3) and incubated for 5 min at room temperature. To artificially block pilus retraction, 500 µM of methoxy-polyethylene glycol maleimide with an average molecular weight of 5000 Da (PEG5000-mal, Sigma) was added to the *C. crescentus* cell culture immediately prior to the addition of 25 µg/mL of AF488-mal. Labelled and/or blocked cells were collected by centrifugation for 1 min at 5000 × *g* and washed once with 100 µL of PYE to remove excess dye. The cell pellets were resuspended in 30 µL of PYE, 1 µL of which was spotted onto a 1% agarose PYE pad (SeaKem LE, Lonza Bioscience). The agarose pad was sandwiched between glass coverslips for imaging, which was performed using a Nikon Ti2 inverted fluorescence microscope with a Plan Apo 60× objective, a green fluorescent protein (GFP) filter cube, a Prime BSI Express sCMOS camera, and Nikon NIS Elements imaging software. The percentage of cells within the population with non-retracting pili, the number of non-retracting pili produced per piliated cell, the average length of non-retracting pili, and the percentage of cells within the population with fluorescent cell bodies, were quantified manually using ImageJ software (version 1.54p; *32*).

Western blot analysis

To determine the amount of CpaF produced by different *C. crescentus* strains, approximately $10^9$ cells from early exponential phase cultures ($OD_{600}$ = 0.1–0.3) were collected by centrifugation for 5 min at 5000 × *g*. The supernatant was removed, and the cell pellets were resuspended in 100 µL of 4× SDS loading buffer (200 mM Tris-HCl pH 6.8, 40% (v/v) glycerol, 8% (w/v) SDS, 20% (v/v) β-mercaptoethanol, and 0.005% (w/v) bromophenol blue). Samples were boiled for 10 min and then separated by SDS-PAGE using 12% gels. The samples were transferred to nitrocellulose membranes (100 V, 1.5 h) and blocked in 5% (w/v) skim milk powder resuspended in Tris buffered saline with Tween-20 (TBS-T; 10 mM Tris-HCl pH 7.5, 150 mM NaCl, 0.05% (v/v) Tween-20) for 2–4 h. Membranes were probed with α-CpaF antibodies (Biomatik) (*18*) at 1:5000 dilution in 1% (w/v) skim milk powder resuspended in TBS-T for 16 to 20 h. Membranes were then washed four times with TBS-T and probed with horseradish peroxidase-conjugated goat anti-rabbit antibody (Pierce, #1858415) at 1:5000 dilution in 1% (w/v) skim milk powder resuspended in TBS-T for 1 h. Membranes were washed again four times with TBS-T, developed using SuperSignal West Pico Plus chemiluminescent substrate (ThermoFisher Scientific), and imaged on a Bio-Rad ChemiDoc MP imaging system. As a loading control, membranes were probed with α-GAPDH antibodies (Biomatik) (*17*) at 1:5000 dilution, as described above. The Western blot images shown are representative of three independent biological replicates.

Structural modeling and analysis

All TFF motor subcomplex predictions were generated using AlphaFold3 (AF3, https://alphafoldserver.com) (*33*) with a minimum of five predictions performed with random seeding. The identity of each protein used for these predictions is listed in Table S6. Six copies of the ATPase and three copies of the platform protein(s) were provided as input for each prediction. All structure manipulation, including colouring of predictions by their predicted local distance difference test (pLDDT) scores and the generation of predicted aligned error (PAE) plots, was performed using ChimeraX (version 1.11; *34*). Mapping of residue conservation onto the surface of protein structures was performed using Consurf (*35*). Surface electrostatics were calculated and mapped onto protein structures using the coulombic function of ChimeraX. Sequence alignments were generated using ESPript 3 (*36*).

### Supplementary Results

<u>Other single motor type IV filament machines are unlikely to undergo ATPase inversion</u>

Structural prediction of representative TFF motors by AF3 revealed that only the archaeal Epd and bacterial Tad pilus systems were predicted to adopt both extension and retraction conformations (Table S3). All other motors were modelled only in the extension conformation, regardless of whether the variable N1 domains of the ATPases were present or not. Although these results may indicate that these motors are incapable of adopting retraction orientations, there could also be unrecognized biases that favour extension conformation predictions, such as system-specific interaction partners that might stabilize the retraction orientation. We reasoned that, if this hypothesis is correct, both the N- and C-terminal surfaces of the hexameric ATPase complexes from these systems should have regions of conserved residues that mediate their interactions with the platform protein complex. If not, the absence of evolutionary pressure to maintain such an interaction interface would result in sequence degeneracy. As a proof-of-principle, we examined surface residue conservation of the extension and retraction specific ATPases from the T4aP, T4bP, MSHA, and Tad pilus systems (Figure 1C-F). This analysis revealed highly conserved patches of sequence on the N-terminal faces of the extension ATPases that correlated with the predicted platform interaction interface, while the C-terminal faces were more variable (Figure S19). Conversely, the N-terminal faces of the retraction-specific ATPases exhibited more sequence variability, while the C-terminal faces, with which the platform complex is predicted to interact, were more conserved. For the Tad pilus ATPase CpaF, both the N- and C-terminal faces of the hexamer exhibited comparable surface residue conservation to the extension- and retraction-specific ATPases, respectively (Figure S19), as expected for a motor that can engage with the platforms using either hexameric surface. We next selected a representative example from each of the remaining single-motor TFF systems (Table S3, Figures S20 and S21) for surface residue conservation analysis. The archaeal UV-inducible pilus system (Ups), T2SS, and Gram-positive Com pilus motors all exhibited surface residue conservation patterns comparable to the extension-specific ATPases (Figure S22, compare to Figure S19), further suggesting that these motors may only adopt the extension conformation. The Epd pilus motor had residue conservation patterns on both hexameric faces that were comparable to the Tad pilus motor (Figure S22, compare to Figure S19), consistent with predictions suggesting that it can adopt extension and retraction conformations (Figure S20). However, both the Tcp and archaellar ATPases also had significant residue conservation on both the N- and C-terminal faces of the hexamer (Figure S22), at odds with the outcome of the AF3 prediction analyses (Table S3).

Residue conservation analyses alone can only suggest that a surface is or is not experiencing selective pressure to retain an interaction partner, not what that interaction partner might be. Thus, we complemented this analysis with an examination of surface electrostatics to determine whether either face of the ATPase hexamer is compatible with platform protein interactions. Using the dual-motor T4P systems as a proof of principle, we found that surface electrostatics of the platform complexes was compatible with the N-terminal faces of the extension-specific ATPases, and largely incompatible with the C-terminal faces, and that this pattern was reversed for the retraction-specific ATPases (Figure S23). Furthermore, for the dual orientation Tad and Epd pilus motors, both N- and C-terminal faces of the ATPases were electrostatically complementary to the platform protein complex (Figure S24). These results are consistent with those obtained from the surface residue conservation analysis (Figures S19 and S22). For all the remaining single ATPase TFF systems, including the Tcp and archaellar motors, there were clear patterns of complementary electrostatics between the platform complex and the N-terminal faces of the ATPases, and obvious incompatibilities with the C-terminal hexameric faces (Figure S25). This finding further supports the outcomes of the AF3 predictions and suggests that surface residue conservation on the C-terminal faces of the Tcp and archaellar motors is likely coincidental. Collectively, these analyses, summarized in Table S4, suggest that most single motor TFF systems, apart from the Tad and Epd pili, utilize ATPases that do not undergo inversion relative to the platform proteins to achieve activity reversals.

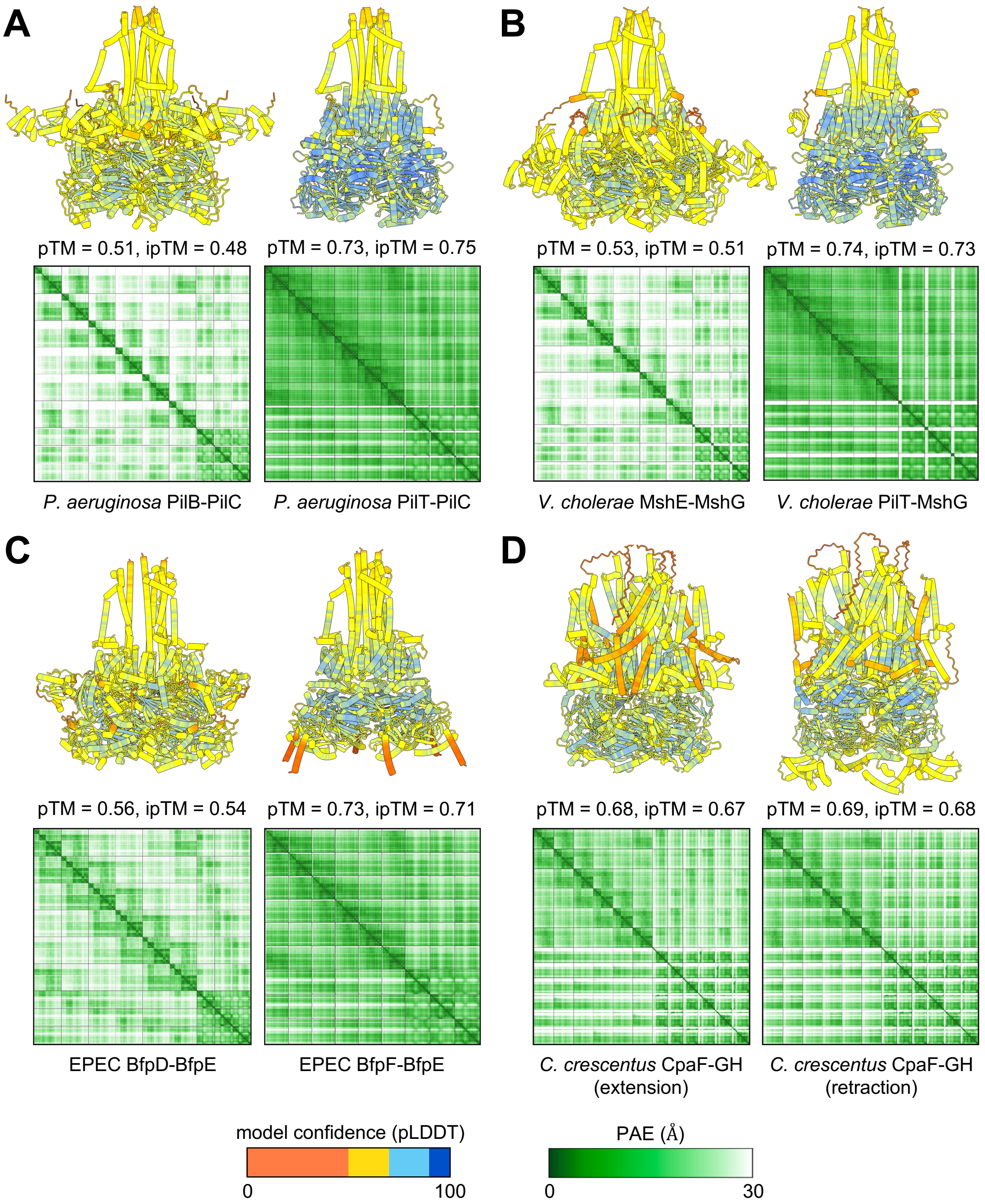


**Fig. S1: Confidence metrics generated by AF3 for the pilus motor models depicted in Figure 1, which are comparable for the two conformations of the Tad motor.** (**A**) The extension (left) and retraction (right) motors of the *Pseudomonas aeruginosa* type IVa pilus. (**B**) The extension (left) and retraction (right) motors of the *Vibrio cholerae* mannose sensitive hemagglutinin (MSHA) pilus. (**C**) The extension (left) and retraction (right) motors of the enteropathogenic *Escherichia coli* (EPEC) type IVb bundle forming pilus. (**D**) The extension (left) and retraction (right) conformations of the *Caulobacter crescentus* Tad motor. All predictions were performed a minimum of five times with random seeding; only the top scoring, top-ranked model is depicted.

Models are coloured according to the predicted local distance difference test (pLDDT) scores (legend at bottom left). Below each model, the corresponding predicted aligned error (PAE) plots are depicted (legend at bottom right). The predicted template modelling (pTM) and interface predicted template modelling (ipTM) scores are provided below each model.

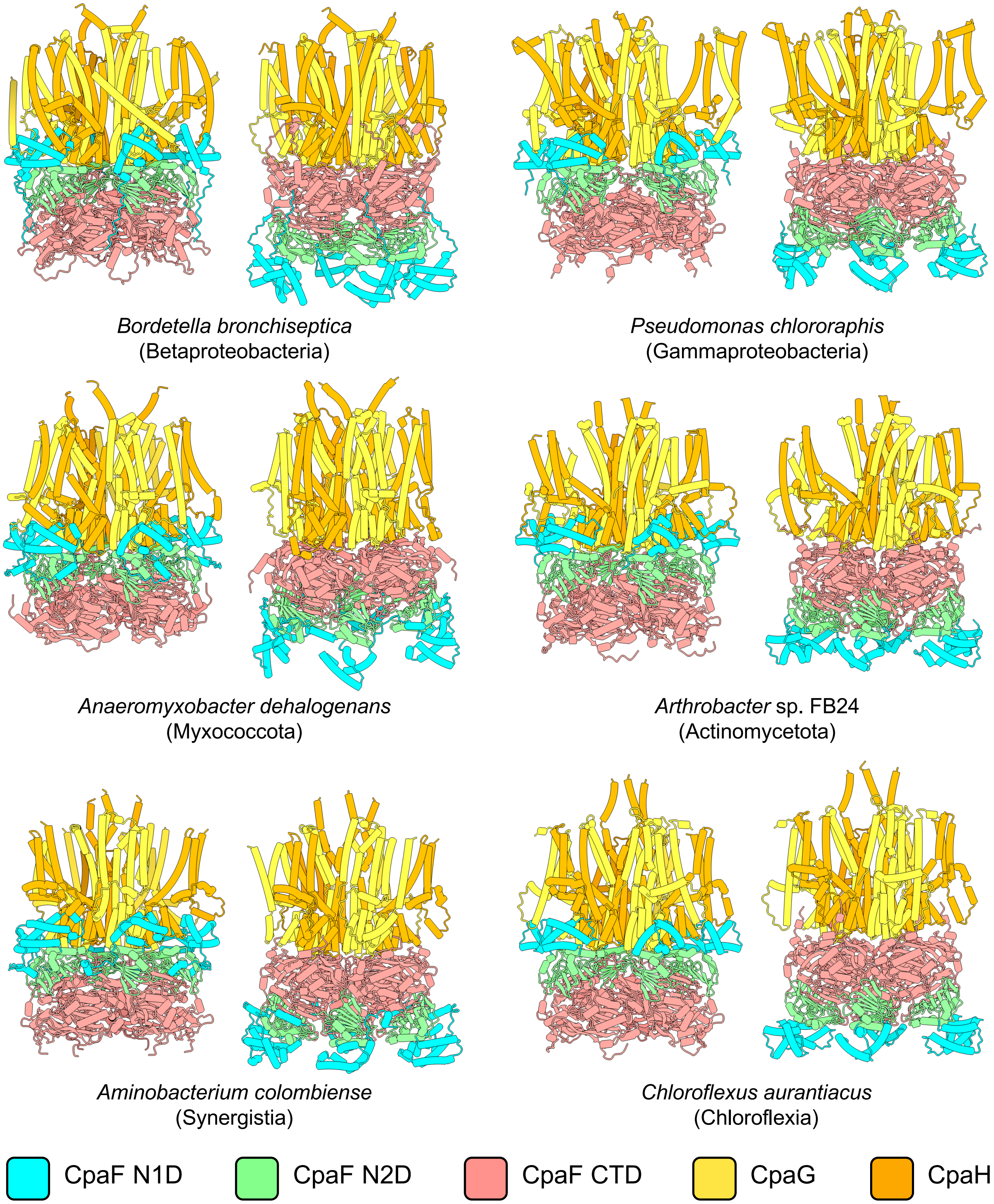


**Fig. S2: Tad motor orthologs from across bacterial taxa are predicted by AF3 to adopt extension- and retraction-specific conformations.** For each motor ortholog, the extension orientation is shown on the left and the retraction orientation is shown on the right. Colours correspond to specific proteins or protein domains, as indicated in the legend at the bottom. N1D, variable N-terminal domain; N2D, conserved N-terminal domain; CTD, C-terminal domain. All predictions were performed a minimum of five times with random seeding; only the top scoring, top-ranked model is depicted.

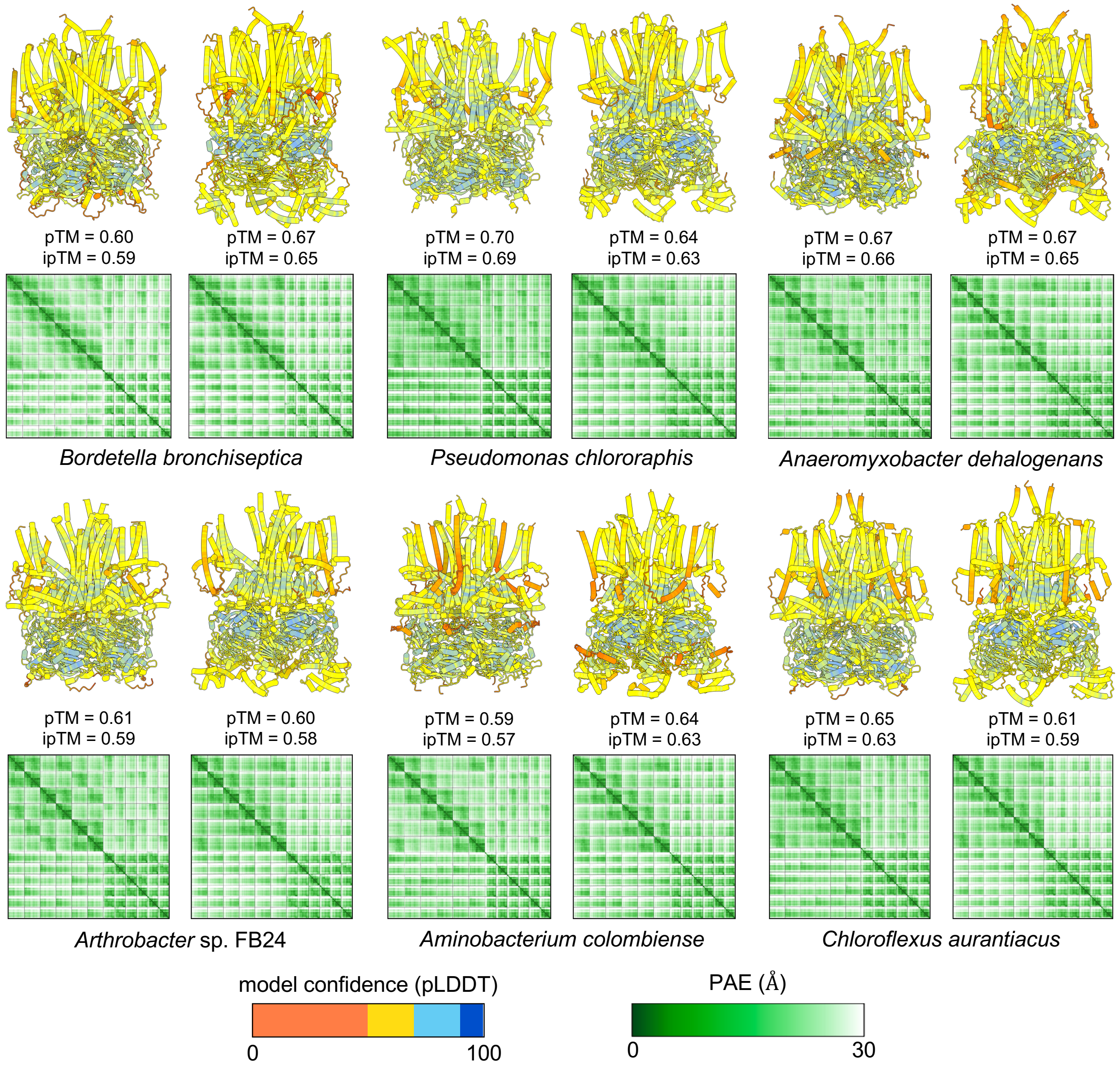


**Fig. S3: The confidence metrics generated by AF3 for the Tad motor models depicted in Figure S2 are comparable between extension and retraction orientations.** For each motor ortholog, the extension orientation is shown on the left and the retraction orientation is shown on the right. All predictions were performed a minimum of five times with random seeding; only the top scoring, top-ranked model is depicted. Models are coloured according to the predicted local distance difference test (pLDDT) scores (legend at bottom left). Below each model, the corresponding predicted aligned error (PAE) plots are depicted (legend at bottom right). The predicted template modelling (pTM) and interface predicted template modelling (ipTM) scores are provided below each model.

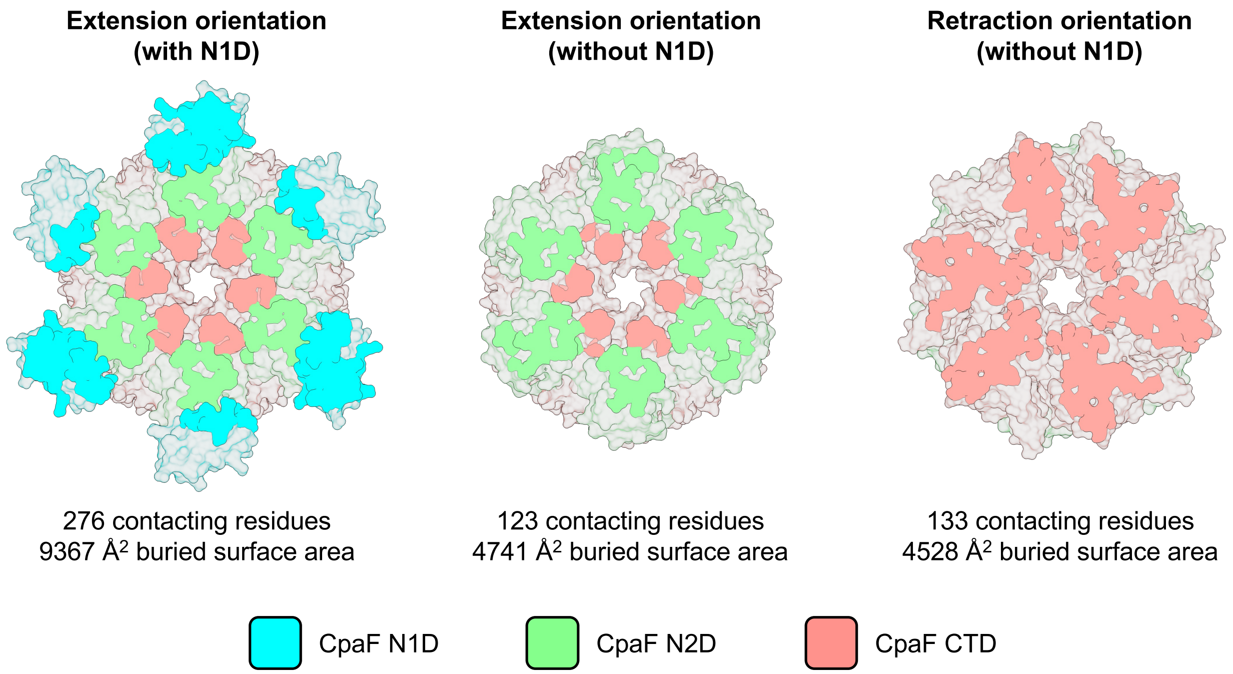


**Fig. S4: Removal of the CpaF N1 domain reduces modeling bias towards the extension conformation of the Tad motor.** *Stenotrophomonas maltophilia* CpaF was chosen as a representative example of CpaF orthologs that could only be modelled in the retraction conformation when the N1D was removed (see Table S1). *Left*: The structure of the *S. maltophilia* CpaF hexamer predicted by AF3, viewed from the N-terminal face. *Middle*: The structure of the *S. maltophilia* CpaF hexamer with the N1 domain removed, viewed from the N-terminal face. *Right*: The structure of the *S. maltophilia* CpaF hexamer with the N1 domain removed, viewed from the C-terminal face. For each model, the platform protein interaction interface is highlighted by colour, while surfaces of the protein that are not involved in platform protein interactions are transparent (grey). The number of contacting residues between the ATPase and platform complexes and the total surface area buried by the interaction are indicated below each model. Colours correspond to specific protein domains, as indicated in the legend at the bottom. N1D, variable N-terminal domain; N2D, conserved N-terminal domain; CTD, C-terminal domain.

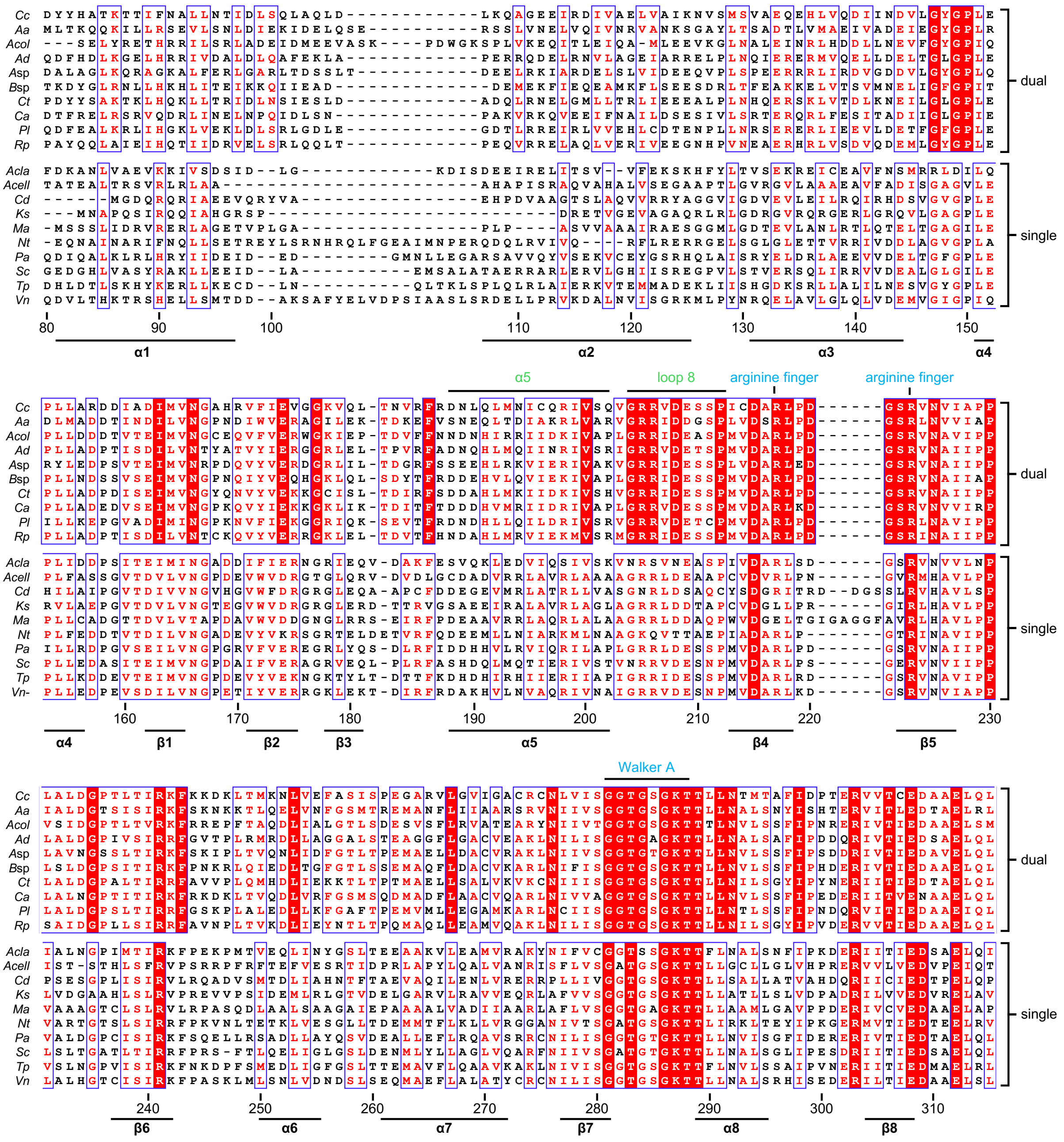


**alignment continued on next page**

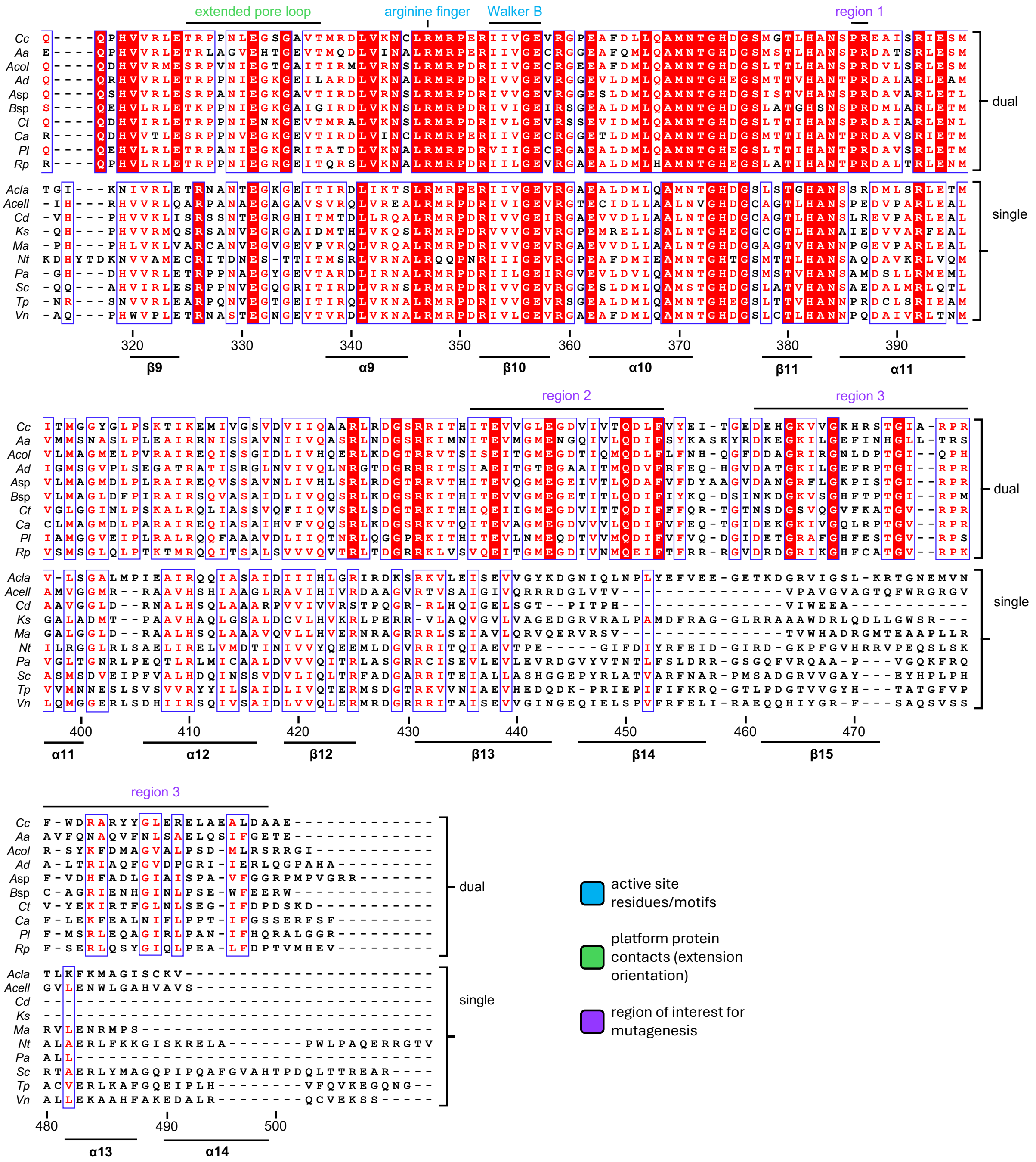


**Fig. S5: Sequence differences between single- and dual-orientation CpaF orthologs are restricted to the C-terminal region**. Sequence alignment of ten representative CpaF orthologs that are predicted by AF3 to adopt extension and retraction orientations (dual) compared to the sequence alignment of ten representative CpaF orthologs that are predicted by AF3 to adopt only the extension orientation (single). Positions with invariant residues are coloured white and highlighted with a red background, positions with ≥ 70% equivalent residues are coloured red. Residues or sequence motifs important for ATPase activity are highlighted in blue above the sequence, while motifs predicted to be important for interactions with the platform proteins in the extension conformation are highlighted in green. Regions of sequence difference between single and dual orientation motors that were targeted for further structural and mutagenic analysis are highlighted in purple. Residue numbering and secondary structural elements, based on the *Caulobacter crescentus* sequence and structure, are indicated below the sequence. *Cc*, *C. crescentus*, *Aa*, *Aggregatibacter actinomycetemcomitans*; *Acol*, *Aminobacterium colombiense*; *Ad*, *Anaeromyxobacter dehalogenans*; *A*sp, *Arthrobacter* sp. FB24; *B*sp, *Bacillus* sp. 1NLA3E; *Ct*, *Chlorobaculum tepidum*; *Ca*, *Chloroflexus aurantiacus*; *Pl*, *Planctopirus limnophila*; *Rp*, *Ralstonia pseudosolanacearum*; *Acla*, *Acetivibrio clariflavus*; *Cd*, *Corynebacterium diphtheriae*; *Ks*, *Kytococcus sedentarius*; *Ma*, *Mycobacterium avium*; *Nt*, *Novibacillus thermophilus*; *Pa*, *Pseudomonas aeruginosa*; *Sc*, *Streptomyces coelicolor*; *Tp*, *Thermincola potens*; *Vn*, *Vibrio natriegens*.

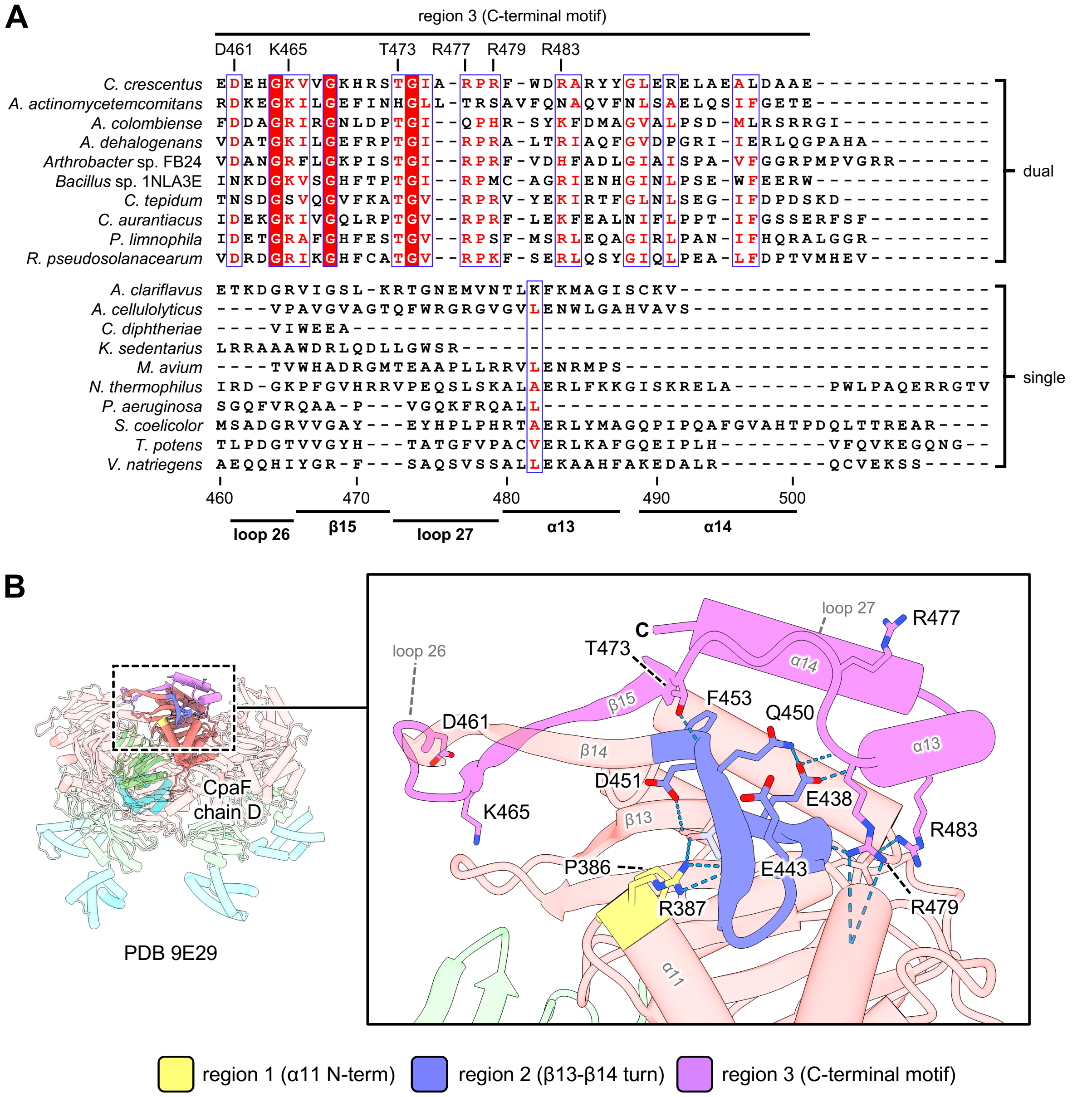


**Fig. S6: Residues conserved in dual orientation CpaF orthologs versus their single-orientation counterparts are located on the C-terminal face of the hexamer.** (**A**) Sequence comparison of the C-terminal motif (residues 460-501) of CpaF orthologs that are predicted by AF3 to adopt extension and retraction orientations (dual, top) and ten representative CpaF orthologs that are predicted by AF3 to adopt only the extension orientation (single, bottom). Positions with invariant residues are coloured white and highlighted with a red background, positions with ≥ 70% equivalent residues are coloured red. Residues of interest that are conserved in the dual-orientation alignment but not the single-orientation alignment are indicated at the top. Residue numbering and secondary structural elements, based on the *C. crescentus* sequence and structure, are indicated at the bottom. (**B**) The structure of the *C. crescentus* CpaF hexamer in the expanded conformation (PDB 9E29) highlighting conserved residues on chain D of the C-terminal face. Predicted hydrogen bonding interactions are highlighted with blue pseudobonds. Structural motifs are coloured as indicated in the legend at the bottom.

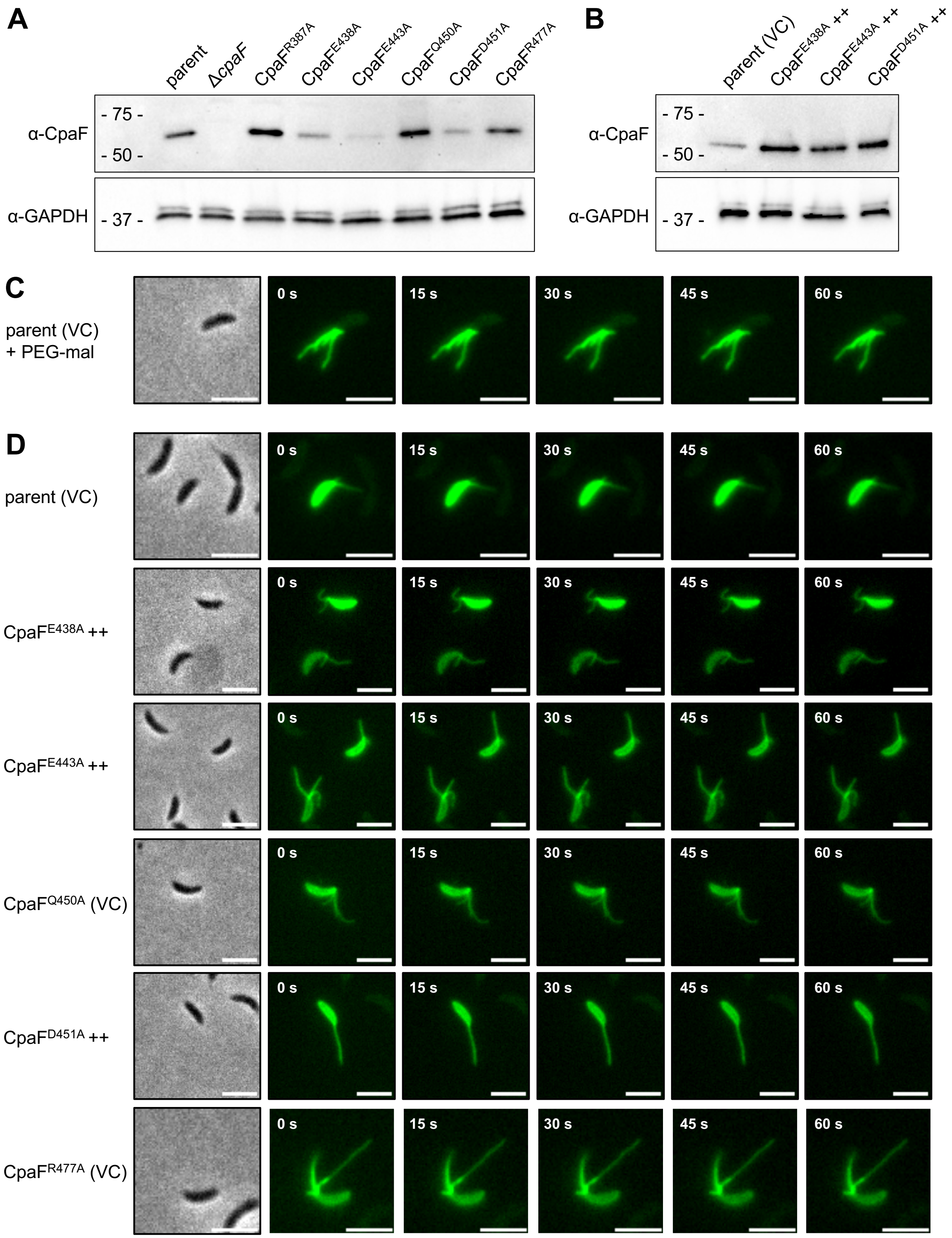


**Fig. S7: Non-retracting pili are observed in populations of all analyzed *C. crescentus* strains.** (**A**) Western blot depicting the expression of the indicated chromosomal CpaF point mutations from *C. crescentus* whole cell lysates probed using CpaF-specific antibodies. (**B**) Western blot depicting the elevated expression of the indicated CpaF point mutants expressed from a replicating plasmid from *C. crescentus* whole cell lysates probed using CpaF-specific antibodies. For panels A and B, the antibody against GAPDH was used as a loading control. (**C**) Representative time-lapse images of the parental strain where pili are labelled with AF488-maleimide (green) while pilus retraction is blocked by PEG5000-maleimide (PEG-mal). Maleimide spontaneously reacts with an engineered cysteine residue in the major pilin subunit, PilA. (**D**) Representative time-lapse images of the indicated strains labelled with AF488-maleimide (green). A representative phase contrast image of each time-series is shown on the left. Scale bars, 3 μm. Parent = NA1000 *pilA*$^{T36C}$. ++, overexpression of the indicated CpaF point mutant from pJC585 in the equivalent chromosomal point mutant background; VC, empty pJC585$^{-}$ vector control.

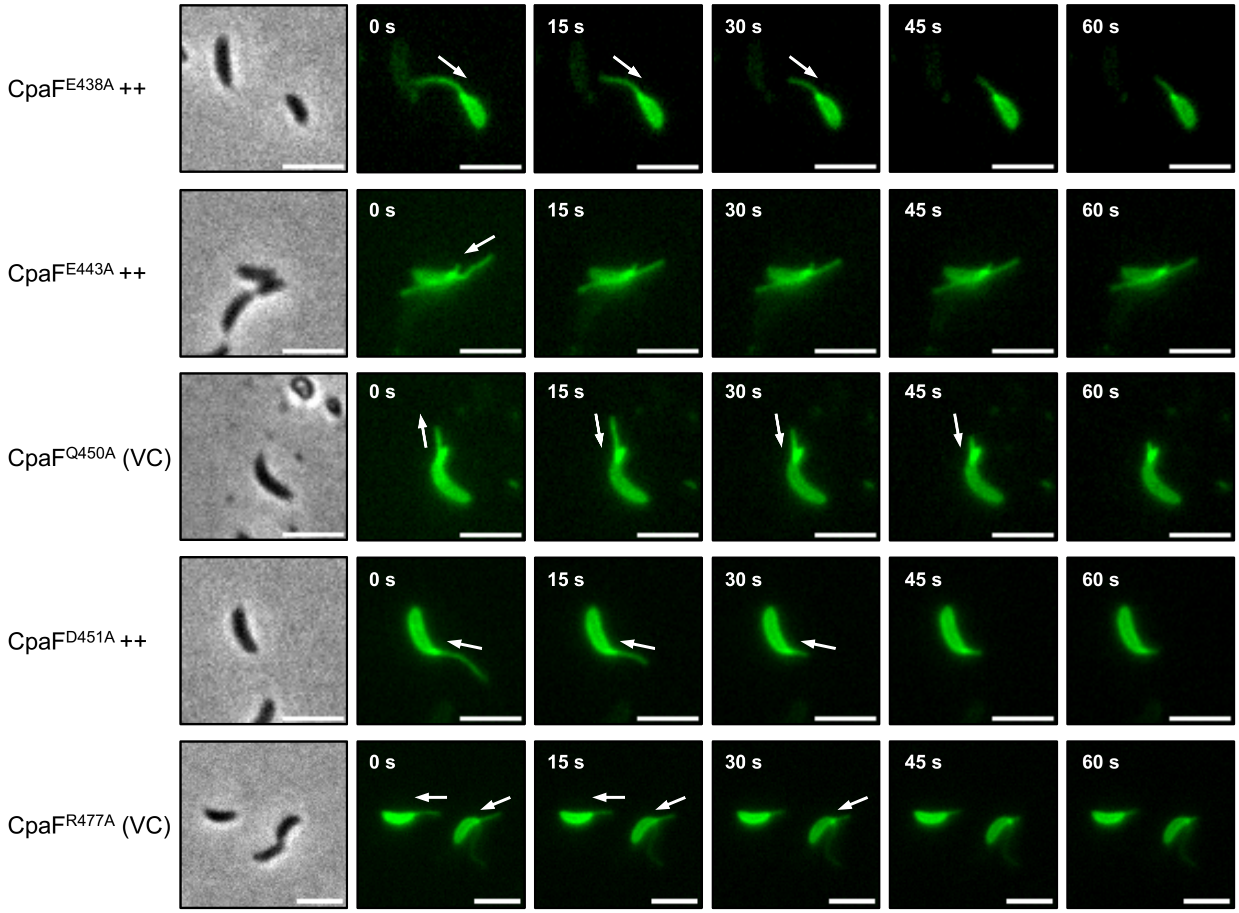


**Fig. S8: Pilus retraction is still observed in CpaF point mutants with intermediate retraction phenotypes.** Representative time-lapse images of the indicated strains where pili are labelled with AF488-maleimide (green). Maleimide spontaneously reacts with an engineered cysteine residue in the major pilin subunit, PilA. A representative phase contrast image of each time-series is shown on the left. White arrows indicate the direction of movement (extension or retraction) of pili. Scale bars, 3 µm. ++, overexpression of the indicated CpaF point mutant from pJC585 in the equivalent chromosomal point mutant background; VC, empty pJC585$^{-}$ vector control.

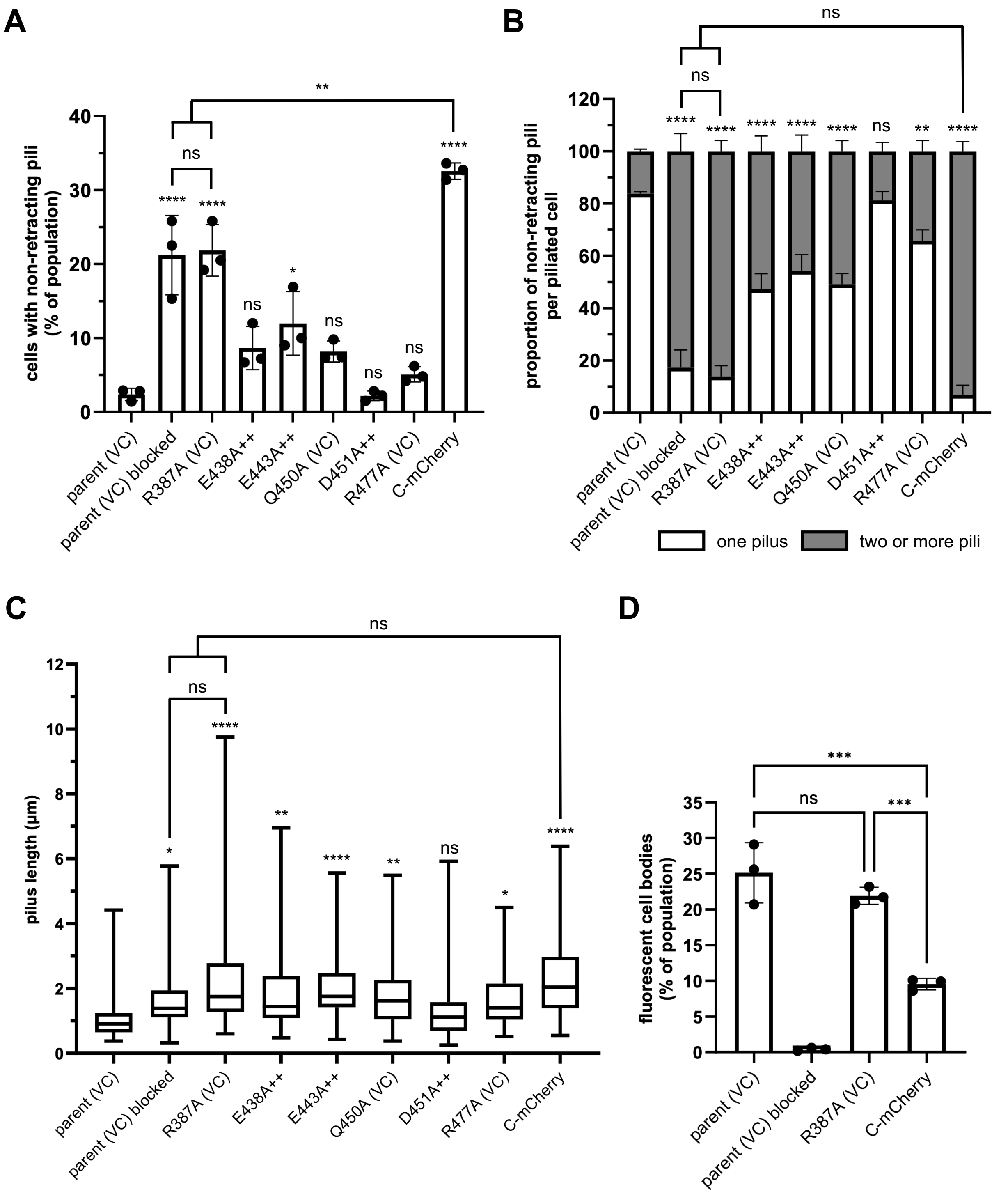


**Fig. S9: Mutation of conserved residues on the C-terminal face of CpaF or sterically blocking access to this surface through the addition of a bulky tag increases the abundance and length of non-retracting pili in *C. crescentus* populations.** (**A**) Quantification of the percentage of cells that produce pili that do not retract from mixed populations of the indicated *C. crescentus* strains labelled with AF488-maleimide. Results are the mean of three independent biological replicates with at least 500 cells analyzed per replicate. Error bars represent the standard error of the mean

(SEM). (**B)** Quantification of the number of non-retracting pili produced per piliated cell of the indicated strains, measured from mixed populations of cells labelled with AF488-maleimide. Results are the mean of three independent biological replicates with at least 50 piliated cells analyzed per replicate. Error bars represent the SEM. **(C)** Average length of non-retracting pili produced by mixed populations of the indicated strains labelled with AF488-maleimide. Sixty pili were measured for each strain. Error bars indicate the minimum to maximum range of pilus lengths. (**D)** Quantification of the percentage of cells with fluorescent cell bodies in mixed populations of the indicated strains labeled with AF488-maleimide. Results are the mean of three independent biological replicates with at least 500 cells analyzed per replicate. Error bars represent the SEM. Statistical comparisons were made using a one-way (panels A, C, and D) or two-way (panel B) ANOVA with Tukey's multiple comparisons test. ****, $P < 0.0001$; ***, $P < 0.001$; **, $P < 0.01$; *, $P < 0.05$; ns, no significant difference. In all panels, statistical test results located immediately above the error bars are for the comparison to the parental strain. Parent = NA1000 *pilA*$^{T36C}$; blocked = addition of PEG5000-maleimide to artificially block pilus retraction. ++, overexpression of the indicated CpaF point mutant from pJC585 in the equivalent chromosomal point mutant background; VC, empty pJC585$^{-}$ vector control.

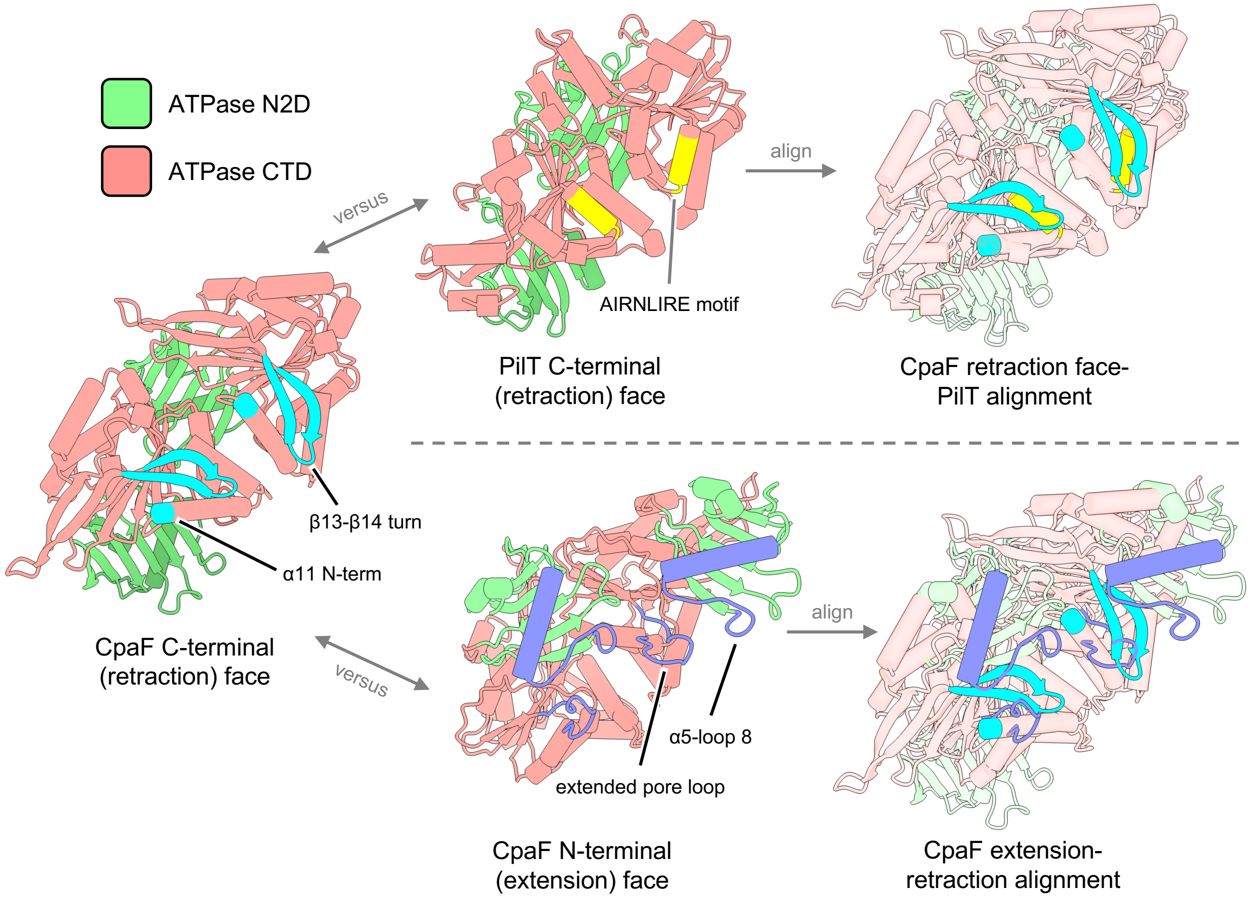


**Fig. S10: Comparison of extension- and retraction-specific structural motifs of CpaF and the retraction ATPase PilT.** *Left*: The C-terminal face of the *Caulobacter crescentus* CpaF hexamer with the indicated retraction-specific structural motifs highlighted in light blue. *Top middle*: The C-terminal face of the *Pseudomonas aeruginosa* retraction-specific ATPase PilT with the conserved AIRNLIRE motif highlighted in yellow. *Top right*: Alignment of CpaF and PilT, as viewed from the C-terminal faces of the hexamers. *Bottom middle*: The N-terminal face of the CpaF hexamer with the indicated extension-specific structural motifs highlighted in purple. *Bottom right*: Alignment of the platform protein complexes (hidden) of the extension- and retraction-specific Tad motors predicted by AF3, to bring the N- and C-terminal faces of the CpaF hexamer into alignment. Protein domains are coloured as indicated in the legend (top left). N2D, conserved N-terminal domain; CTD, C-terminal domain. In each structure, two adjacent chains (i.e. subunits) of the hexameric complexes are shown, and the N1 domains have been omitted from the models for clarity.

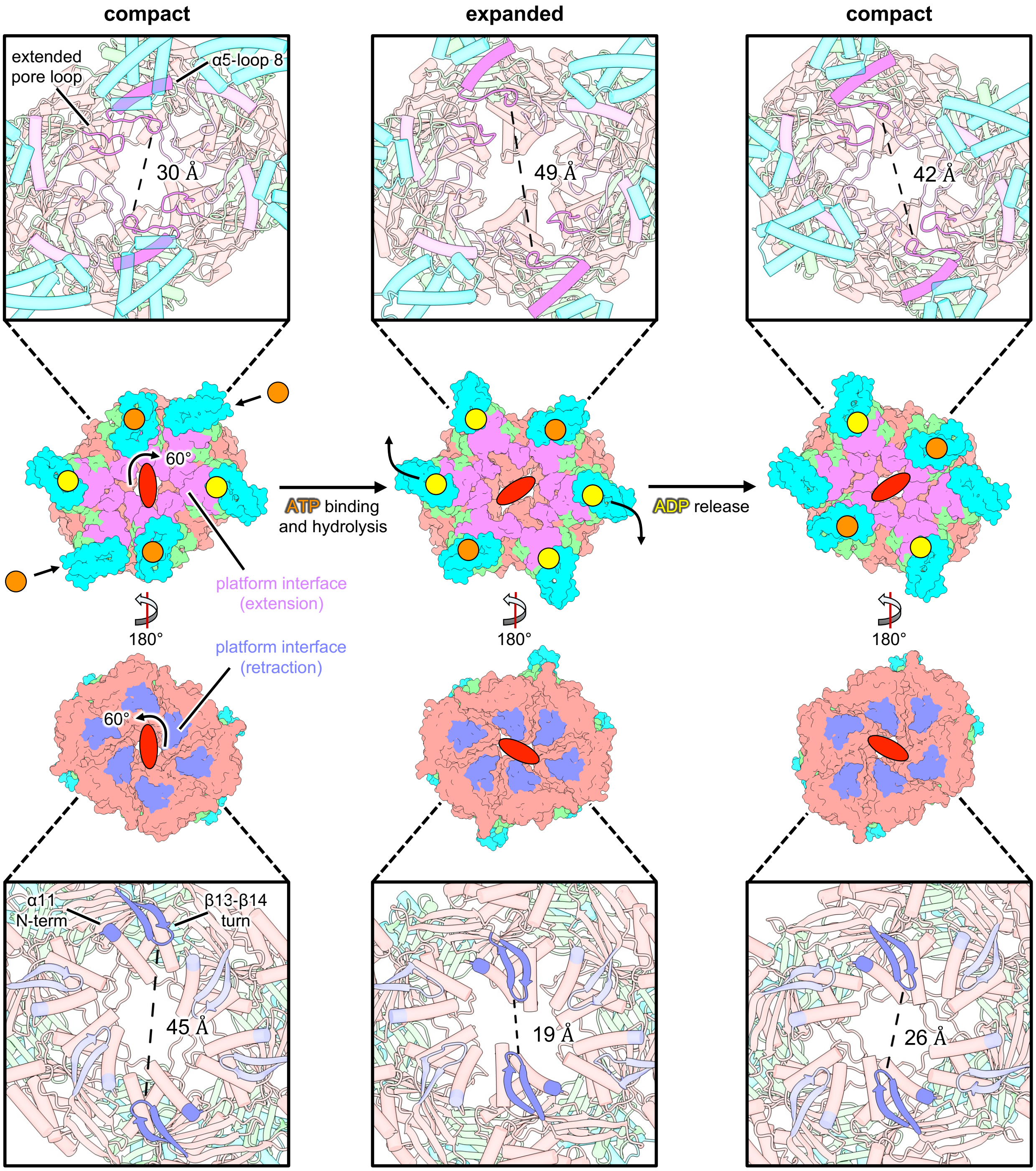


**Fig. S11: The extension- and retraction-specific surfaces of the CpaF hexamer undergo opposing inter-chain conformational changes during nucleotide turnover.** *Top*: Conformational transitions of CpaF during one round of catalysis as viewed from the N-terminal face of the hexamer. The C2 axis of symmetry (red oval) undergoes a 60° clockwise rotation as the compact conformation (PDB 9E26) transitions to the expanded conformation (PDB 9E29). The inset depicts the relative positions of opposing extension-specific platform interaction motifs (α5-loop 8 and extended pore loop, magenta) during nucleotide turnover. *Bottom*: Conformational transitions of CpaF viewed from the C-terminal face of the hexamer. The C2 axis of symmetry undergoes a 60° counterclockwise rotation during nucleotide turnover. The inset depicts the relative positions of opposing retraction-specific platform interaction motifs (α11 N-term and β13-β14 turn, purple) during nucleotide turnover. The movements of the retraction motifs directly oppose those of the extension motifs shown at the top. CpaF domains are coloured as indicated in Figure 1. Yellow circles represent ADP; orange circles represent ATP.

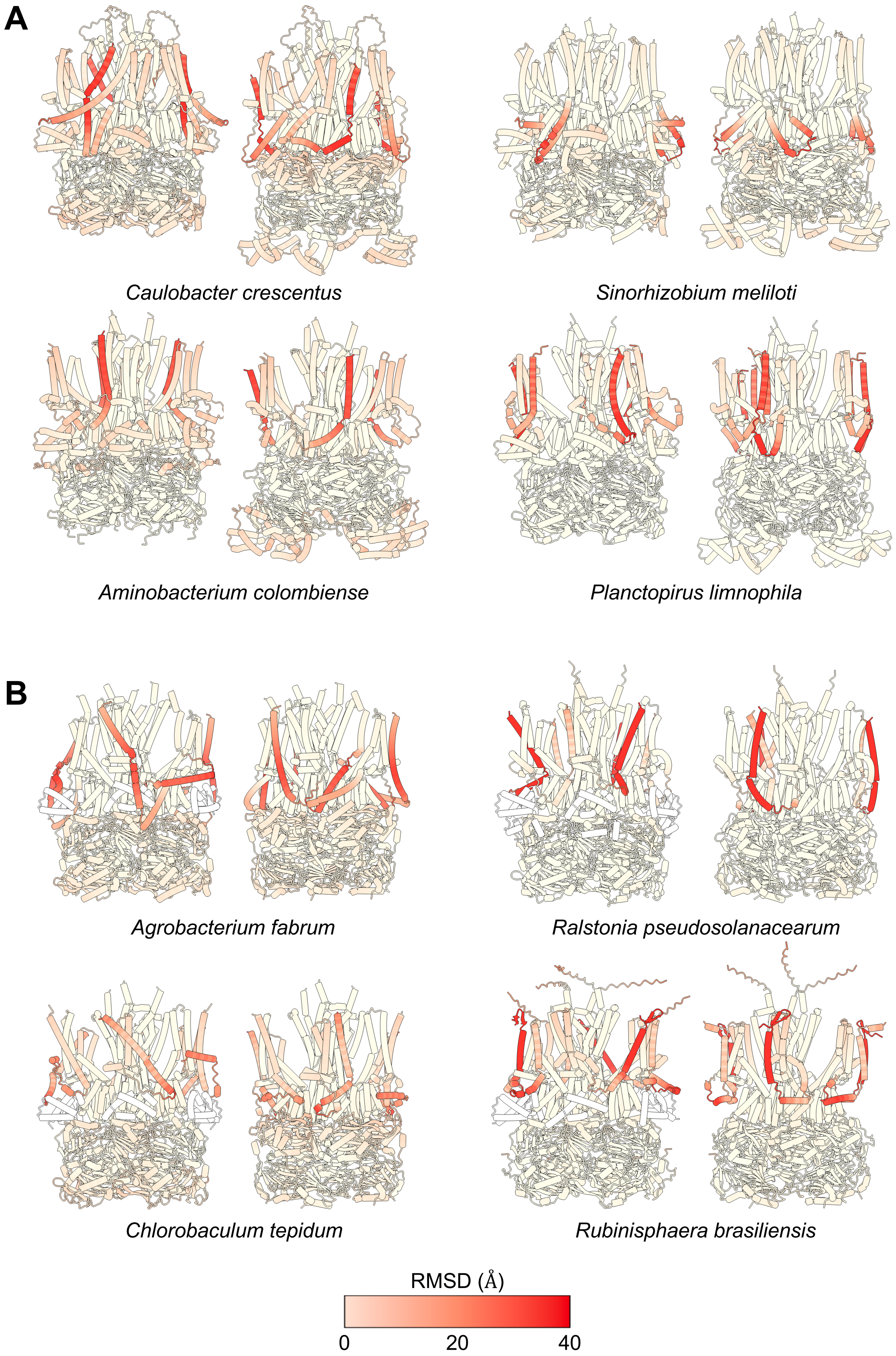
A
Caulobacter crescentus
Sinorhizobium meliloti
Aminobacterium colombiense
Planctopirus limnophila
B
Agrobacterium fabrum
Ralstonia pseudosolanacearum
Chlorobaculum tepidum
Rubinisphaera brasiliensis
RMSD (Å)
0
20
40

**Fig. S12: The platform protein N-termini undergo the largest conformational changes between AF3-predicted extension and retraction motor models.** Models of the extension (left) and retraction (right) conformations of Tad motor orthologs from representative bacterial species were aligned and coloured according to the root mean square deviation (RMSD, legend at bottom) at each amino acid position. The platform protein complex and the CpaF hexameric complex were compared independently to exclude conformational deviations due to ATPase inversion. Motors that were modelled in both orientations with the CpaF N1 domain present (**A**), as well as those that could only be modelled in both conformations after removal of the N1 domain (**B**) are shown.

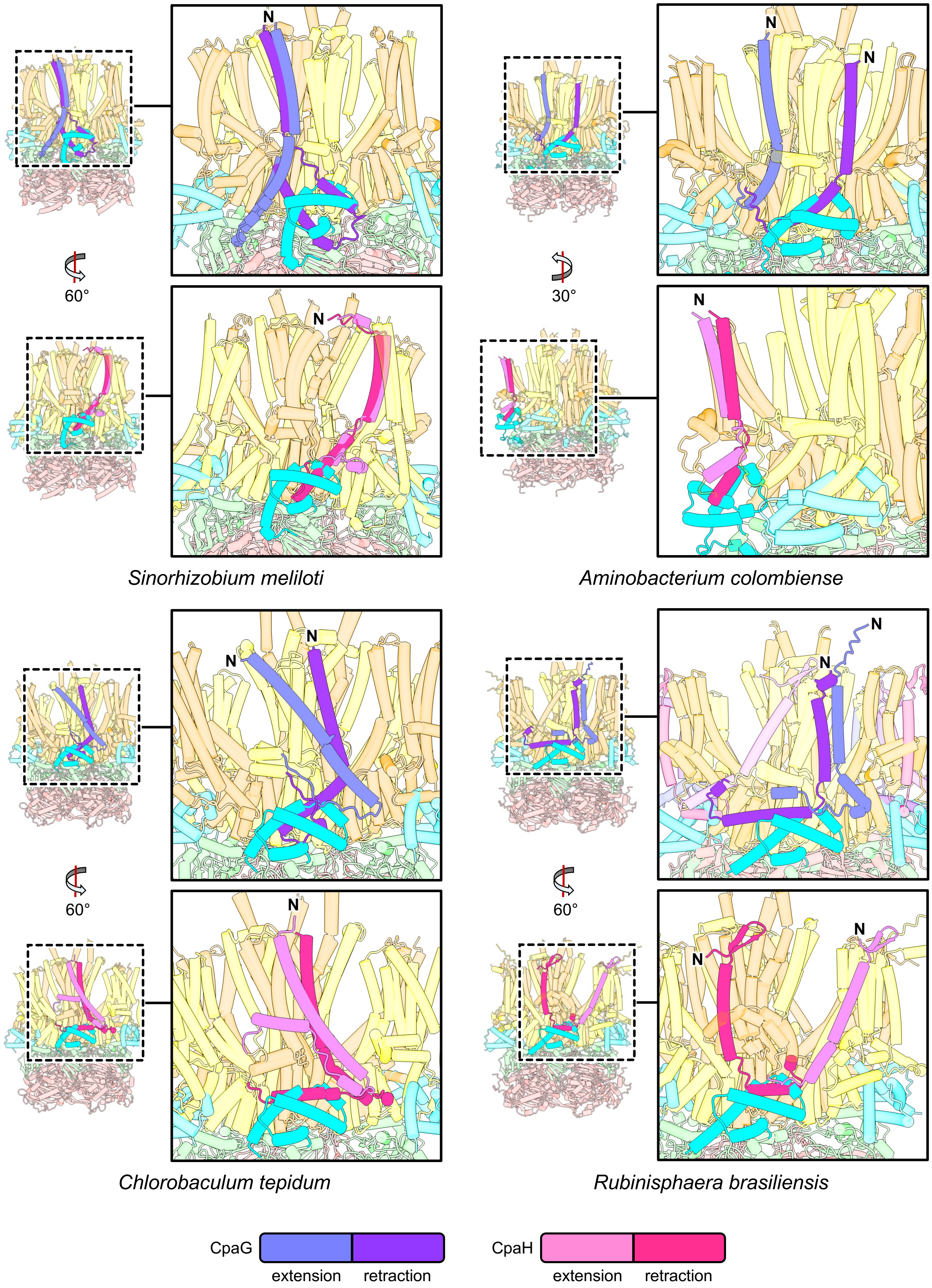
N
N
N
60°
30°
N
N
Sinorhizobium meliloti
Aminobacterium colombiense
N
N
N
N
60°
60°
N
N
N
Chlorobaculum tepidum
Rubinisphaera brasiliensis
CpaG
extension
retraction
CpaH
extension
retraction

**Fig. S13: Representative examples of platform protein N-termini undergoing large conformational changes that could force ATPase reorientation through steric clashes with the N1 domain of CpaF.** Structural alignment of the extension orientation of the indicated Tad motor orthologs predicted by AF3 with the platform protein complex from the retraction orientation model. For each example, the top panel focuses on the conformational change of the CpaG N-terminus between the extension (light purple) and retraction (dark purple) conformations, and the bottom panel focuses on the conformational change of the CpaH N-terminus between the extension (light pink) and retraction (dark pink) conformations. Protein and protein domain colouring is as indicated in Figure 1.

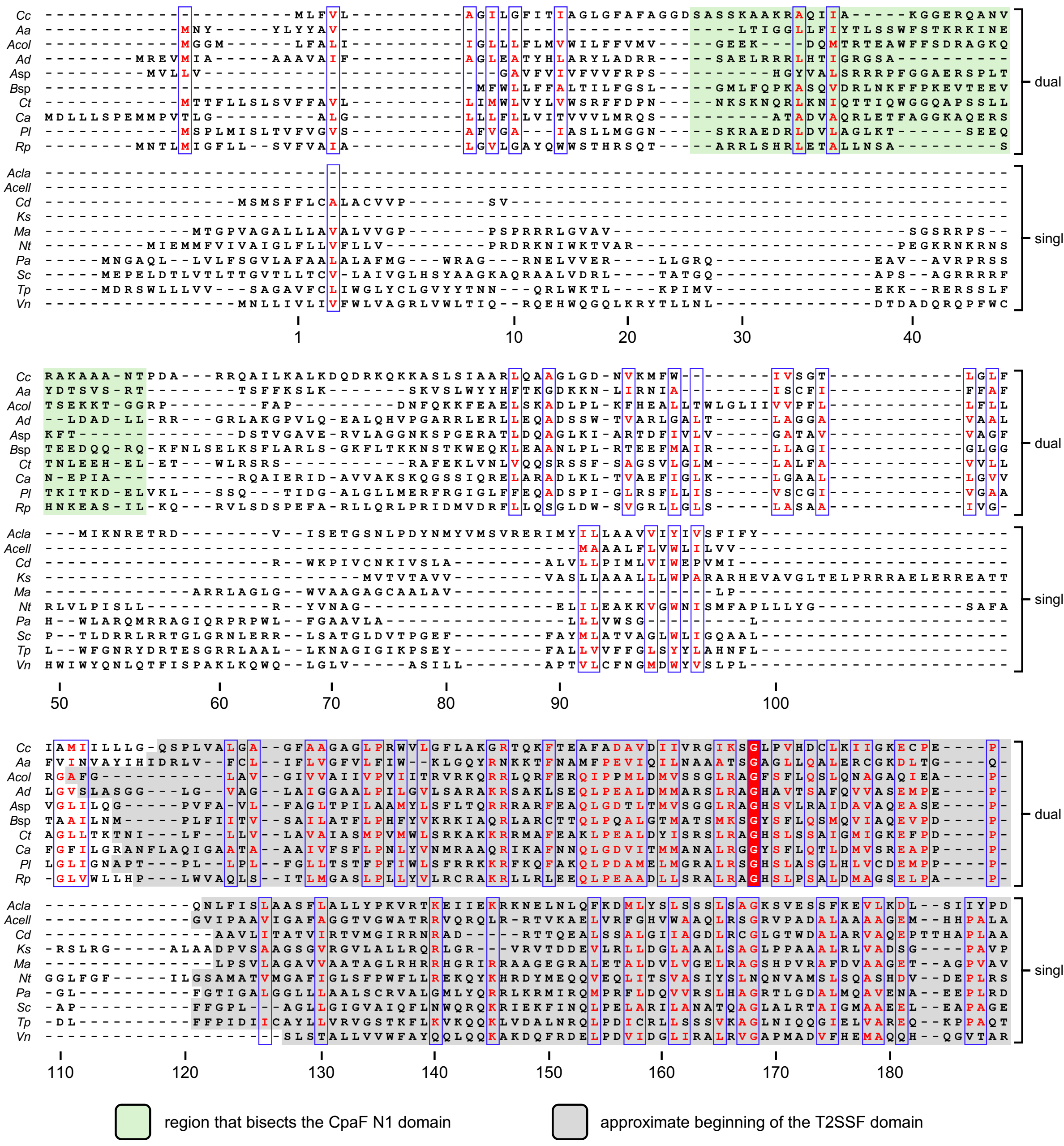


**Fig. S14: The N-terminal region adjacent to the type II secretion system protein (T2SSF) domain of CpaG orthologs is highly variable.** Sequence alignment of ten representative CpaG orthologs that are part of motors predicted by AF3 to adopt extension and retraction orientations (dual) compared to the sequence alignment of ten representative CpaG orthologs that are part of motors predicted by AF3 to adopt only the extension orientation (single). Positions with invariant residues are coloured white and highlighted with a red background, positions with ≥ 70% equivalent residues are coloured red. The region of the CpaG N-terminal domain that bisects the N1 domain of CpaF in retraction models is highlighted with a green background, while the beginning of the conserved T2SSF domain is indicated with a grey background. Residue numbering, based on the *Caulobacter crescentus* protein, is indicated below the sequence. *Cc*, *C. crescentus*, *Aa*, *Aggregatibacter actinomycetemcomitans*; *Acol*, *Aminobacterium colombiense*; *Ad*, *Anaeromyxobacter dehalogenans*; *A*sp, *Arthrobacter* sp. FB24; *B*sp, *Bacillus* sp. 1NLA3E; *Ct*,

*Chlorobaculum tepidum*; *Ca*, *Chloroflexus aurantiacus*; *Pl*, *Planctopirus limnophila*; *Rp*, *Ralstonia pseudosolanacearum*; *Acla*, *Acetivibrio clariflavus*; *Cd*, *Corynebacterium diphtheriae*; *Ks*, *Kytococcus sedentarius*; *Ma*, *Mycobacterium avium*; *Nt*, *Novibacillus thermophilus*; *Pa*, *Pseudomonas aeruginosa*; *Sc*, *Streptomyces coelicolor*; *Tp*, *Thermincola potens*; *Vn*, *Vibrio natriegens*.

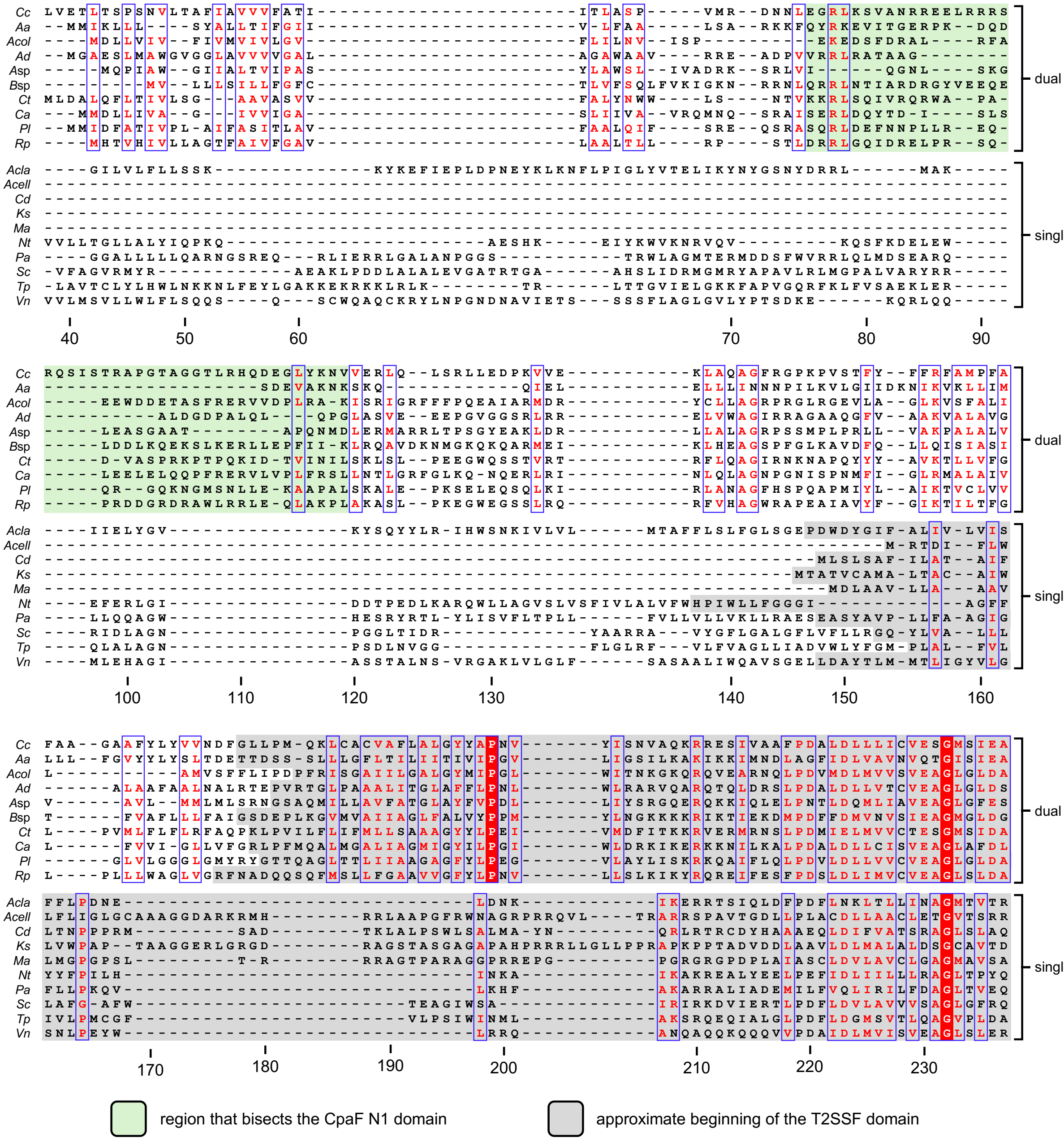


**Fig. S15: The N-terminal region adjacent to the type II secretion system protein F (T2SSF) domain of CpaH orthologs is highly variable.** Sequence alignment of ten representative CpaH orthologs that are part of motors predicted by AF3 to adopt extension and retraction orientations (dual) compared to the sequence alignment of ten representative CpaH orthologs that are part of motors predicted by AF3 to adopt only the extension orientation (single). Positions with invariant residues are coloured white and highlighted with a red background, positions with ≥ 70% equivalent residues are coloured red. The region of the CpaH N-terminal domain that bisects the N1 domain of CpaF in retraction models is highlighted with a green background, while the beginning of the conserved T2SSF domain is indicated with a grey background. Residue numbering, based on the *Caulobacter crescentus* protein, is indicated below the sequence. Numbering begins at residue 40 as the disordered N-terminal region of *C. crescentus* CpaH has been omitted. *Cc*, *C. crescentus*, *Aa*, *Aggregatibacter actinomycetemcomitans*; *Acol*,

*Aminobacterium colombiense*; *Ad*, *Anaeromyxobacter dehalogenans*; *A*sp, *Arthrobacter* sp. FB24; *B*sp, *Bacillus* sp. 1NLA3E; *Ct*, *Chlorobaculum tepidum*; *Ca*, *Chloroflexus aurantiacus*; *Pl*, *Planctopirus limnophila*; *Rp*, *Ralstonia pseudosolanacearum*; *Acla*, *Acetivibrio clariflavus*; *Cd*, *Corynebacterium diphtheriae*; *Ks*, *Kytococcus sedentarius*; *Ma*, *Mycobacterium avium*; *Nt*, *Novibacillus thermophilus*; *Pa*, *Pseudomonas aeruginosa*; *Sc*, *Streptomyces coelicolor*; *Tp*, *Thermincola potens*; *Vn*, *Vibrio natriegens*.

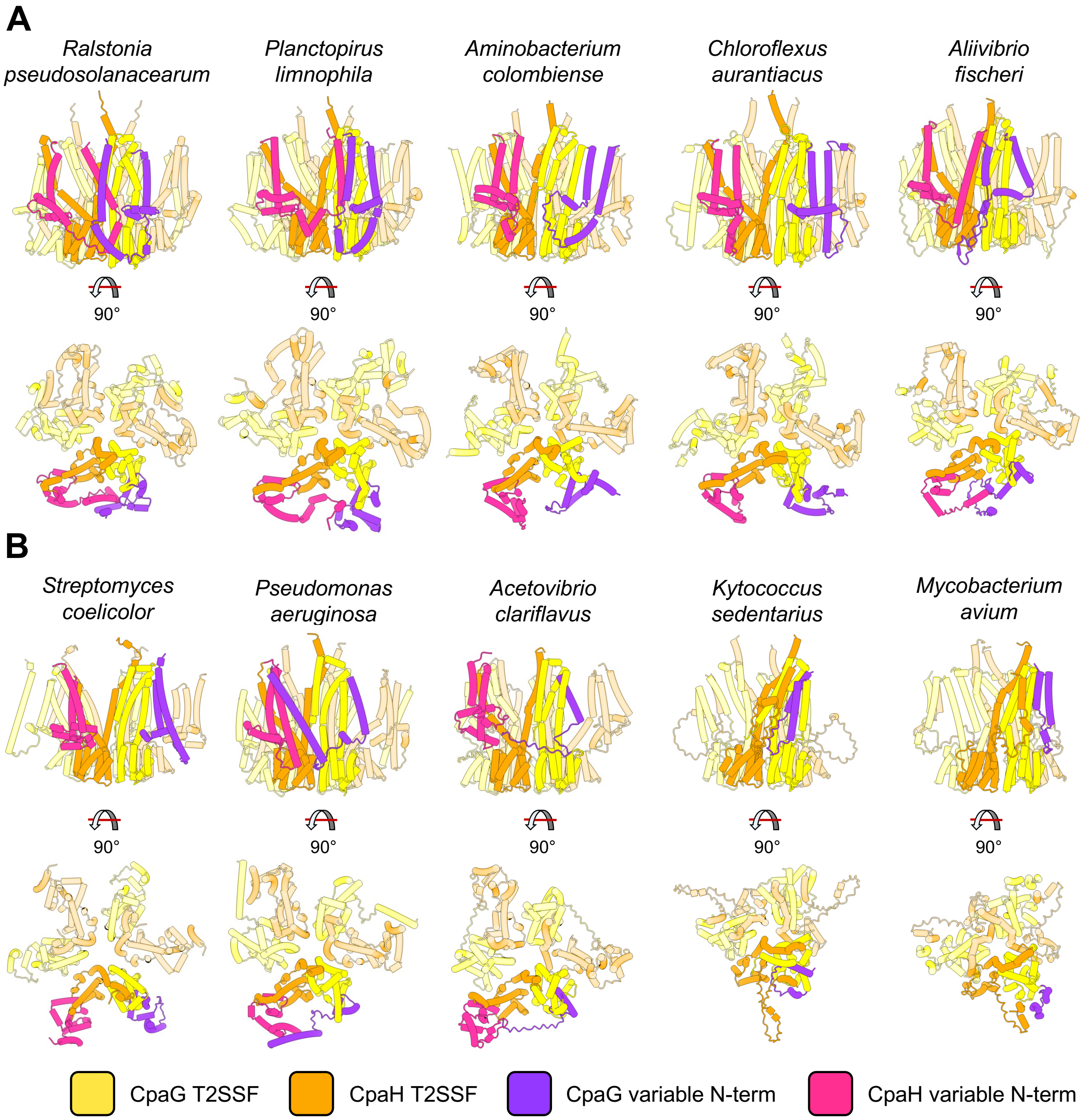


**Fig. S16: The variable N-terminal regions of single-orientation CpaG and CpaH orthologs undergo varying degrees of truncation compared to their dual orientation counterparts.** Comparison of the N-terminal region adjacent to the conserved type II secretion system protein F (T2SSF) domain of the platform proteins from (**A**) dual-orientation motors and (**B**) single-orientation motors as predicted by AF3. The N-terminal regions are coloured only for one of the three CpaG-CpaH heterodimers in the trimeric platform complex. Protein domains are coloured as indicated in the legend at the bottom.

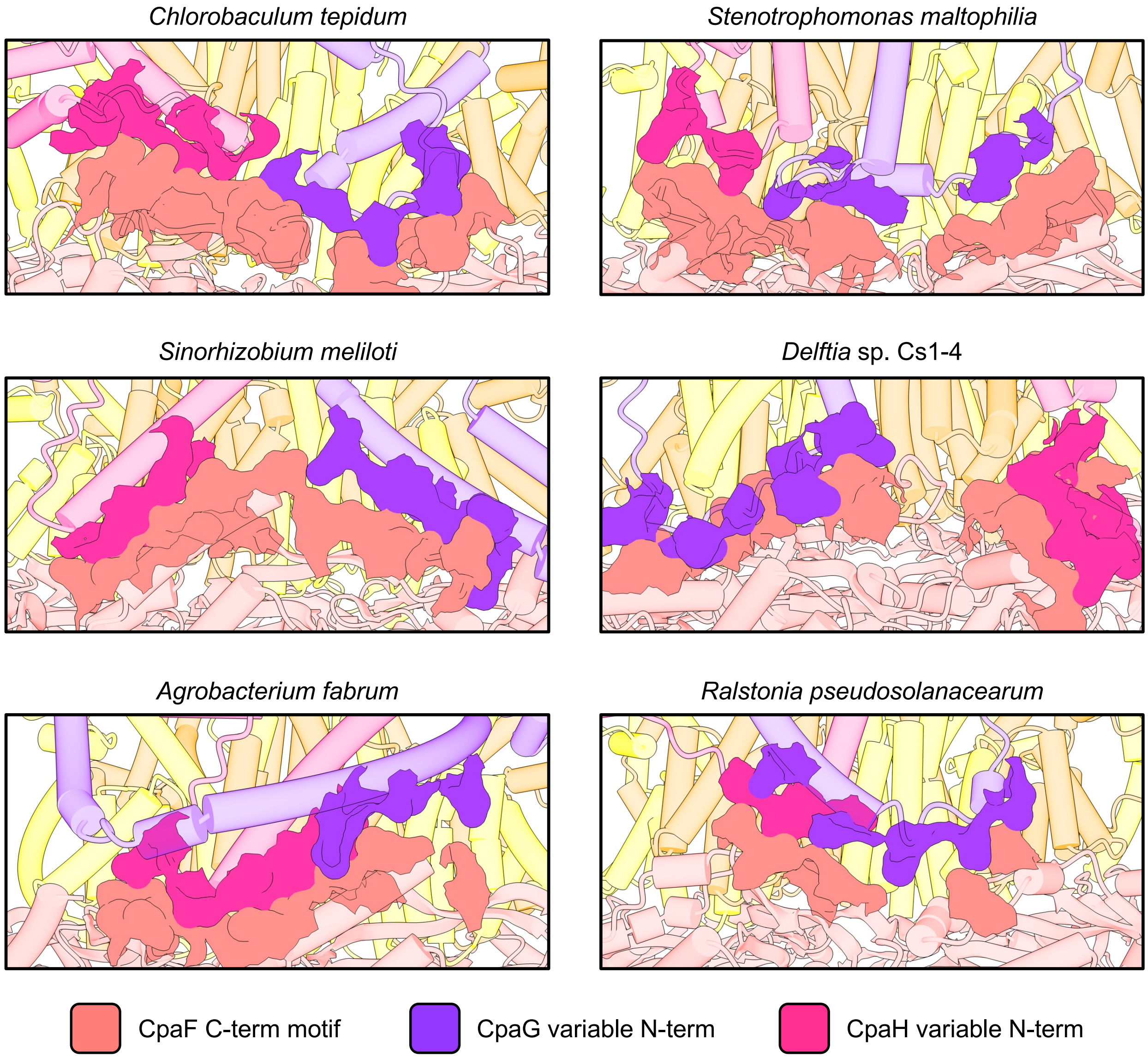


**Fig. S17: Platform protein N-termini are predicted to interact with the C-terminal motif of the ATPase in dual-orientation motors.** The interaction interface between the platform protein N-termini and the C-terminal motif of the ATPase is depicted from the retraction-orientation predictions of the indicated motors generated by AF3. Regions of interfacing residues are highlighted by the surface representation. Colouring is as indicated in the legend at the bottom.

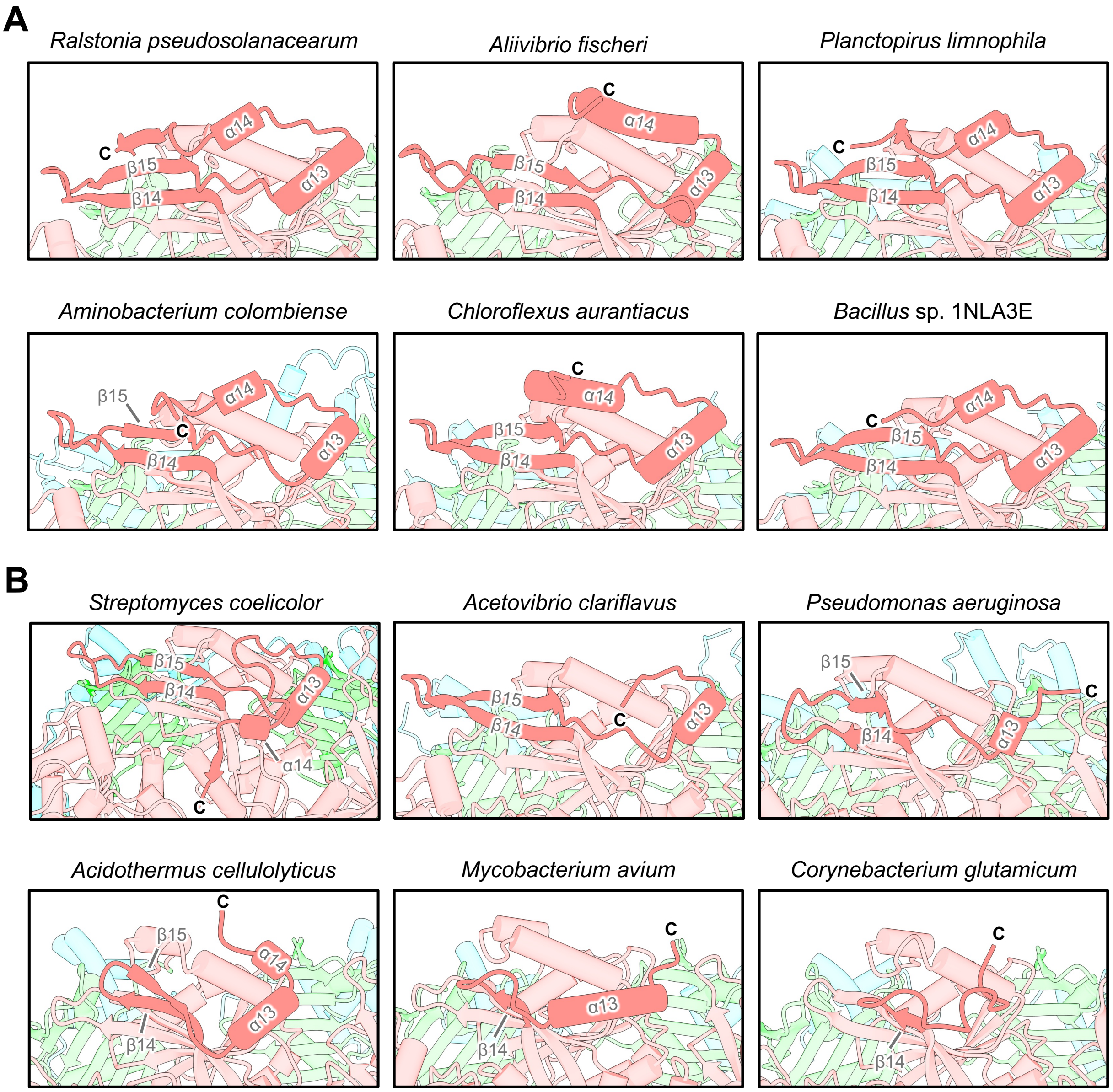


**Fig. S18: The C-terminal motif in single-orientation CpaF orthologs undergoes various degrees of structural degeneration compared to their dual-orientation counterparts.** Representative examples of the CpaF C-terminal motif from (**A**) dual-orientation motors and (**B**) single-orientation motors, as predicted by AF3.

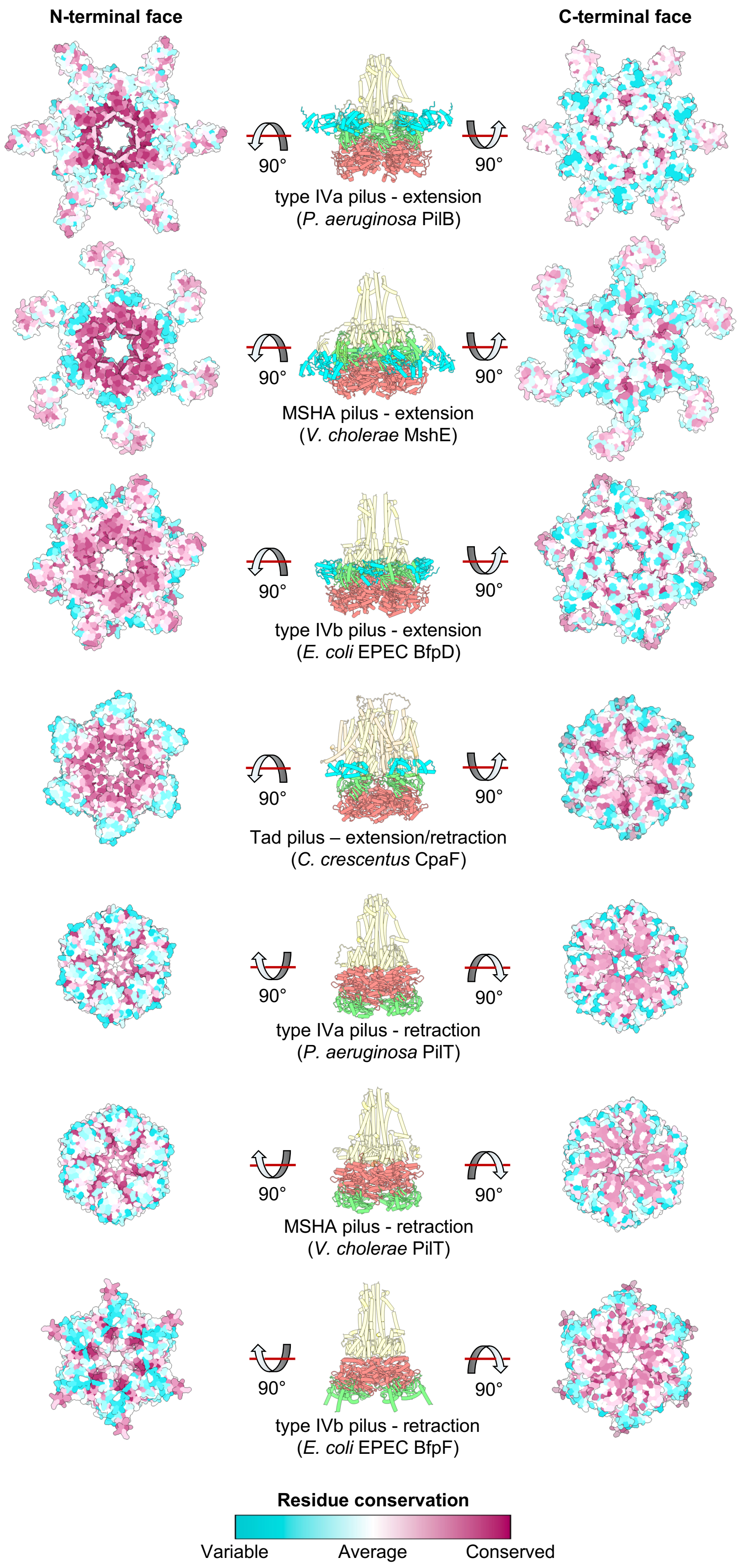

N-terminal face
C-terminal face
90°
type IVa pilus - extension
(*P. aeruginosa* PilB)
MSHA pilus - extension
(*V. cholerae* MshE)
type IVb pilus - extension
(*E. coli* EPEC BfpD)
Tad pilus – extension/retraction
(*C. crescentus* CpaF)
type IVa pilus - retraction
(*P. aeruginosa* PilT)
MSHA pilus - retraction
(*V. cholerae* PilT)
type IVb pilus - retraction
(*E. coli* EPEC BfpF)
Residue conservation
Variable
Average
Conserved

**Fig. S19: Patterns of surface residue conservation are consistent with the known activity of monofunctional and bifunctional ATPases from bacterial type IV pilus systems.** The degree of conservation of surface residues was mapped onto the AF3-predicted structures of the indicated hexameric ATPases using the Consurf server (36718848). Residues with the highest level of conservation are coloured magenta, those with average levels of conservation are coloured white, and those with the highest degree of variability are coloured turquoise, as indicated in the legend at the bottom. The N-terminal face corresponds to the hexameric surface on which the variable and/or conserved N-terminal domains of the ATPase are located, while the C-terminal face corresponds to the surface on which the conserved C-terminal domain is located. Domain colouring of the motor structure in the middle are as indicated in Figure 1.

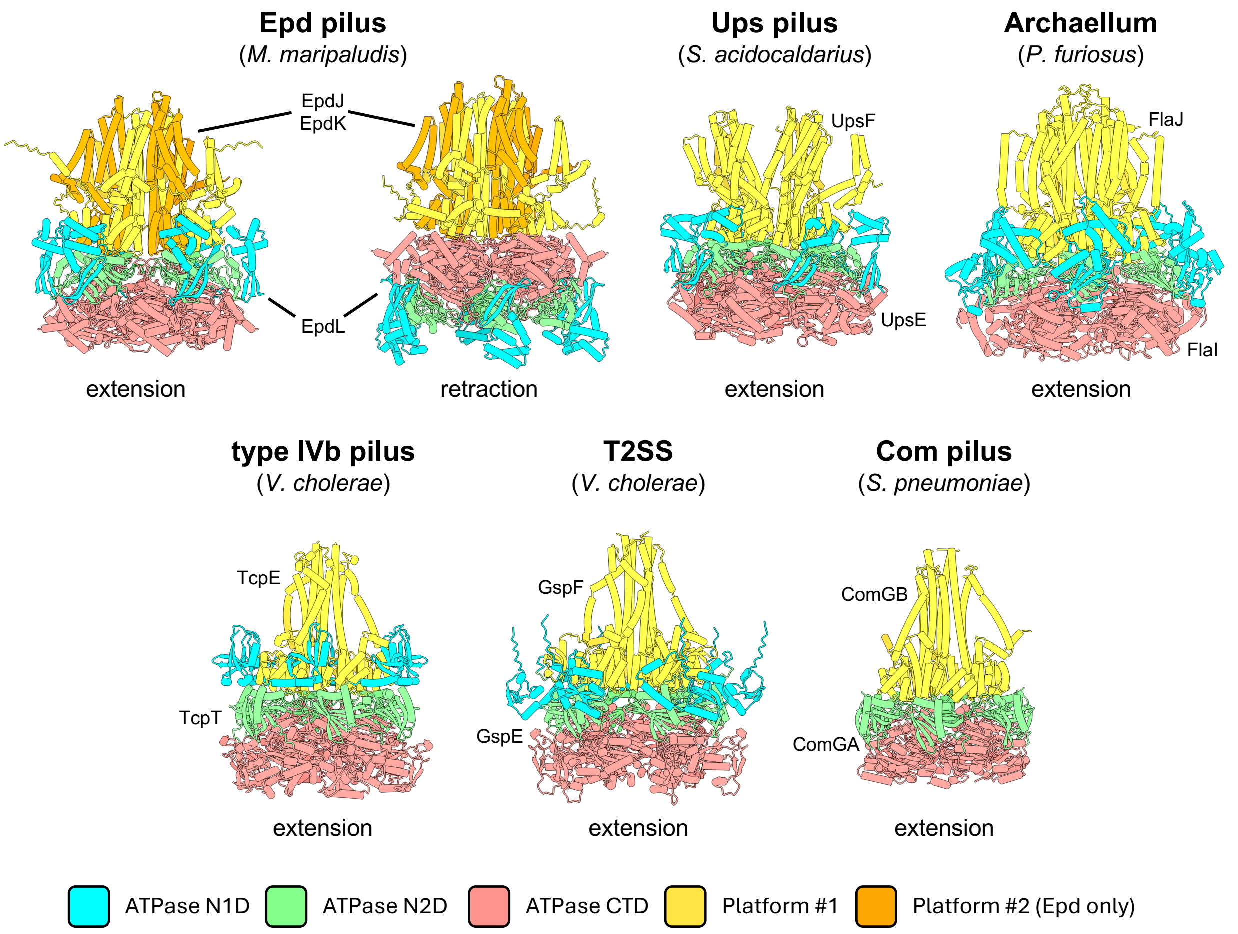


**Fig. S20: Of the single-motor type IV filament systems, only the archaeal EppA-dependent (Epd) pilus motor is predicted to adopt both extension and retraction orientations.** Structures of representative motors from single-motor type IV filament systems predicted by AF3 are shown. All predictions were performed a minimum of five times with random seeding; only the top scoring, top-ranked model is depicted. Colours correspond to specific proteins or protein domains, as shown in the legend at the bottom. The names of the specific platform proteins and ATPases are indicated beside each structure. N1D, variable N-terminal domain; N2D, conserved N-terminal domain; CTD, C-terminal domain; Ups, UV-inducible pilus system; T2SS, type II secretion system; Com, competence.

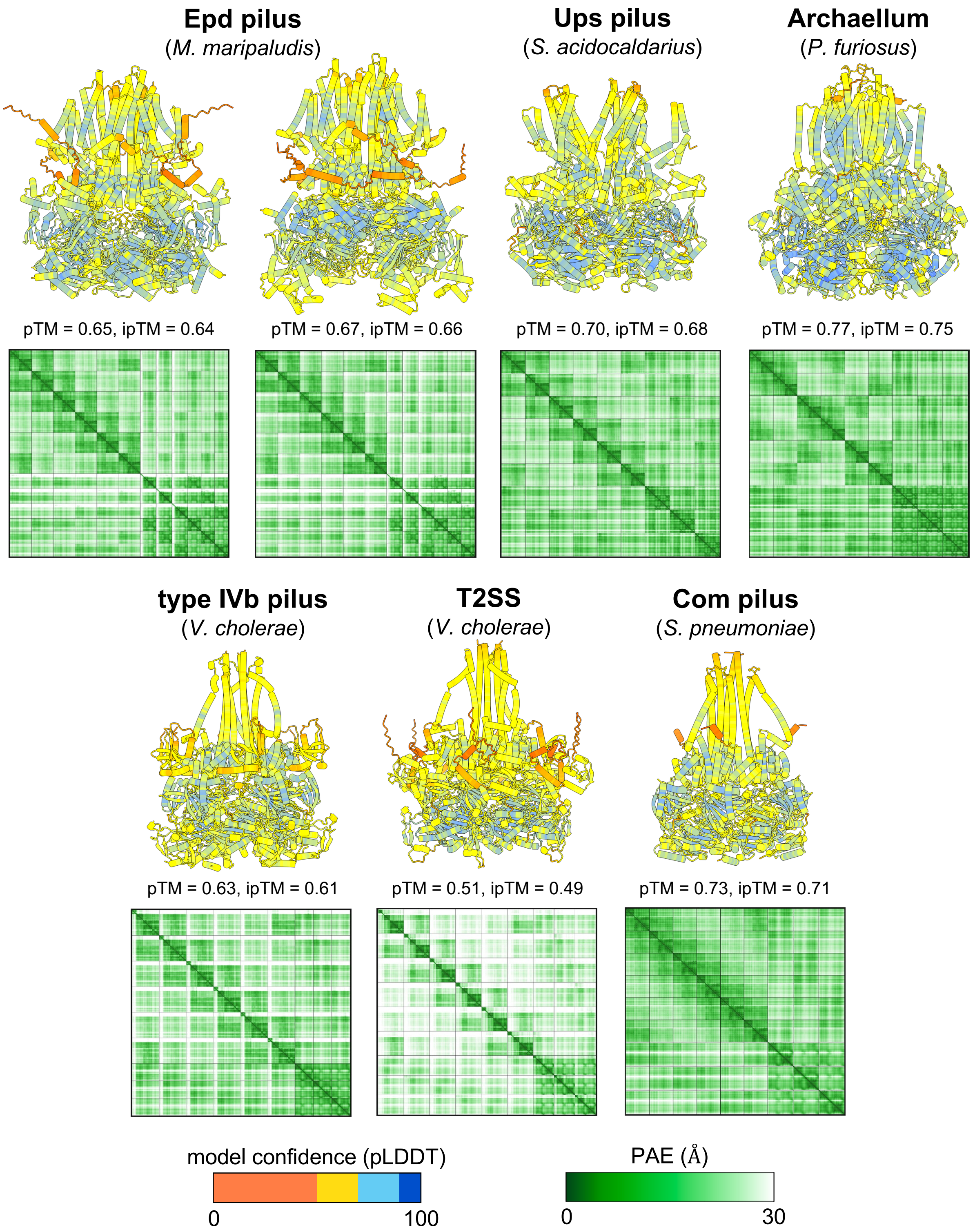


**Fig. S21: Confidence metrics generated by AF3 for the indicated type IV filament motor models depicted in Figure S20**. All predictions were performed a minimum of five times with random seeding; only the top scoring, top-ranked model is depicted. Models are coloured according to the predicted local distance difference test (pLDDT) scores (legend at bottom left). Below each model, the corresponding predicted aligned error (PAE) plots are shown (legend at bottom right). The predicted template modelling (pTM) and interface predicted template modelling (ipTM) scores are provided below each model. For the EppA-dependent (Epd) models, the extension motor is depicted on the left and the retraction motor is depicted on the right.

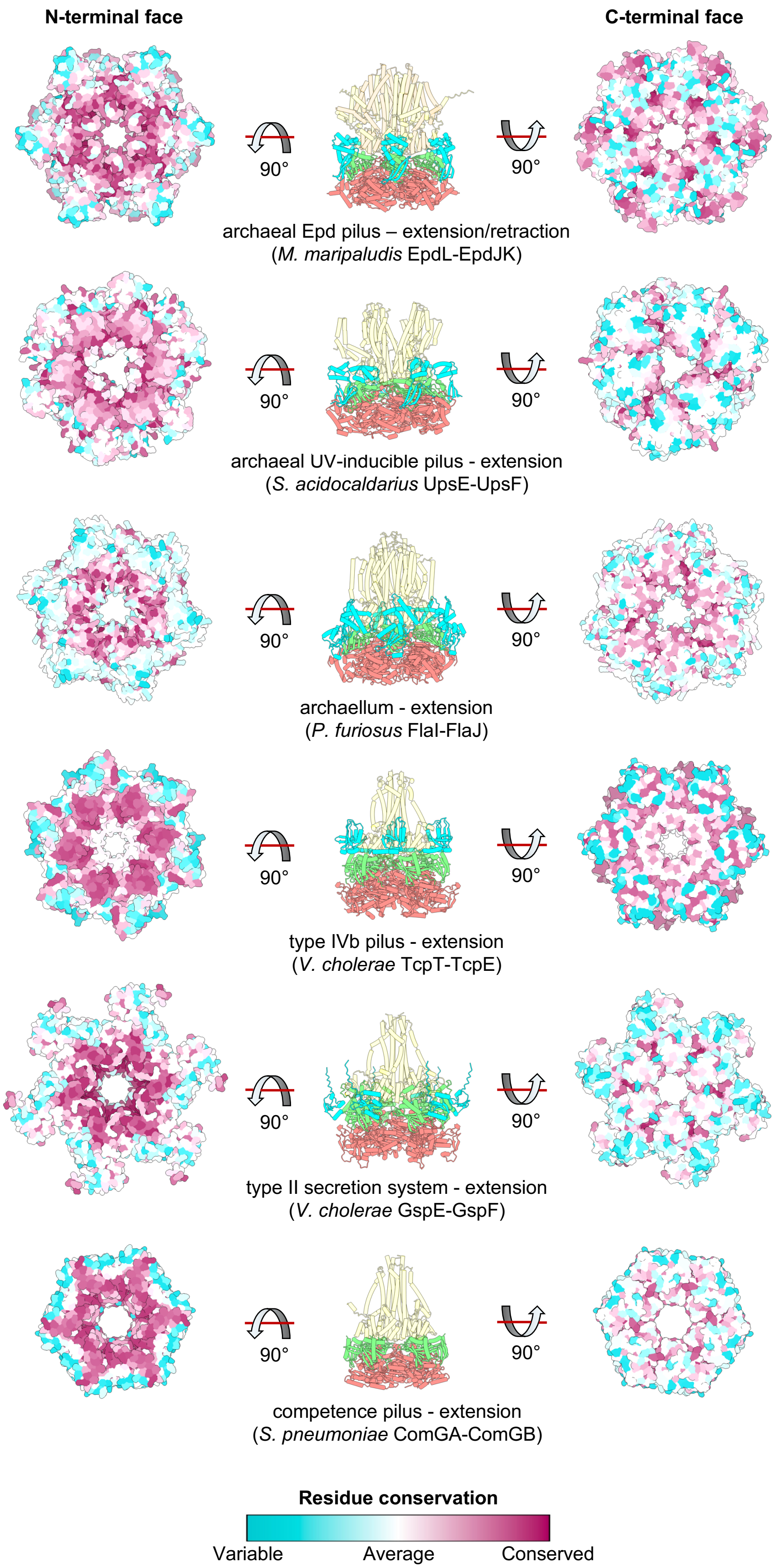
N-terminal face
C-terminal face
90°
90°
archaeal Epd pilus – extension/retraction
(*M. maripaludis* EpdL-EpdJK)
archaeal UV-inducible pilus - extension
(*S. acidocaldarius* UpsE-UpsF)
archaellum - extension
(*P. furiosus* FlaI-FlaJ)
type IVb pilus - extension
(*V. cholerae* TcpT-TcpE)
type II secretion system - extension
(*V. cholerae* GspE-GspF)
competence pilus - extension
(*S. pneumoniae* ComGA-ComGB)
Residue conservation
Variable
Average
Conserved

**Fig. S22: Patterns of surface residue conservation favour the N-terminal face of some ATPase hexamers from single-motor type IV filament systems, suggesting that they do not utilize ATPase inversion to achieve bifunctionality.** The degree of conservation of surface residues was mapped onto the AF3-predicted structures of the indicated hexameric ATPases using the Consurf server (*35*). Residues with the highest level of conservation are coloured magenta, those with average levels of conservation are coloured white, and those with the highest degree of variability are coloured turquoise, as indicated in the legend at the bottom. The N-terminal face corresponds to the hexameric surface on which the variable and/or conserved N-terminal domains of the ATPase are located, while the C-terminal face corresponds to the surface on which the conserved C-terminal domain is located. Domain colouring of the motor structures in the middle are as indicated in Figure 1.

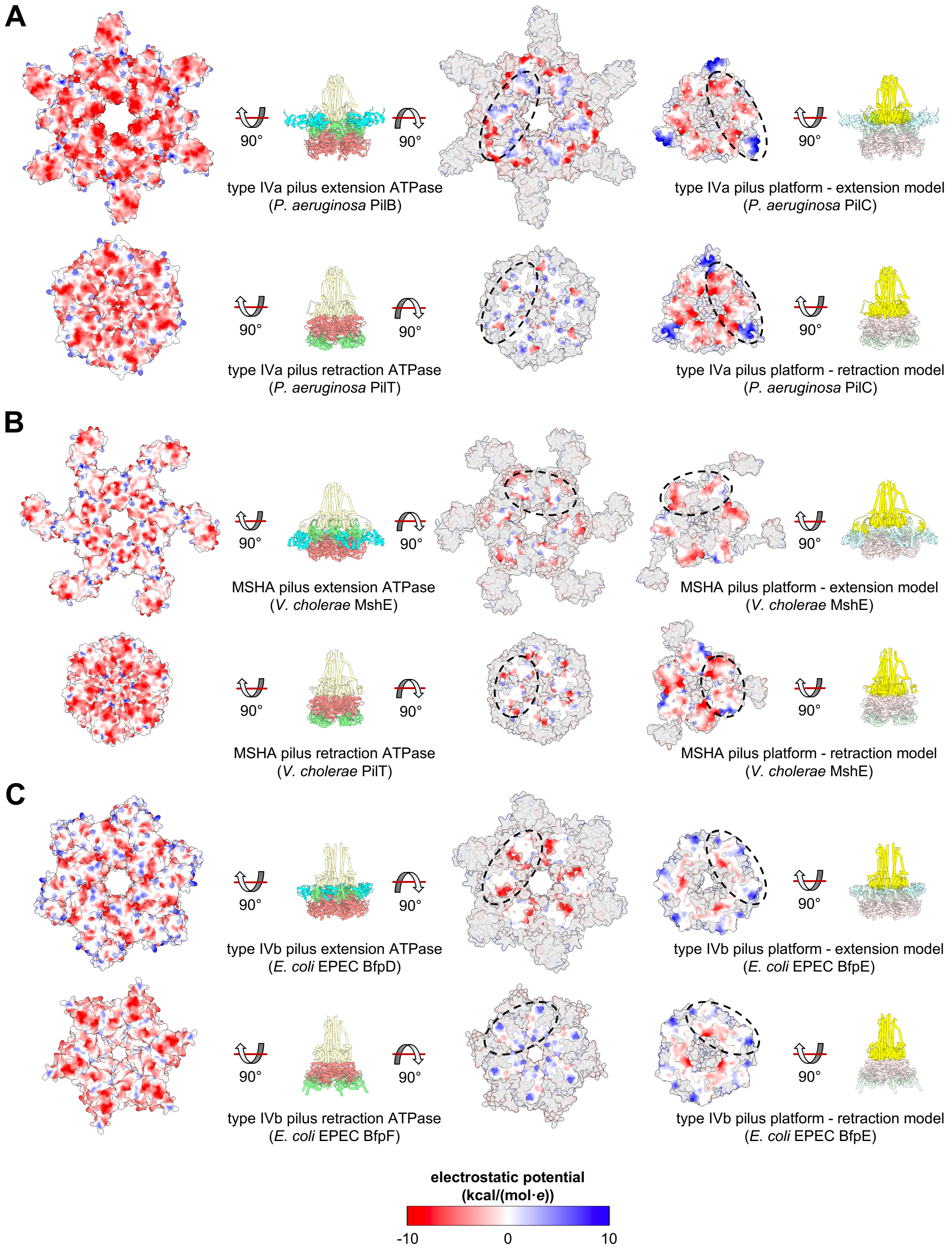


**Fig. S23: Charge distribution on the surface of monofunctional ATPase hexamers and their cognate platform proteins from type IV filament systems suggests that they only adopt one**

**orientation, consistent with their known activity.** (**A-C**) Electrostatic potential was mapped onto the AF3-predicted structures of the extension- and retraction-specific motor ATPases of the type IVa pilus from *Pseudomonas aeruginosa* (panel A), the mannose sensitive hemagglutinin (MSHA) pilus from *Vibrio cholerae* (panel B), and the type IVb bundle forming pilus from enteropathogenic *Escherichia coli* (EPEC; panel C) using the coulombic function in ChimeraX (*34*). Negatively charged regions are coloured red, neutral regions are coloured white, and positively charged regions are coloured blue, as indicated in the legend at the bottom. For clarity on the right-hand side, only the directly interfacing surfaces of the platform and ATPase complexes are coloured, while the remaining surfaces are set to transparent. The approximate interfacing surface of one platform protein subunit with the ATPase hexamer is indicated with the black dashed ovals. On the left-hand side, the entire surface is coloured because no platform interaction interface was predicted by AF3. Protein and protein domain colouring is as indicated in Figure 1.

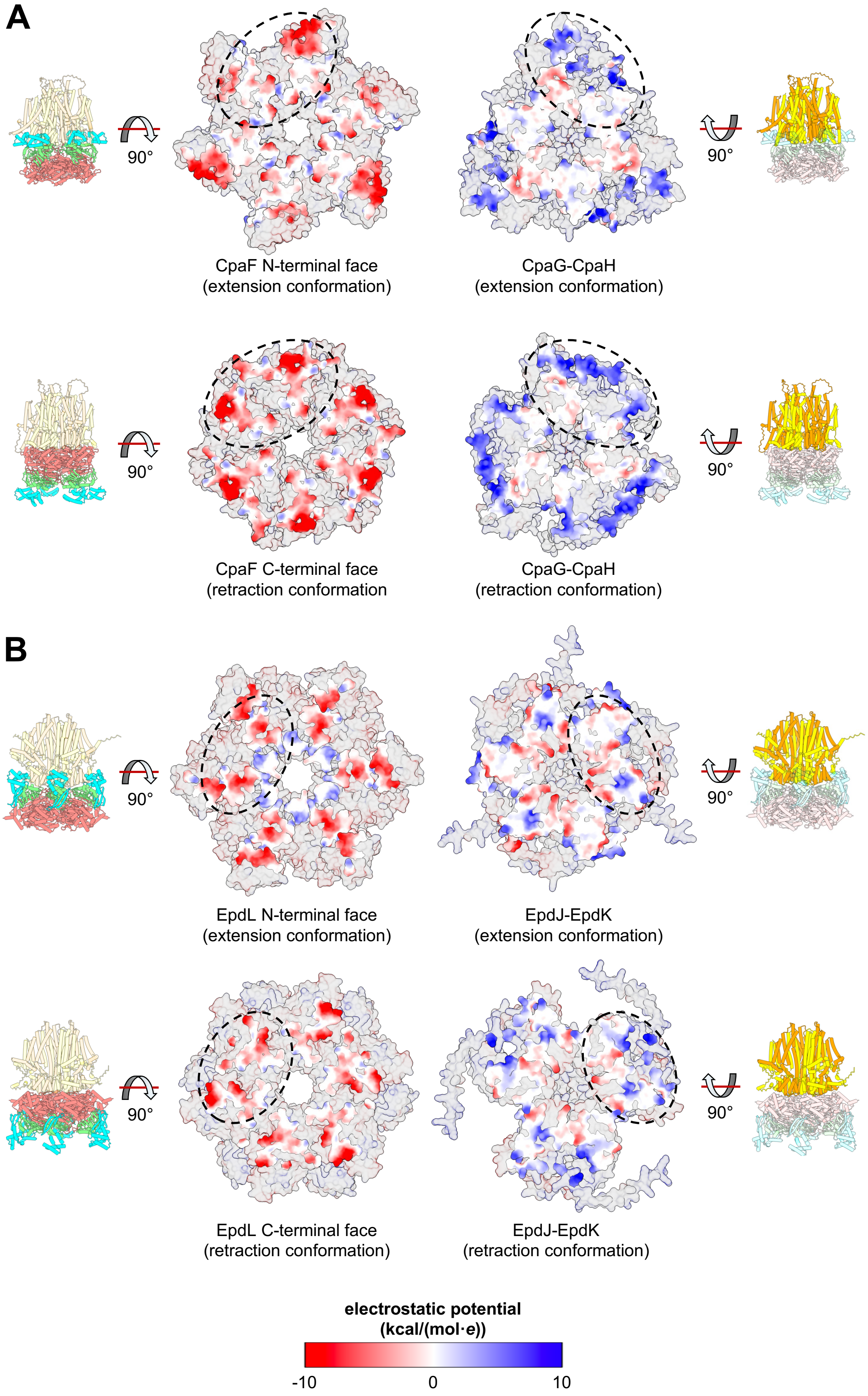
A
90°
90°
CpaF N-terminal face
(extension conformation)
CpaG-CpaH
(extension conformation)
90°
90°
CpaF C-terminal face
(retraction conformation
CpaG-CpaH
(retraction conformation)
B
90°
90°
EpdL N-terminal face
(extension conformation)
EpdJ-EpdK
(extension conformation)
90°
90°
EpdL C-terminal face
(retraction conformation)
EpdJ-EpdK
(retraction conformation)
electrostatic potential
(kcal/(mol·e))
-10
0
10

**Fig. S24: Charge distribution on the surfaces of bifunctional ATPase hexamers and their cognate platform proteins from type IV filament systems is compatible with inversion of the ATPase relative to the platforms.** (**A-B**) Electrostatic potential was mapped onto the AF3-predicted structures of the extension- and retraction-specific motor orientations of the Tad pilus from *Caulobacter crescentus* (panel A) and EppA-dependent (Epd) pilus from *Methanococcus maripaludis* (panel B) using the coulombic function in ChimeraX (*34*). Negatively charged regions are coloured red, neutral regions are coloured white, and positively charged regions are coloured blue, as indicated in the legend at the bottom. For clarity, only the directly interfacing surfaces of the platform and ATPase complexes are coloured, while the remaining surfaces are set to transparent. The approximate interfacing surface of one platform protein subunit with the ATPase hexamer is indicated with the black dashed ovals. Protein and protein domain colouring is as indicated in Figure 1.

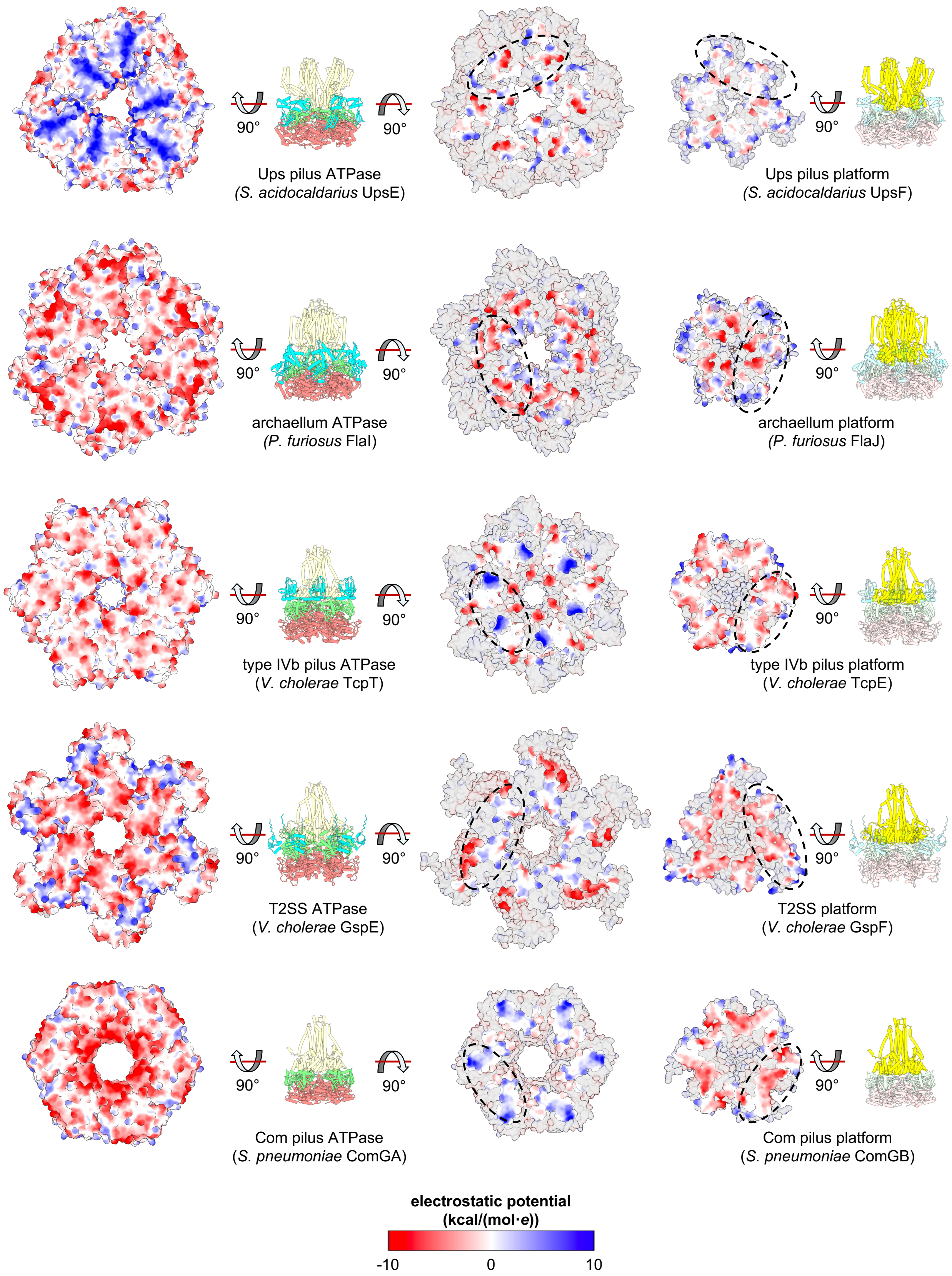
90°
90°
Ups pilus ATPase
(S. acidocaldarius UpsE)
90°
Ups pilus platform
(S. acidocaldarius UpsF)
90°
90°
archaellum ATPase
(P. furiosus FlaI)
90°
archaellum platform
(P. furiosus FlaJ)
90°
90°
type IVb pilus ATPase
(V. cholerae TcpT)
90°
type IVb pilus platform
(V. cholerae TcpE)
90°
90°
T2SS ATPase
(V. cholerae GspE)
90°
T2SS platform
(V. cholerae GspF)
90°
90°
Com pilus ATPase
(S. pneumoniae ComGA)
90°
Com pilus platform
(S. pneumoniae ComGB)
electrostatic potential
(kcal/(mol·e))
-10
0
10

**Fig. S25: Charge distribution on the surfaces of ATPase hexamers and their cognate platform proteins from single-motor type IV filament systems is not compatible with inversion of the ATPase relative to the platforms.** Electrostatic potential was mapped onto the AF3-predicted structures of the motors of, from top to bottom, the UV-inducible pilus system (Ups) from *Sulfolobus acidocaldarius*, the archaellum from *Pyrococcus furiosus*, the type IVb toxin coregulated pilus (Tcp) from *Vibrio cholerae*, the type II secretion system (T2SS) from *Vibrio cholerae*, and the competence (Com) pilus from *Streptococcus pneumoniae,* using the coulombic function in ChimeraX (*34*). Negatively charged regions are coloured red, neutral regions are coloured white, and positively charged regions are coloured blue, as indicated in the legend at the bottom. For clarity on the right-hand side, only the directly interfacing surfaces of the platform and ATPase complexes are coloured, while the remaining surfaces are set to transparent. The approximate interfacing surface of one platform protein subunit with the ATPase hexamer is indicated with the black dashed ovals. On the left-hand side, the entire surface is coloured because no platform interaction interface was predicted by AF3. Protein and protein domain colouring is as indicated in Figure 1.

| Species | Taxonomic group | Ratio ext/ret (full-length) | pTM/ipTM (full-length) | Ratio ext/ret (-N1D) | pTM/ipTM (-N1D) |
|---|---|---|---|---|---|
| | | *Motors predicted to adopt extension and retraction conformations* | | | |
| *Agrobacterium fabrum* | Alphaproteobacteria | 5/0 | ext: 0.66/0.65 | 0/5 | ret: 0.73/0.72 |
| *Caulobacter crescentus** | Alphaproteobacteria | 1/4 | ext: 0.68/0.67, ret: 0.69/0.68 | 0/5 | ret: 0.73/0.72 |
| *Hyphomonas neptunium* | Alphaproteobacteria | 3/2 | ext: 0.59/0.57, ret: 0.69/0.67 | 0/5 | ret: 0.72/0.71 |
| *Sinorhizobium meliloti* | Alphaproteobacteria | 3/2 | ext: 0.67/0.66, ret: 0.67/0.67 | 0/5 | ret: 0.70/0.71 |
| *Sphingobium indicum* | Alphaproteobacteria | 1/4 | ext: 0.71/0.70, ret: 0.67/0.65 | 0/5 | ret: 0.67/0.66 |
| *Basilea psittacipulmonis* | Betaproteobacteria | 5/0 | ext: 0.68/0.67 | 2/3 | ext: 0.63/0.61, ret: 0.68/0.67 |
| *Delftia sp. Cs1-4* | Betaproteobacteria | 4/1 | ext: 0.63/0.62, ret: 0.66/0.64 | 0/5 | ret: 0.69/0.67 |
| *Paraburkholderia phytofirmans* | Betaproteobacteria | 5/0 | ext: 0.69/0.68 | 0/5 | ret: 0.76/0.75 |
| *Ralstonia pseudosolanacearum** | Betaproteobacteria | 5/0 | ext: 0.71/0.70 | 0/5 | ret: 0.68/0.66 |
| *Aggregatibacter actinomycetemcomitans** | Gammaproteobacteria | 5/0 | ext: 0.70/0.69 | 2/3 | ext: 0.66/0.64, ret: 0.71/0.70 |
| *Aliivibrio fischeri* | Gammaproteobacteria | 5/0 | ext: 0.69/0.69 | 3/2 | ext: 0.67/0.66, ret: 0.72/0.71 |
| *Pseudomonas chlororaphis* | Gammaproteobacteria | 4/1 | ext: 0.70/0.69, ret: 0.64/0.63 | 1/4 | ext: 0.63/0.61, ret: 0.67/0.65 |
| *Stenotrophomonas maltophilia* | Gammaproteobacteria | 5/0 | ext: 0.65/0.63 | 1/4 | ext: 0.58/0.55, ret: 0.69/0.68 |
| *Yersinia pestis* | Gammaproteobacteria | 5/0 | ext: 0.74/0.74 | 3/2 | ext: 0.72/0.71, ret: 0.74/0.74 |
| *Arthrobacter sp. FB24** | Actinomycetota | 1/4 | ext: 0.61/0.59, ret: 0.60/0.58 | 0/5 | ret: 0.63/0.61 |
| *Micrococcus luteus* | Actinomycetota | 5/0 | ext: 0.66/0.65 | 4/1 | ext: 0.67/0.65, ret: 0.67/0.65 |
| *Nocardioides sp. JS614* | Actinomycetota | 5/0 | ext: 0.68/0.66 | 2/3 | ext: 0.63/0.66, ret: 0.67/0.65 |
| *Bacillus sp. 1NLA3E** | Bacillota | 0/5 | ret: 0.64/0.62 | 0/5 | ret: 0.63/0.62 |
| *Chlorobaculum tepidum** | Chlorobiota | 5/0 | ext: 0.69/0.68 | 0/5 | ret: 0.74/0.74 |
| *Chloroflexus aurantiacus** | Chloroflexota | 1/4 | ext: 0.65/0.63, ret: 0.61/0.59 | 0/5 | ret: 0.63/0.62 |
| *Anaeromyxobacter dehalogenans** | Myxococcota | 3/2 | ext: 0.67/0.66, ret: 0.67/0.65 | 0/5 | ret: 0.70/0.69 |
| *Planctopirus limnophila** | Planctomycetota | 2/3 | ext: 0.69/0.68, ret: 0.67/0.66 | 0/5 | ret: 0.71/0.70 |
| *Rubinisphaera brasiliensis* | Planctomycetota | 5/0 | ext: 0.67/0.66 | 0/5 | ret: 0.65/0.64 |
| *Aminobacterium colombiense** | Synergistia | 1/4 | ext: 0.59/0.57, ret: 0.64/0.63 | 0/5 | ret: 0.65/0.64 |
| | | *Motors only predicted to adopt the extension conformation* | | | |
| *Pseudomonas aeruginosa** | Gammaproteobacteria | 5/0 | ext: 0.70/0.69 | 5/0 | ext: 0.75/0.74 |
| *Vibrio natriegens** | Gammaproteobacteria | 5/0 | ext: 0.71/0.70 | 5/0 | ext: 0.66/0.65 |
| *Acidothermus cellulolyticus** | Actinomycetota | 5/0 | ext: 0.70/0.70 | 5/0 | ext: 0.72/0.72 |
| *Arcanobacterium haemolyticum* | Actinomycetota | 5/0 | ext: 0.67/0.64 | 5/0 | ext: 0.65/0.63 |
| *Corynebacterium diphtheriae** | Actinomycetota | 5/0 | ext: 0.70/0.69 | 5/0 | ext: 0.71/0.70 |
| *Corynebacterium glutamicum* | Actinomycetota | 5/0 | ext: 0.66/0.65 | 5/0 | ext: 0.65/0.64 |
| *Kytococcus sedentarius** | Actinomycetota | 5/0 | ext: 0.72/0.71 | 5/0 | ext: 0.71/0.70 |
| *Mycobacterium avium** | Actinomycetota | 5/0 | ext: 0.75/0.75 | 5/0 | ext: 0.70/0.69 |
| *Streptomyces coelicolor** | Actinomycetota | 5/0 | ext: 0.63/0.62 | 5/0 | ext: 0.61/0.59 |
| *Acetivibrio clariflavus** | Bacillota | 5/0 | ext: 0.67/0.66 | 5/0 | ext: 0.68/0.66 |
| *Novibacillus thermophilus** | Bacillota | 5/0 | ext: 0.67/0.66 | 5/0 | ext: 0.65/0.63 |
| *Paenibacillus polymyxa* | Bacillota | 5/0 | ext: 0.66/0.64 | 5/0 | ext: 0.65/0.63 |
| *Thermincola potens** | Bacillota | 5/0 | ext: 0.61/0.59 | 5/0 | ext: 0.57/0.55 |

**Table S1: Summary of Tad pilus motor subcomplex structural predictions performed by AF3.** * = CpaF orthologs selected for multiple sequence alignment; ext, extension; ret, retraction; -N1D, N1 domain removed from CpaF sequence for prediction; pTM, predicted template modelling score; ipTM, interface predicted template modelling score. pTM and ipTM scores are shown for the top-ranking extension and retraction models only. Protein accession information can be found in Table S6.

| Species | Taxonomic group | CpaG N-term length | CpaH N-term length | CpaF region II + III length | CpaF region II + III %ID (%sim) to *Cc* |
|---|---|---|---|---|---|
| | *Motors predicted to adopt extension and retraction conformations* | | | | |
| *Agrobacterium fabrum* | Alphaproteobacteria | 128 | 147 | 68 | 66 (82) |
| *Caulobacter crescentus** | Alphaproteobacteria | 119 | 180 | 65 | --- |
| *Hyphomonas neptunium* | Alphaproteobacteria | 121 | 129 | 65 | 34 (51) |
| *Sinorhizobium meliloti* | Alphaproteobacteria | 127 | 141 | 70 | 72 (85) |
| *Sphingobium indicum* | Alphaproteobacteria | 92 | 105 | 53 | 32 (46) |
| *Basilea psittacipulmonis* | Betaproteobacteria | 95 | 101 | 69 | 32 (54) |
| *Delftia* sp. Cs1-4 | Betaproteobacteria | 116 | 147 | 61 | 21 (45) |
| *Paraburkholderia phytofirmans* | Betaproteobacteria | 117 | 132 | 62 | 45 (60) |
| *Ralstonia pseudosolanacearum** | Betaproteobacteria | 120 | 125 | 67 | 33 (56) |
| *Aggregatibacter actinomycetemcomitans** | Gammaproteobacteria | 92 | 107 | 67 | 32 (55) |
| *Aliivibrio fischeri* | Gammaproteobacteria | 102 | 110 | 71 | 32 (62) |
| *Pseudomonas chlororaphis* | Gammaproteobacteria | 123 | 138 | 66 | 31 (52) |
| *Stenotrophomonas maltophilia* | Gammaproteobacteria | 114 | 125 | 60 | 25 (46) |
| *Yersinia pestis* | Gammaproteobacteria | 88 | 94 | 68 | 25 (53) |
| *Arthrobacter* sp. FB24* | Actinomycetota | 99 | 114 | 70 | 37 (51) |
| *Micrococcus luteus* | Actinomycetota | 75 | 135 | 56 | 18 (28) |
| *Nocardioides* sp. JS614 | Actinomycetota | 74 | 130 | 56 | 29 (38) |
| *Bacillus* sp. 1NLA3E* | Bacillota | 114 | 130 | 63 | 37 (54) |
| *Chlorobaculum tepidum** | Chlorobiota | 117 | 128 | 65 | 39 (64) |
| *Chloroflexus aurantiacus** | Chloroflexota | 119 | 124 | 67 | 43 (60) |
| *Anaeromyxobacter dehalogenans** | Myxococcota | 114 | 120 | 67 | 38 (53) |
| *Planctopirus limnophila** | Planctomycetota | 116 | 128 | 67 | 34 (47) |
| *Rubinisphaera brasiliensis* | Planctomycetota | 117 | 135 | 78 | 26 (41) |
| *Aminobacterium colombiense** | Synergistia | 104 | 118 | 65 | 31 (49) |
| | **Average** | **108** | **127** | **65** | **35 (54)** |
| | *Motors only predicted to adopt the extension conformation* | | | | |
| *Pseudomonas aeruginosa** | Gammaproteobacteria | 90 | 122 | 43 | 17 (26) |
| *Vibrio natriegens** | Gammaproteobacteria | 93 | 125 | 63 | 20 (49) |
| *Acidothermus cellulolyticus** | Actinomycetota | 15 | 0 | 45 | 20 (29) |
| *Arcanobacterium haemolyticum* | Actinomycetota | 52 | 114 | 64 | 24 (40) |
| *Corynebacterium diphtheriae** | Actinomycetota | 44 | 0 | 20 | 10 (19) |
| *Corynebacterium glutamicum* | Actinomycetota | 53 | 0 | 31 | 12 (20) |
| *Kytococcus sedentarius** | Actinomycetota | 59 | 0 | 42 | 18 (28) |
| *Mycobacterium avium** | Actinomycetota | 60 | 0 | 38 | 16 (24) |
| *Streptomyces coelicolor** | Actinomycetota | 120 | 111 | 76 | 16 (23) |
| *Acetivibrio clariflavus** | Bacillota | 52 | 114 | 56 | 23 (42) |
| *Novibacillus thermophilus** | Bacillota | 96 | 87 | 67 | 32 (41) |
| *Paenibacillus polymyxa* | Bacillota | 115 | 52 | 65 | 20 (38) |
| *Thermincola potens** | Bacillota | 110 | 136 | 66 | 25 (40) |
| | **Average** | **74** | **66** | **52** | **19 (32)** |

**Table S2: Length and conservation of CpaG and CpaH N-terminal regions from single- and dual-orientation Tad pilus motor orthologs.** * = selected as a representative for multiple sequence alignments; CpaG/CpaH N-term length ; length of the variable N-terminal region adjacent to the conserved T2SSF domain of each ortholog; CpaF region II + III length, the length from β-strand 14 to the C-terminus of each ortholog (see Figure S18 for examples); CpaF region II + III %ID (%sim) to *Cc*, percent sequence identity and percent sequence similarity of the region encompassed by β-strand 14 and the C-terminus of each ortholog compared to the same region in the *C. crescentus* ortholog. Protein accession information can be found in Table S6.

| Single motor TFF family | System examined | Ratio ext/ret (full-length) | pTM/ipTM (full-length) | Ratio ext/ret (-N1D) | pTM/ipTM (-N1D) |
|---|---|---|---|---|---|
| type IVb pilus | *V. cholerae* toxin coregulated pilus* | 5/0 | ext: 0.63/0.61 | 5/0 | ext: 0.65/0.62 |
| | *E. coli* longus pilus | 5/0 | ext: 0.66/0.64 | 5/0 | ext: 0.66/0.64 |
| | *C. rodentium* CFC pilus | 5/0 | ext: 0.67/0.65 | 5/0 | ext: 0.69/0.67 |
| | *S. enterica* serovar Typhi pilus | 5/0 | ext: 0.68/0.66 | 5/0 | ext: 0.69/0.66 |
| | R64 plasmid thin pilus | 5/0 | ext: 0.66/0.64 | 5/0 | ext: 0.67/0.65 |
| Gram positive Com pilus | *S. pneumoniae* Com* | 5/0 | ext: 0.73/0.71 | n/a | n/a |
| | *S. aureus* Com | 5/0 | ext: 0.65/0.63 | n/a | n/a |
| | *B. subtilis* Com | 5/0 | ext: 0.60/0.56 | n/a | n/a |
| type II secretion system | *V. cholerae* Gsp* | 5/0 | ext: 0.51/0.49 | 5/0 | ext: 0.53/0.49 |
| | *P. aeruginosa* Xcp | 5/0 | ext: 0.53/0.51 | 5/0 | ext: 0.68/0.70 |
| | *K. pneumoniae* Pul | 5/0 | ext: 0.50/0.48 | 5/0 | ext: 0.51/0.48 |
| | *D. dadantii* Out | 5/0 | ext: 0.53/0.51 | 5/0 | ext: 0.52/0.49 |
| | *B. pseudomallei* Gsp | 5/0 | ext: 0.58/0.55 | 5/0 | ext: 0.65/0.62 |
| archaeal T4P | *S. acidocaldarius* UV-inducible pilus* | 5/0 | ext: 0.70/0.68 | 5/0 | ext: 0.70/0.68 |
| | *S. acidocaldarius* adhesion pilus | 5/0 | ext: 0.63/0.60 | 5/0 | ext: 0.57/0.53 |
| | *M. maripaludis* Epd pilus* | 1/4 | ext: 0.65/0.64, ret: 0.67/0.66 | 4/1 | ext: 0.64/0.62, ret: 0.67/0.65 |
| | *M. aeolicus* Epd pilus | 4/1 | ext: 0.65/0.64, ret: 0.63/0.62 | 0/5 | ret: 0.67/0.65 |
| | *M. infernus* Epd pilus | 4/1 | ext: 0.66/0.64, ret: 0.62/0.60 | 2/3 | ext: 0.65/0.64, ret: 0.65/0.64 |
| archaellum | *P. furiosus* archaellum* | 5/0 | ext: 0.77/0.75 | 5/0 | ext: 0.75/0.72 |
| | *S. acidocaldarius* archaellum | 5/0 | ext: 0.78/0.76 | 5/0 | ext: 0.77/0.75 |
| | *M. voltae* archaellum | 5/0 | ext: 0.73/0.71 | 5/0 | ext: 0.68/0.64 |
| | *H. volcanii* archaellum | 5/0 | ext: 0.69/0.66 | 5/0 | ext: 0.64/0.60 |

**Table S3: Summary of structural predictions of select type IV filament motor subcomplexes performed by AF3.** * = selected as representative example for surface residue conservation and electrostatics analysis; ext, extension; ret, retraction; -N1D, N1 domain removed from the ATPase sequence for prediction (Com pilus ATPases do not have a N1D); pTM, predicted template modelling score; ipTM, interface predicted template modelling score. pTM and ipTM scores are shown for the top extension and retraction model only. Protein accession information can be found in Table S6.

| Type IV filament class | Representative system | Retraction motor? | AF3 prediction | ATPase surface residue conservation | ATPase-platform surface electrostatics | Motor class |
|---|---|---|---|---|---|---|
| type IVa pilus | *P. aeruginosa* | Y | PilB – extension orientation only | N-term – conserved<br>C-term – variable | N-term – match<br>C-term – clash | I |
| | | | PilT – retraction orientation only | N-term – variable<br>C-term – conserved | N-term – clash<br>C-term – neutral | |
| MSHA pilus | *V. cholerae* | Y | MshE – extension orientation only | N-term – conserved<br>C-term – variable | N-term – neutral<br>C-term – clash | I |
| | | | PilT – retraction orientation only | N-term – variable<br>C-term – conserved | N-term – clash<br>C-term – neutral | |
| type IVb pilus | *E. coli* bundle forming pilus (Bfp) | Y | BfpD – extension orientation only | N-term – conserved<br>C-term – variable | N-term – match<br>C-term – neutral | I |
| | | | BfpF – retraction orientation only | N-term – variable<br>C-term – conserved | N-term – neutral<br>C-term – clash | |
| | *V. cholerae* toxin co-regulated pilus (Tcp) | N | TcpE – extension orientation only | N-term – conserved<br>C-term – conserved | N-term – match<br>C-term – clash | II |
| Com pilus | *S. pneumoniae* | N | ComGA – extension orientation only | N-term – conserved<br>C-term – variable | N-term – match<br>C-term – clash | II |
| type II secretion system | *V. cholerae* | N | GspE – extension orientation only | N-term – conserved<br>C-term – variable | N-term – match<br>C-term – clash | II |
| archaellum | *P. furiosus* | N | FlaI – extension orientation only | N-term – conserved<br>C-term – conserved | N-term – match<br>C-term – clash | II |
| archaeal type IV pilus | *S. acidocaldarius* Ups pilus | N | UpsE – extension orientation only | N-term – conserved<br>C-term – variable | N-term – neutral<br>C-term – clash | II |
| | *M. maripaludis* Epd pilus | N | EpdL – motor inversion | N-term – conserved<br>C-term – conserved | N-term – match<br>C-term – match | III |
| Tad (type IVc) pilus | *C. crescentus* | N | CpaF – motor inversion | N-term – conserved<br>C-term – conserved | N-term – match<br>C-term – match | III |

**Table S4: Summary of surface residue conservation and electrostatics analysis for select type IV filament motor subcomplexes.** ATPase surface residue conservation findings are based on results presented in Figures S19 and S22. ATPase-platform surface electrostatics findings are based on results presented in Figures S23-S25. Motor class I - dual monofunctional ATPases, Motor class II - single ATPase incapable of inversion, Motor class III - single bifunctional ATPase capable of inversion. MSHA, mannose-sensitive hemagglutinin; Com, competence; Ups, UV-inducible pilus system; Epd, EppA-dependent.

| Strain | Description | Source |
| --- | --- | --- |
| ***Escherichia coli* strains** | | |
| DH5α | Cloning strain; F⁻*mcrA* Δ(*mrr-hsd*RMS-*mcr*BC) φ80*lacZ*ΔM15 Δ*lac*X74 *rec*A1 *ara*D139 Δ(*ara-leu*)7697 *gal*U *gal*K λ–*rps*L(Str$^{R}$) *end*A1 *nup*G | Invitrogen |
| NEB5α | DH5α derivative, *fhuA2Δ*(*argF-lacZ*)*U169 phoA glnV44 Φ80Δ*(*lacZ*)*M15 gyrA96 recA1 relA1 endA1 thi-1 hsdR17* | New England Biolabs |
| YB10363 | NEB5α pNPTS138::*cpaF*$^{R387A}$ | This study |
| YB10364 | NEB5α pNPTS138::*cpaF*$^{E438A}$ | This study |
| YB10365 | NEB5α pNPTS138::*cpaF*$^{E443A}$ | This study |
| YB10366 | NEB5α pNPTS138::*cpaF*$^{Q450A}$ | This study |
| YB10367 | NEB5α pNPTS138::*cpaF*$^{D451A}$ | This study |
| YB10368 | NEB5α pNPTS138::*cpaF*$^{R477A}$ | This study |
| YB10369 | NEB5α pNPTS138::*cpaF*$^{mCherry}$ | This study |
| YB10226 | NEB5α pJC585$^{-}$ | 40268890 |
| YB10370 | NEB5α pJC585::*cpaF*$^{E438A}$ | This study |
| YB10371 | NEB5α pJC585::*cpaF*$^{E443A}$ | This study |
| YB10372 | NEB5α pJC585::*cpaF*$^{D451A}$ | This study |
| ***Caulobacter crescentus* strains** | | |
| NA1000 | Synchronizable *C. crescentus* lab adapted strain that does not produce a holdfast | 334726 |
| YB8288 | NA1000 *pilA*$^{T36C}$; pili can be labelled with maleimide-conjugated fluorophores | 29074778 |
| YB8446 | NA1000 *pilA*$^{T36C}$ Δ*cpaF*, unmarked, non-polar deletion of the *cpaF* ORF | 40268890 |
| YB10373 | NA1000 *pilA*$^{T36C}$ *cpaF*$^{R387A}$, allelic exchange with plasmid from YB10363 electroporated into YB8288 | This study |
| YB10374 | NA1000 *pilA*$^{T36C}$ *cpaF*$^{E438A}$, allelic exchange with plasmid from YB10364 electroporated into YB8288 | This study |
| YB10375 | NA1000 *pilA*$^{T36C}$ *cpaF*$^{E443A}$, allelic exchange with plasmid from YB10365 electroporated into YB8288 | This study |

| | | |
|---|---|---|
| YB10376 | NA1000 *pilA*$^{T36C}$ *cpaF*$^{Q450A}$, allelic exchange with plasmid from YB10366 electroporated into YB8288 | This study |
| YB10377 | NA1000 *pilA*$^{T36C}$ *cpaF*$^{D451A}$, allelic exchange with plasmid from YB10367 electroporated into YB8288 | This study |
| YB10378 | NA1000 *pilA*$^{T36C}$ *cpaF*$^{R477A}$, allelic exchange with plasmid from YB10368 electroporated into YB8288 | This study |
| YB10379 | NA1000 *pilA*$^{T36C}$ *cpaF*$^{mCherry}$, allelic exchange with plasmid from YB10369 electroporated into YB8288 | This study |
| YB10237 | NA1000 *pilA*$^{T36C}$ pJC585$^{-}$, electroporation of plasmid from YB10226 into YB8288 | 40268890 |
| YB10380 | NA1000 *pilA*$^{T36C}$ *cpaF*$^{R387A}$ pJC585$^{-}$, electroporation of plasmid from YB10226 into YB10373 | This study |
| YB10381 | NA1000 *pilA*$^{T36C}$ *cpaF*$^{E438A}$ pJC585::*cpaF*$^{E438A}$, electroporation of plasmid from YB10370 into YB10374 | This study |
| YB10382 | NA1000 *pilA*$^{T36C}$ *cpaF*$^{E443A}$ pJC585::*cpaF*$^{E443A}$, electroporation of plasmid from YB10371 into YB10375 | This study |
| YB10383 | NA1000 *pilA*$^{T36C}$ *cpaF*$^{Q450A}$ pJC585$^{-}$, electroporation of plasmid from YB10226 into YB10376 | This study |
| YB10384 | NA1000 *pilA*$^{T36C}$ *cpaF*$^{D451A}$ pJC585::*cpaF*$^{D451A}$, electroporation of plasmid from YB10372 into YB10377 | This study |
| YB10385 | NA1000 *pilA*$^{T36C}$ *cpaF*$^{R477A}$ pJC585$^{-}$, electroporation of plasmid from YB10226 into YB10378 | This study |
| **Plasmids** | | |
| pNPTS138 | Litmus 38 derivative, *nptI oriT sacB, Kan*$^{R}$; used for allelic exchange in *C. crescentus* | M.R.K Alley, unpublished |
| pNPTS138:: *cpaF*$^{R387A}$ | pNPTS138 containing a genomic fragment 204 bp upstream of *cpaF* codon 387 to 677 bp downstream of *cpaF* codon 387 at the EcoRV site, with an arginine to alanine mutation in codon 387 (R387A); used to introduce the R387A mutation into the *cpaF* ORF from the NA1000 genome | This study |
| pNPTS138:: *cpaF*$^{E438A}$ | pNPTS138 containing a genomic fragment 357 bp upstream of *cpaF* codon 438 to 523 bp downstream of *cpaF* codon 438 at the EcoRV site, with a glutamate to alanine mutation in codon 438 (E438A); used to introduce the E438A mutation into the *cpaF* ORF from the NA1000 genome | This study |

| | | |
|---|---|---|
| pNPTS138:: *cpaF*$^{E443A}$ | pNPTS138 containing a genomic fragment 372 bp upstream of *cpaF* codon 443 to 508 bp downstream of *cpaF* codon 443 at the EcoRV site, with a glutamate to alanine mutation in codon 443 (E443A); used to introduce the E443A mutation into the *cpaF* ORF from the NA1000 genome | This study |
| pNPTS138:: *cpaF*$^{Q450A}$ | pNPTS138 containing a genomic fragment 393 bp upstream of *cpaF* codon 450 to 487 bp downstream of *cpaF* codon 450 at the EcoRV site, with a glutamine to alanine mutation in codon 450 (Q450A); used to introduce the Q450A mutation into the *cpaF* ORF from the NA1000 genome | This study |
| pNPTS138:: *cpaF*$^{D451A}$ | pNPTS138 containing a genomic fragment 396 bp upstream of *cpaF* codon 451 to 484 bp downstream of *cpaF* codon 451 at the EcoRV site, with an aspartate to alanine mutation in codon 451 (D451A); used to introduce the D451A mutation into the *cpaF* ORF from the NA1000 genome | This study |
| pNPTS138:: *cpaF*$^{R477A}$ | pNPTS138 containing a genomic fragment 474 bp upstream of *cpaF* codon 477 to 406 bp downstream of *cpaF* codon 477 at the EcoRV site, with an arginine to alanine mutation in codon 477 (R477A); used to introduce the R477A mutation into the *cpaF* ORF from the NA1000 genome | This study |
| pRVCHYN-2 | *nptI vanR Pvan-mCherry trfA bla oriT Kan*$^{R}$; mCherry under the control a vanillate-inducible promoter | 17959646 |
| pNPTS138:: *cpaF*$^{mCherry}$ | pNPTS138 containing the mCherry coding sequence from pRVCHYN-2 with methionine 10 mutated to glutamine (M10Q), sandwiched between 484 bp upstream of the *cpaF* stop codon and 503 bp downstream of and including the *cpaF* stop codon, at the EcoRV site; used to fuse mCherry to the C-terminus of the *cpaF* ORF from the NA1000 genome with a GSAGSAAGSGEF linker | This study |
| pJC585$^{-}$ | pJC585 digested with KpnI to remove part of the vector-encoded *rfp* sequence, used as an empty vector control | 40268890 |
| pJC585::*cpaF*$^{E438A}$ | pJC585 containing *cpaF*$^{E438A}$ fused to a synthetic RBS, inserted between the EcoRI and BamHI sites, under the control of a taurine-inducible promoter | This study |
| pJC585::*cpaF*$^{E443A}$ | pJC585 containing *cpaF*$^{E443A}$ fused to a synthetic RBS, inserted between the EcoRI and BamHI sites, under the control of a taurine-inducible promoter | This study |

| | | |
|---|---|---|
| pJC585::*cpaF*$^{D451A}$ | pJC585 containing *cpaF*$^{D451A}$ fused to a synthetic RBS, inserted between the EcoRI and BamHI sites, under the control of a taurine-inducible promoter | This study |

**Primers**

| Name | Sequence* | Description |
|---|---|---|
| *cpaF*$^{C\text{-term mutant}}$ upF | **GCCAAGCTTCTCTGCAGGAT**CACGTGGTGCGCCTGGAAACC | Used to make all pNPTS138 *cpaF* point mutant alleles |
| *cpaF*$^{R387A}$ upR | *TGATCGCCTCAgcCGGGCTGTTG*GCGTGCAGCGTGCCCATCGA | Used to make pNPTS138::*cpaF*$^{R387A}$ |
| *cpaF*$^{R387A}$ downF | *CAACAGCCCGgcTGAGGCGATCA*GCCGGATCGAGAGCATGATCAC | Used to make pNPTS138::*cpaF*$^{R387A}$ |
| *cpaF*$^{C\text{-term mutant}}$ downR | **GCGAATTCGTGGATCCAGAT**GCGATGAATAGGCCAAGCGTCC | Used to make all pNPTS138 *cpaF* point mutant alleles |
| *cpaF*$^{E438A}$ upR | *GCCCACGACCgCGGTGATGTG*GGTGATGCGGCGCGAACCGT | Used to make pNPTS138::*cpaF*$^{E438A}$ |
| *cpaF*$^{E438A}$ downF | *CACATCACCGcGGTCGTGGGC*CTGGAAGGCGACGTGATCGTCA | Used to make pNPTS138::*cpaF*$^{E438A}$ |
| *cpaF*$^{E443A}$ upR | *CACGTCGCCTgCCAGGCCCAC*GACCTCGGTGATGTGGGTGATGC | Used to make pNPTS138::*cpaF*$^{E443A}$ |
| *cpaF*$^{E443A}$ downF | *GTGGGCCTGGcAGGCGACGTG*ATCGTCACCCAGGACCTCTTCGTC | Used to make pNPTS138::*cpaF*$^{E443A}$ |
| *cpaF*$^{Q450A}$ upR | *GAAGAGGTCCgcGGTGACGATC*ACGTCGCCTTCCAGGCCCACGA | Used to make pNPTS138::*cpaF*$^{Q450A}$ |
| *cpaF*$^{Q450A}$ downF | *GATCGTCACCgcGGACCTCTTC*GTCTACGAGATCACCGGCGAGGA | Used to make pNPTS138::*cpaF*$^{Q450A}$ |
| *cpaF*$^{D451A}$ upR | *GACGAAGAGGgCCTGGGTGAC*GATCACGTCGCCTTCCAGGCC | Used to make pNPTS138::*cpaF*$^{D451A}$ |
| *cpaF*$^{D451A}$ downF | *GTCACCCAGGcCCTCTTCGTC*TACGAGATCACCGGCGAGGACG | Used to make pNPTS138::*cpaF*$^{D451A}$ |
| *cpaF*$^{R477A}$ upR | *GAAGCGCGGAgcGGCGATGCC*GGTCGAGCGGTGCTTGCCCACGACCTT | Used to make pNPTS138::*cpaF*$^{R477A}$ |
| *cpaF*$^{R477A}$ downF | *GGCATCGCCgcTCCGCGCTTC*TGGGATCGCGCCCGCTACTACGGG | Used to make pNPTS138::*cpaF*$^{R477A}$ |

| | | |
|---|---|---|
| *cpaF*$^{mCherry}$ upF | **GCCAAGCTTCTCTGCAGGAT**TCTGGTCAAGAACTGTCTGCGGA | Used to make pNPTS138::*cpaF*$^{C-mCherry}$ |
| *cpaF*$^{mCherry}$ upR | *GCCCGAGCCGGCGGCCGAGCCGGCCGAGC*CCTCCGCCGCGTCGAGGGCTT | Used to make pNPTS138::*cpaF*$^{C-mCherry}$ |
| *cpaF*$^{mCherry}$ midF | *GCCGGCTCGGCCGCCGGCTCGGGCGAGTTC*ATGGTGAGCAAGGGCGAGGAGGA | Used to make pNPTS138::*cpaF*$^{C-mCherry}$ |
| *cpaF*$^{mCherry}$ midR | *CCAGGACGAACAGCATGGCCTA*CTTGTACAGCTCGTCCATGCCG | Used to make pNPTS138::*cpaF*$^{C-mCherry}$ |
| *cpaF*$^{mCherry}$ downF | *CGGCATGGACGAGCTGTACAAG*TAGGCCATGCTGTTCGTCCTGG | Used to make pNPTS138::*cpaF*$^{C-mCherry}$ |
| *cpaF*$^{mCherry}$ downR | **GCGAATTCGTGGATCCAGAT**TTGATGCCGCGTACGATGATGTC | Used to make pNPTS138::*cpaF*$^{C-mCherry}$ |
| mCherry$^{M10Q}$ upR | *GATGATGGCCtgGTTATCCTCC*TCGCCCTTGCTCACCATGAACT | Used to make pNPTS138::*cpaF*$^{C-mCherry}$ |
| mCherry$^{M10Q}$ downF | *GGAGGATAACcaGGCCATCATC*AAGGAGTTCATGCGCTTCAAGGTG | Used to make pNPTS138::*cpaF*$^{C-mCherry}$ |
| $P_{tau}$-*cpaF* upF | GTGTG**GAATTC**TTTAAGAAGGAGATATACATATGTTCGGAAAGCGCGACTCGTCAG | Used to make all pJC585 *cpaF* expression plasmids |
| $P_{tau}$-*cpaF*$^{E438A}$ upR | *GCCCACGACCgCGGTGATGTG*GGTGATGCGGCGCGAACCGT | Used to make pJC585::*cpaF*$^{E438A}$ |
| $P_{tau}$-*cpaF*$^{E438A}$ downF | *CACATCACCGcGGTCGTGGGC*CTGGAAGGCGACGTGATCGTCA | Used to make pJC585::*cpaF*$^{E438A}$ |
| $P_{tau}$-*cpaF* downR | GTGTG**GGATCC**CTACTCCGCCGCGTCGAGGG | Used to make all pJC585 *cpaF* expression plasmids |
| $P_{tau}$-*cpaF*$^{E443A}$ upR | *CACGTCGCCTgCCAGGCCCAC*GACCTCGGTGATGTGGGTGATGC | Used to make pJC585::*cpaF*$^{E443A}$ |
| $P_{tau}$-*cpaF*$^{E443A}$ downF | *GTGGGCCTGGcAGGCGACGTG*ATCGTCACCCAGGACCTCTTCGTC | Used to make pJC585::*cpaF*$^{E443A}$ |

| | | |
|---|---|---|
| P$_{tau}$-*cpaF*$^{D451A}$ upR | *GACGAAGAGGgCCTGGGTGAC*GATCACGTCG CCTTCCAGGCC | Used to make pJC585::*cpaF*$^{D451A}$ |
| P$_{tau}$-*cpaF*$^{D451A}$ downF | *GTCACCCAGGcCCTCTTCGTC*TACGAGATCA CCGGCGAGGACG | Used to make pJC585::*cpaF*$^{D451A}$ |

**Table S5: Bacterial strains, plasmids, and primers used in this study.** *Sequences for Gibson assembly into destination plasmids, or restriction enzyme recognition sequences, are bolded; regions of complementarity to the target amplicon are underlined; regions of reverse complementarity to facilitate splicing are italicized; nucleotides that differ from the coding sequence (to introduce point mutations) are indicated in lower case.

| TFF system | Organism | ATPase | | Platform | |
|---|---|---|---|---|---|
| | | Protein | Accession | Protein | Accession |
| Type IVa pilus | *Pseudomonas aeruginosa* PAO1 | PilB | WP_003112841.1 | PilC | WP_209243908.1 |
| | | PilT | WP_003084552.1 | | |
| MSHA pilus | *Vibrio cholerae* O1 biovar El Tor str. N16961 | MshE | WP_001122154.1 | MshG | WP_000190444.1 |
| | | PilT | WP_000350195.1 | | |
| Type IVb pilus | *Citrobacter rodentium* DSM 16636 | CfcH | WP_012908545.1 | CfcI | WP_012908546.1 |
| | *Escherichia coli* ETEC 1392/75 | LngH | WP_000526869.1 | LngI | WP_001393141.1 |
| | *Escherichia coli* O127:H6 str. E2348/69 | BfpD | WP_012477166.1 | BfpE | WP_000660204.1 |
| | | BfpF | WP_001183608.1 | | |
| | *Salmonella enterica* subsp. *enterica* serovar *Typhi* str. Ty2 | PilQ | WP_001289222.1 | PilR | WP_000074784.1 |
| | *Salmonella enterica* subsp. *enterica* ATCC 43971 plasmid R64 | PilQ | WP_000362202.1 | PilR | WP_001208805.1 |
| | *Vibrio cholerae* O1 biovar El Tor str. N16961 | TcpT | WP_000020697.1 | TcpE | WP_000691946.1 |
| Tad pilus | *Acetivibrio clariflavus* DSM 19732 | CpaF | WP_014255951.1 | CpaG | WP_027621775.1 |
| | | | | CpaH | WP_014255949.1 |
| | *Acidothermus cellulolyticus* 11B | CpaF | WP_011720817.1 | CpaG | WP_049751509.1 |
| | | | | CpaH | WP_011720815.1 |
| | *Aggregatibacter actinomycetemcomitans* D11S-1 | TadA | WP_005546820.1 | TadB | WP_005546823.1 |
| | | | | TadC | WP_005546825.1 |
| | *Agrobacterium fabrum* str. C58 | CtpG | WP_010970737.1 | CtpH | WP_010970736.1 |
| | | | | CtpI | WP_006310061.1 |
| | *Aliivibrio fischeri* ES114 | CpaF | WP_011261282.1 | CpaG | WP_011261283.1 |
| | | | | CpaH | WP_011261284.1 |
| | *Aminobacterium colombiense* DSM 12261 | CpaF | WP_013048584.1 | CpaG | WP_013048583.1 |
| | | | | CpaH | WP_013048582.1 |
| | *Anaeromyxobacter dehalogenans* 2CP-C | CpaF | WP_011421877.1 | CpaG | WP_011421876.1 |
| | | | | CpaH | WP_011421875.1 |
| | *Arcanobacterium haemolyticum* DSM 20595 | CpaF | WP_013170425.1 | CpaG | WP_013170424.1 |
| | | | | CpaH | WP_013170423.1 |
| | *Arthrobacter* sp. FB24 | CpaF | WP_011692763.1 | CpaG | WP_232223525.1 |
| | | | | CpaH | WP_011692765.1 |
| | *Bacillus* sp. 1NLA3E | CpaF | WP_015595996.1 | CpaG | WP_051120182.1 |
| | | | | CpaH | WP_015595994.1 |

| | | | | |
|---|---|---|---|---|
| *Basilea psittacipulmonis* DSM 24701 | CpaF | WP_038498858.1 | CpaG | WP_038498861.1 |
| | | | CpaH | WP_038501428.1 |
| *Caulobacter crescentus* NA1000 | CpaF | WP_010920779.1 | CpaG | WP_010920778.1 |
| | | | CpaH | WP_010920777.1 |
| *Chlorobaculum tepidum* TLS | CpaF | WP_010932125.1 | CpaG | WP_164927209.1 |
| | | | CpaH | WP_010932127.1 |
| *Chloroflexus aurantiacus* J-10-fl | CpaF | WP_012256416.1 | CpaG | WP_012256415.1 |
| | | | CpaH | WP_012256414.1 |
| *Corynebacterium diphtheriae* NCTC 13129 | CpaF | WP_010934203.1 | CpaG | WP_010934204.1 |
| | | | CpaH | WP_010934205.1 |
| *Corynebacterium glutamicum* R | CpaF | WP_011896610.1 | CpaG | WP_011896611.1 |
| | | | CpaH | WP_011896612.1 |
| *Delftia* sp. Cs1-4 | CpaF | WP_012203993.1 | CpaG | WP_013803755.1 |
| | | | CpaH | WP_013803756.1 |
| *Hyphomonas neptunium* ATCC 15444 | CpaF | WP_011646172.1 | CpaG | WP_233352007.1 |
| | | | CpaH | WP_011646174.1 |
| *Kytococcus sedentarius* DSM 20547 | CpaF | WP_015780419.1 | CpaG | WP_015780418.1 |
| | | | CpaH | WP_015780417.1 |
| *Micrococcus luteus* NCTC 2665 | CpaF | WP_010079081.1 | CpaG | WP_010079080.1 |
| | | | CpaH | WP_010079079.1 |
| *Mycobacterium avium* subsp. *paratuberculosis* K-10 | CpaF | WP_003873281.1 | CpaG | WP_003873280.1 |
| | | | CpaH | WP_003873279.1 |
| *Nocardioides* sp. JS614 | CpaF | WP_011754714.1 | CpaG | WP_011754715.1 |
| | | | CpaH | WP_011754716.1 |
| *Novibacillus thermophilus* SG-1 | CpaF | WP_077719531.1 | CpaG | WP_077719532.1 |
| | | | CpaH | WP_077719533.1 |
| *Paenibacillus polymyxa* E681 | CpaF | WP_013309275.1 | CpaG | WP_043882104.1 |
| | | | CpaH | WP_080942820.1 |
| *Paraburkholderia phytofirmans* OLGA172 | CpaF | WP_063498652.1 | CpaG | WP_063498651.1 |
| | | | CpaH | WP_063498650.1 |
| *Planctopirus limnophila* DSM 3776 | CpaF | WP_013110179.1 | CpaG | WP_013110178.1 |
| | | | CpaH | WP_013110177.1 |
| *Pseudomonas aeruginosa* PAO1 | CpaF | WP_003103953.1 | CpaG | WP_003114981.1 |
| | | | CpaH | WP_003103955.1 |
| *Pseudomonas chlororaphis* YL-1 | CpaF | WP_009050437.1 | CpaG | WP_009050436.1 |
| | | | CpaH | WP_009050435.1 |

| | | | | | |
|---|---|---|---|---|---|
| | *Ralstonia pseudosolanacearum* GMI1000 | CpaF | WP_011000609.1 | CpaG | WP_011000608.1 |
| | | | | CpaH | WP_011000607.1 |
| | *Rubinisphaera brasiliensis* DSM 5305 | CpaF | WP_013629583.1 | CpaG | WP_013629584.1 |
| | | | | CpaH | WP_013629585.1 |
| | *Sinorhizobium meliloti* 1021 | CpaF2 | WP_010967819.1 | CpaG | WP_010967818.1 |
| | | | | CpaH | WP_010967817.1 |
| | *Sphingobium indicum* B90A | CpaF | WP_007684462.1 | CpaG | WP_073507211.1 |
| | | | | CpaH | WP_007684463.1 |
| | *Stenotrophomonas maltophilia* R551-3 | CpaF | WP_012511344.1 | CpaG | WP_012511345.1 |
| | | | | CpaH | WP_012511346.1 |
| | *Streptomyces coelicolor* A3(2) | CpaF | WP_003973968.1 | CpaG | WP_011030000.1 |
| | | | | CpaH | WP_011030001.1 |
| | *Thermincola potens* JR | CpaF | WP_013119292.1 | CpaG | WP_013119293.1 |
| | | | | CpaH | WP_013119294.1 |
| | *Vibrio natriegens* ATCC 14048 | CpaF | WP_020336263.1 | CpaG | WP_020336264.1 |
| | | | | CpaH | WP_020336265.1 |
| | *Yersinia pestis* KIM10+ | CpaF | WP_002213785.1 | CpaG | WP_002212164.1 |
| | | | | CpaH | WP_002212165.1 |
| Com pilus | *Bacillus subtilis* subsp. *subtilis* str. 168 | ComGA | WP_004399124.1 | ComGB | WP_003246116.1 |
| | *Staphylococcus aureus* subsp. *aureus* str. Newman | ComGA | WP_000697228.1 | ComGB | WP_000775708.1 |
| | *Streptococcus pneumoniae* TIGR4 | ComGA | WP_000249564.1 | ComGB | WP_074196785.1 |
| T2SS | *Burkholderia pseudomallei* 1026b | GspE | WP_004553718.1 | GspF | WP_004196821.1 |
| | *Dickeya dadantii* 3937 | OutE | WP_013318809.1 | OutF | WP_013318808.1 |
| | *Klebsiella pneumoniae* | PulE | WP_049088228.1 | PulF | AAA25128.1 |
| | *Pseudomonas aeruginosa* PAO1 | XcpR | WP_003091377.1 | XcpS | WP_003091376.1 |
| | *Vibrio cholerae* O1 biovar El Tor str. N16961 | GspE | WP_000138174.1 | GspF | WP_000718700.1 |
| Archaeal pilus | *Methanocaldococcus infernus* ME | EpdL | WP_013099734.1 | EpdJ | WP_013099736.1 |
| | | | | EpdK | WP_013099735.1 |
| | *Methanococcus aeolicus* Nankai-3 | EpdL | WP_011973863.1 | EpdJ | WP_011973861.1 |
| | | | | EpdK | WP_011973862.1 |
| | *Methanococcus maripaludis* S2 | EpdL | WP_011169984.1 | EpdJ | WP_011169982.1 |
| | | | | EpdK | WP_011169983.1 |

| | | | | | |
|---|---|---|---|---|---|
| | *Sulfolobus acidocaldarius* DSM 639 | AapE | WP_011279104.1 | AapF | WP_011279105.1 |
| | *Sulfolobus acidocaldarius* DSM 639 | UpsE | WP_011278317.1 | UpsF | WP_011278318.1 |
| Archaellum | *Haloferax volcanii* DS2 | FlaI | WP_004043725.1 | FlaJ | WP_004043724.1 |
| | *Methanococcus voltae* C2 | FlaI | WP_013180672.1 | FlaJ | WP_013180671.1 |
| | *Pyrococcus furiosus* DSM 3638 | FlaI | WP_011011446.1 | FlaJ | WP_011011445.1 |
| | *Sulfolobus acidocaldarius* DSM 639 | FlaI | WP_011278022.1 | FlaJ | WP_011278021.1 |

**Table S6: Accession numbers of pilus motor proteins used for AF3 modeling.**

**Movie S1:** Time-lapse of parent (NA1000 *pilA*$^{T36C}$) cells extending and retracting pili after labeling with AF488-mal. Movie corresponds to Figure 2C. The phase channel is shown on the left and the GFP channel is shown on the right. The capture rate is 3 s/frame. Scale bar is 3 μm.

**Movie S2:** Time-lapse of CpaF$^{R387A}$ cells that are unable to retract their pili after labeling with AF488-mal. Movie corresponds to Figure 2D. The phase channel is shown on the left and the GFP channel is shown on the right. The capture rate is 3 s/frame. Scale bar is 3 μm.

**Movie S3:** Time-lapse of parent (NA1000 *pilA*$^{T36C}$) cells that are unable to retract their pili after artificially blocking pilus retraction wth PEG5000-mal and labeling with AF488-mal. Movie corresponds to Figure S7C. The phase channel is shown on the left and the GFP channel is shown on the right. The capture rate is 3 s/frame. Scale bar is 3 μm.

**Movie S4:** Time-lapse of CpaF$^{E438A}$ cells that are unable to retract their pili after labeling with AF488-mal. Movie corresponds to Figure S7D. The phase channel is shown on the left and the GFP channel is shown on the right. The capture rate is 3 s/frame. Scale bar is 3 μm.

**Movie S5:** Time-lapse of CpaF$^{E443A}$ cells that are unable to retract their pili after labeling with AF488-mal. Movie corresponds to Figure S7D. The phase channel is shown on the left and the GFP channel is shown on the right. The capture rate is 3 s/frame. Scale bar is 3 μm.

**Movie S6:** Time-lapse of CpaF$^{Q450A}$ cells that are unable to retract their pili after labeling with AF488-mal. Movie corresponds to Figure S7D. The phase channel is shown on the left and the GFP channel is shown on the right. The capture rate is 3 s/frame. Scale bar is 3 μm.

**Movie S7:** Time-lapse of CpaF$^{D451A}$ cells that are unable to retract their pili after labeling with AF488-mal. Movie corresponds to Figure S7D. The phase channel is shown on the left and the GFP channel is shown on the right. The capture rate is 3 s/frame. Scale bar is 3 μm.

**Movie S8:** Time-lapse of CpaF$^{R477A}$ cells that are unable to retract their pili after labeling with AF488-mal. Movie corresponds to Figure S7D. The phase channel is shown on the left and the GFP channel is shown on the right. The capture rate is 3 s/frame. Scale bar is 3 μm.

**Movie S9:** Time-lapse of CpaF$^{E438A}$ cells that can retract their pili after labeling with AF488-mal. Movie corresponds to Figure S8. The phase channel is shown on the left and the GFP channel is shown on the right. The capture rate is 3 s/frame. Scale bar is 3 μm.

**Movie S10:** Time-lapse of CpaF$^{E443A}$ cells that can retract their pili after labeling with AF488-mal. Movie corresponds to Figure S8. The phase channel is shown on the left and the GFP channel is shown on the right. The capture rate is 3 s/frame. Scale bar is 3 μm.

**Movie S11:** Time-lapse of CpaF$^{Q450A}$ cells that can retract their pili after labeling with AF488-mal. Movie corresponds to Figure S8. The phase channel is shown on the left and the GFP channel is shown on the right. The capture rate is 3 s/frame. Scale bar is 3 μm.

**Movie S12:** Time-lapse of CpaF$^{D451A}$ cells that can retract their pili after labeling with AF488-mal. Movie corresponds to Figure S8. The phase channel is shown on the left and the GFP channel is shown on the right. The capture rate is 3 s/frame. Scale bar is 3 μm.

**Movie S13:** Time-lapse of CpaF$^{R477A}$ cells that can retract their pili after labeling with AF488-mal. Movie corresponds to Figure S8. The phase channel is shown on the left and the GFP channel is shown on the right. The capture rate is 3 s/frame. Scale bar is 3 μm.

**Movie S14:** Time-lapse of parent (NA1000 *pilA*$^{T36C}$) cells that are unable to retract their pili after labeling with AF488-mal. Movie corresponds to Figure S7D. The phase channel is shown on the left and the GFP channel is shown on the right. The capture rate is 3 s/frame. Scale bar is 3 μm.

**Movie S15:** Time-lapse of CpaF$^{mCherry}$ cells that are unable to retract their pili after labeling with AF488-mal. Movie corresponds to Figure 2E. The phase channel is shown on the left and the GFP channel is shown on the right. The capture rate is 3 s/frame. Scale bar is 3 μm.
